# Dissecting the differential role of C-terminal truncations in the regulation of aSyn pathology formation and the biogenesis of Lewy bodies

**DOI:** 10.1101/2024.11.29.625993

**Authors:** Anne-Laure Mahul-Mellier, Melek Firat Altay, Niran Maharjan, Nadine Ait-Bouziad, Anass Chiki, Somanath Jagannath, Galina Limorenko, Salvatore Novello, Jonathan Ricci, Siv Vingill, Richard Wade-Martins, Janice Holton, Catherine Strand, Caroline Haikal, Jia-Yi Li, Romain Hamelin, Marie Croisier, Graham Knott, Georges Mairet-Coello, Laura Weerens, Anne Michel, Patrick Downey, Martin Citron, Hilal A. Lashuel

**Author notes:** These authors contributed equally (Melek Firat Altay, Niran Maharjan, Nadine Ait-Bouziad, Anass Chiki). To whom correspondence should be addressed: Laboratory of Molecular and Chemical Biology of Neurodegeneration, Brain Mind Institute, Ecole Polytechnique Fédérale de Lausanne, 1015 Lausanne.

## Abstract

Alpha-synuclein (aSyn) post-translational modifications (PTMs), particularly phosphorylation at serine 129 and C-terminal truncations, are highly enriched in Lewy bodies (LBs), Lewy neurites, and other types of aSyn pathological aggregates in the brain of patients with Parkinson’s disease (PD) and other synucleinopathies. However, our knowledge about the precise role of PTMs in regulating the different stages of pathology formation, neurodegeneration, and aSyn pathology spreading remains incomplete. In this work, we applied a systematic approach to address this knowledge gap with an emphasis on mapping and elucidating the role of post-fibrillization C-terminal aSyn truncations in regulating the uptake, processing, seeding activity, and formation of LB-like inclusions and maturations in a well-established neuronal seeding model that recapitulates all the stages leading to LB formation and neurodegeneration. Our work shows that C-terminal cleavage of aSyn fibrils at multiple sites is a conserved process that occurs rapidly after and during the formation of intracellular LB-like aSyn inclusions in all neuronal seeding models. Interestingly, blocking the cleavage of internalized fibrils does not influence their seeding activity, whereas inhibiting the enzymes that regulate the cleavage of newly formed fibrils (e.g., calpains 1 and 2) significantly alters the formation of LB-like inclusions. We also show that C-terminal truncations, in combination with other PTMs, play a crucial role in regulating the interactome and remodeling of newly formed aSyn fibrils, including their shortening, lateral association, and packing during LB formation and maturation. Altogether, our results demonstrate that post-fibrillization C-terminal truncations have a differential role at different stages of aSyn aggregation and pathology formation. These insights, combined with the abundance of truncated aSyn species in LBs, have significant implications in understanding aSyn pathological diversity and developing therapeutic strategies targeting the C-terminus of aSyn or its proteolytic processing.

## Introduction

The intracellular accumulation of aggregated forms of alpha-synuclein (aSyn) in neurons and glial cells represents one of the main pathological hallmarks of Parkinson’s disease (PD) and related synucleinopathies, including dementia with Lewy bodies (DLB) and multiple system atrophy (MSA)^1^. PD and DLB are characterized by the presence of Lewy bodies (LB) in neurons, while in MSA, aSyn inclusions are mainly detected in glial cells and are referred to as glial cytoplasmic inclusions (GCIs). Although it is widely believed that aSyn plays a central role in the pathogenesis of synucleinopathies, our understanding of the molecular and cellular processes that trigger and govern the misfolding, fibrillization, inclusion formation, and spread of aSyn in the brain remains incomplete.

Several post-translational modifications (PTMs), including phosphorylation^2,3^, C-terminal (C-ter) truncation^2,4–7^, ubiquitination^8^ and nitration^9^, have been consistently associated with aggregates and LBs found in postmortem brains of patients with PD. This suggests that these modifications represent either markers of Lewy pathologies or key events that regulate the initiation of aSyn misfolding, aggregation, and/or inclusion formation. The identification of PTMs that enhance or interfere with any of these processes could open new avenues for targeting aSyn and developing strategies to prevent or delay pathology formation and disease progression.

Several studies by our group and others^10–18^ have investigated the role of PD-associated PTMs in regulating aSyn aggregation. Collectively, these studies have shown that most pathology-associated PTMs either inhibit or do not affect aSyn fibril formation *in vitro* and in different synucleinopathy models, with the only notable exception being C-terminal truncations^2,4,19–22^. Although several studies have shown that the C-terminal cleavage of aSyn occurs under both physiological^23,24^ and pathogenic^4,6,22,25^ conditions, studies in cell-free systems^26–32^ and cellular and animal models^25,29–39^ of synucleinopathies have consistently shown that C-terminally truncated aSyn monomeric variants aggregate much faster than the full-length protein and accelerate aSyn aggregation and pathology formation. Furthermore, C-terminally truncated aSyn aggregates retain the ability to seed the aggregation of full-length aSyn^30^.

The development of antibodies that specifically recognized the C-terminally truncated fragments enabled further confirmation of their presence by IHC in the LBs^2,35,40–43^ and GCIs^2^. Consistent with previous studies^2,4,40,44–46^, confocal imaging^40^ and super-resolution microscopy^45^ combined with the use of these C-terminal antibodies demonstrated that the C-terminally truncated species (1-119 and 1-122) were not randomly distributed in LBs, pale bodies (PBs), and Lewy neurites (LNs), but systematically arranged in the inner core of these pathological inclusions. In both studies, full-length (FL) and pS129-positive aSyn were found in the outer layers. These observations suggest that the FL and truncated aSyn distribution inside the inclusions is highly orchestrated. Furthermore, studies^47–49^ using monoclonal antibodies targeting neo-epitopes of C-terminally truncated aSyn, combined with multiplex immunoassays^47,48^ or real-time quaking-induced conversion (RT-QuIC) assay^49^, have uncovered significant variations in αSyn C-terminal truncations and pS129 phosphorylation patterns that are specific to the disease, brain region, and cell type within synucleinopathies such as PD, DLB, Alzheimer disease with amygdala Lewy bodies (AD/ALB), and MSA. These studies demonstrated that C-terminally truncated aSyn species are more prevalent than previously thought in the brain and revealed unique biochemical signatures for each synucleinopathy, suggesting that the heterogeneity in aSyn pathology may contribute to the distinct clinical manifestations and progression observed in these diseases. Moreover, we recently demonstrated that astrocytic aSyn inclusions in PD and other synucleinopathies are made of a nonfibrillar form of aSyn that is nitrated or phosphorylated at Y39 and cleaved at both N-terminal (N-ter) and C-terminal domains of the protein (first 30 a.a. and last 30 a.a. are absent)^50,51^. This is the first report of aSyn accumulation that is made purely of C-terminally truncated forms of aSyn, i.e. no full-length aSyn species could be detected in aSyn astrocytic inclusions. Altogether, these data reinforce the potential role of truncation as a key driver in the formation of aSyn aggregates and biogenesis of LBs and that they may have cell-specific functions in PD and synucleinopathies^22,52^. However, the contributions of C-terminal truncation during the various stages of aSyn pathology formation and their role in the evolution and maturation of LBs have not been systematically investigated.

The majority of previous studies on aSyn truncations have focused primarily on exploring how C-terminal truncations and other PTMs influence the kinetics and aggregation properties of monomeric aSyn. However, increasing evidence suggests that several PTMs, including phosphorylation (pS129)^53–58^ and C-terminal cleavage^56–58^, could also occur after aSyn fibrillization in mammalian cells lines^53,56^, neurons in culture^54,57,58^ and *in vivo*^55,57^. Therefore, post-fibrillization PTMs, such as C-terminal truncations, could also play important roles in regulating secondary aggregation processes, including secondary nucleation, lateral association of aSyn aggregates, LB formation and maturation.

To decipher the roles aSyn C-terminal cleavage at different stages of LB formation, we performed systematic studies to elucidate the nature of post-aggregation PTMs that take place upon the 1) uptake of preformed fibril seeds (PFFs) into neurons; 2) after the generation of newly formed fibrils by endogenous aSyn and 3) during the maturation of LB-like inclusions. These studies were performed in the neuronal seeding model^54,59^, which recapitulates many of the key events and processes that govern aSyn seeding, aggregation, and LB formation^43,44^. Our studies revealed differential C-terminal cleavage and stability of exogenously added PFFs and endogenously seeded aSyn aggregates, suggesting a differential role of this PTM at various stages of LB formation. Furthermore, we show that C-terminal cleavages in combination with other PTMs play important roles in remodeling aSyn fibrils during the formation and evolution of LBs. These findings, combined with the abundance of aSyn truncated species in LBs, have significant implications for ongoing efforts to investigate aSyn pathological diversity, quantitative mapping of aSyn PTMs, and developing therapeutic strategies based on targeting the C-terminus of aSyn or proteolytic processing of this region.

## Results

### Truncation of aSyn PFFs seeds is an early event that occurs rapidly after neuronal internalization

To investigate the role of C-terminal truncation in regulating the seeding activity of aSyn, we took advantage of the neuronal seeding model^54,59^. In this model, aSyn intracellular aggregation is triggered by the addition of a nanomolar concentration (70 nM) of extracellular mouse aSyn PFFs^58^ (Figure S1), which, upon internalization, induce the formation of intracellular LB-like inclusions in a time-dependent manner^58^ in wild-type (WT) hippocampal primary neurons (Figure 1a). As previously shown^54,58^, immunocytochemistry (ICC, antibodies used are described in Figure S2) confirmed that these inclusions contained aSyn pS129 (Figure 1a-c) and were also positive for two other well-established LB markers^60^, namely ubiquitin (ub) and p62 (Figure 1d-e). We have also recently shown that the seeded aggregates are also partially phosphorylated at residues Y39, Y133 and Y136^50^. Moreover, in line with previous reports^54,58^, the increase in phosphorylated aSyn at S129 (at 7-10 days) and its colocalization with other LB markers, observed by ICC, coincides with the shift over time of endogenous aSyn from the soluble (Figure S3a) to the insoluble fraction in the PFFs-treated neurons (Figures 1f and S3b). Western blot (WB) analyses of the insoluble fraction showed a time-dependent increase in high-molecular-weight (HMW) species (∼23, 37, 40, and 50 kDa and smear >50 kDa) which started to appear 4-7 days after treatment with PFFs and were positively stained by pS129 specific antibodies (Figure 1f, middle panel). The use of the pan-synuclein antibody (SYN-1, epitope 91-99) uncovered an additional prominent band at ∼12 kDa (Figure 1f, top panel, single red asterisk), which was detected by aSyn N-ter and non-amyloid component (NAC) domain antibodies (Figure 1g-i) but not by antibodies specific for pS129 (Figure 1f middle panel) or raised against the C-terminal residues 116 to 138 (Figure 1g, i). This indicates that these species correspond to C-terminally truncated forms of aSyn (Figure S4), which persisted for up to 21 days (D21) (Figure S3c)^58^. Importantly, the cleaved aSyn species were never detected in the corresponding soluble fractions (Figure S3a), suggesting that they represent C-terminally cleaved PFFs or C-terminal truncations that occur post-fibrilization of endogenous aSyn.

**Figure 1.**
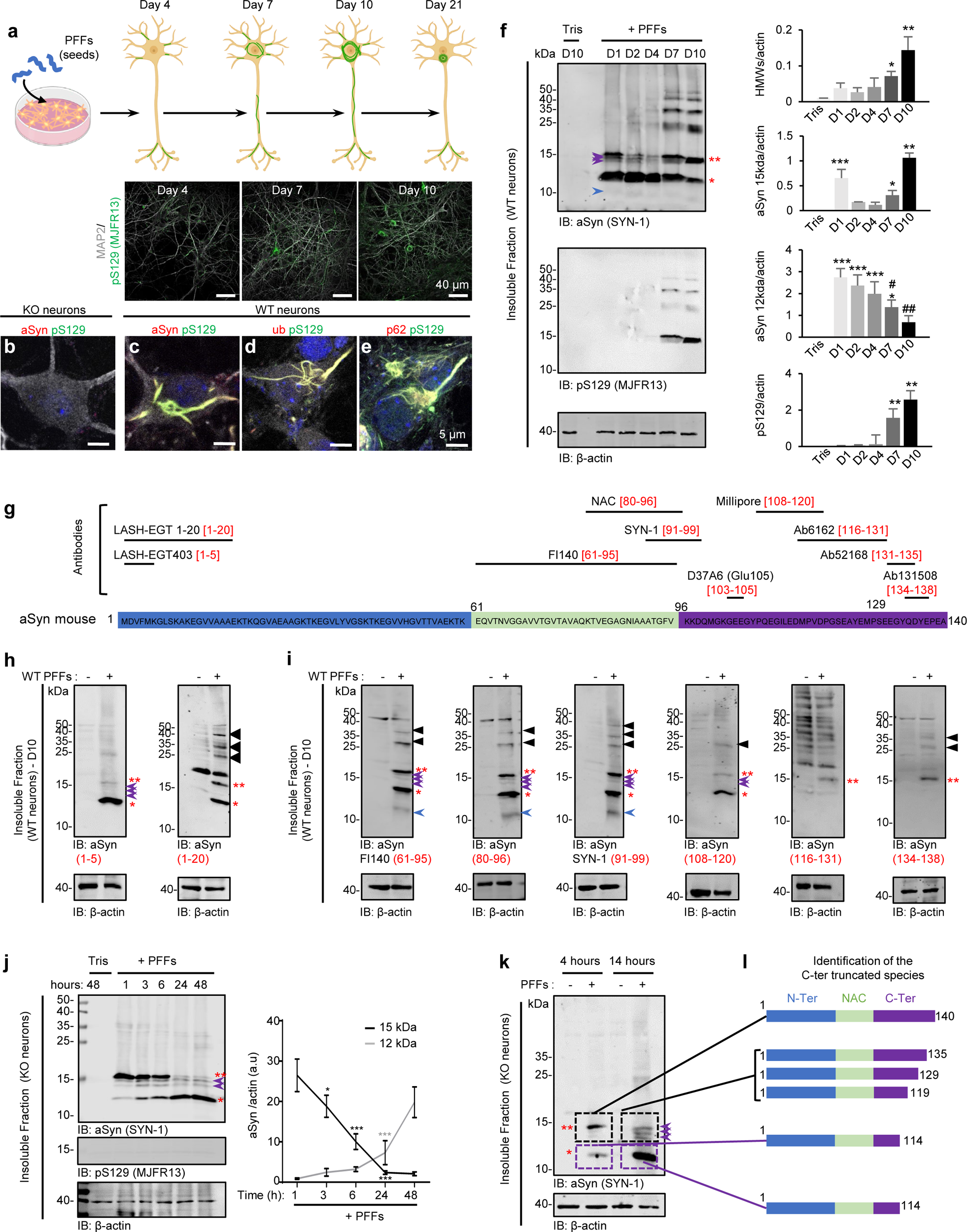
C-terminal truncation is an early event during the seeding process. **a.** Seeding model in primary hippocampal neurons. 70 nM of mouse PFFs were added to neurons at DIV 5 (day *in vitro*). Control neurons were treated with Tris buffer used to prepare PFFs. After 4 days of treatment, positive pS129-aSyn aggregates were detected in the extension of the neurons. After 7 days of treatment, the aggregates appeared in the cytosol of the neurons. The number of LB-like inclusions increased over time, as shown at 10 days of treatment. Scale bars = 20 μm. **b-e.** ICC analysis of the LB-like inclusions that formed at 10 days after adding mouse PFFs to aSyn KO neurons (b) or to WT neurons (c-e). Aggregates were detected using pS129 (MJFR13) in combination with total aSyn (SYN-1), p62, or ubiquitin antibodies. Neurons were counterstained with microtubule-associated protein (MAP2) antibody, and the nucleus was counterstained with DAPI staining. Scale bars = 5 μm. **f-g.** WB analyses of the insoluble fraction of PFFs-treated WT neurons (**f**) or PFFs-treated KO neurons (**g**). Control neurons were treated with Tris buffer (Tris). After sequential extractions of the soluble and insoluble fractions, cell lysates were analyzed by immunoblotting. Total aSyn, pS129 and actin were respectively detected by SYN-1, pS129 (MJFR13), and actin antibodies. Levels of total aSyn (15 kDa, indicated by a double red asterisk; 12 kDa indicated by a single red asterisk or HMW) or pS129-aSyn were estimated by measuring the WB band intensity and normalized to the relative protein levels of actin. Purple arrows indicate the intermediate aSyn-truncated fragments. The graphs represent the mean +/-SD of 3 independent experiments. (**f**) *p<0.01, **p<0.001, ***p<0.0001* (ANOVA followed by Tukey HSD *post-hoc* test, Tris vs. PFFs-treated neurons) and ^#^p<0.01, ^##^p<0.001 (ANOVA followed by Tukey HSD *post-hoc* test, PFFs-treated neurons D10 vs. D7 or D4 or D1). (**g**) *p<0.01, ***p<0.0001 (ANOVA followed by Tukey HSD *post-hoc* test, level of aSyn 15 kDa at 1 hour vs. other time-points or levels of aSyn 12 kDa at 1 hour vs. other time-points or Tris vs. PFFs-treated neurons). **h-i.** Insoluble fractions of aSyn KO primary neurons treated with 70 nM of mouse PFFs for 4 or 14 hours were separated on a 16% Tricine gel. After Coomassie staining, two bands at ∼15 (indicated by a black dashed box) and 12 kDa (indicated by a purple dashed box) were extracted from 16% Tricine gels (See Figure S4). Isolated bands were selected based on the size of the proteolytic fragments observed by WB (**h**) and subjected to proteolytic digestion followed by LC-MS/MS analysis. Proteomic analysis showed that aSyn fragments produced in KO neurons transduced with PFFs result from C-terminal truncation but not from N-terminal cleavage of the PFF seeds. The diagram in **i** shows the different aSyn fragments generated upon C-terminal truncation and their relative position in a WB. Three fragments (1-135, 1-129, and 1-119) were detected in the upper band sliced, and one main fragment (1-114) was found in the lower band. **j.** Epitope mapping of antibodies raised against the NAC, N-terminal, or C-terminal domains of aSyn. **k.** N-terminal antibodies raised against residues 1-5 or residues 1-20 could detect full-length (15 kDa, indicated by a double red asterisk) or truncated (∼12 kDa indicated by a single red asterisk) aSyn in the insoluble fraction of KO neurons treated for 14 hours, confirming that the N-terminal region of aSyn PFF seeds is intact after internalization into the neurons. **l.** Mapping of the C-terminal cleaved product using antibodies raised against the NAC and the C-terminal domains of aSyn. Immunoblots of insoluble fractions of KO neurons treated with aSyn PFFs showed that the fragment 1-114 generated in these neurons was well recognized by the NAC antibodies [(FL-140; 61-95) and (SYN-1; 91-99)] and a C-terminal antibody raised against the residues 108-120. However, it was not recognized by antibodies raised against peptides bearing residues after 116 in the C-terminal domain [(ab6162; 116-131); (ab131508; 134-138) and (ab52168; 131-135)]. Altogether, our data demonstrate that after internalization, aSyn PFF seeds are efficiently C-terminally truncated before the initiation of the intracellular seeding mechanisms.

We consistently observed that the C-terminal truncation fragments appeared rapidly after the internalization of aSyn PFFs into the neurons and before the first newly formed aggregates were detected (Figure 1f, Day 1). Therefore, we sought to determine if C-terminal truncation represents an early event required to initiate the seeding process in neurons. First, we monitored the extent of PFF cleavage and PTMs of PFFs after internalization into aSyn knockout (KO) neurons by WB and confocal imaging approaches. The use of aSyn KO neurons allowed us to monitor the fate of aSyn PFFs without interference from the formation of the new aSyn fibrils and LB-like inclusions since these processes do not occur in the absence of endogenous aSyn^54^ (Figures 1b and Figure S3b). Interestingly, the internalized aSyn PFFs did not undergo phosphorylation at residue S129 (Figure S3b, d) or residues Y39, Y133 or Y136^50^, ubiquitination (Figure S3e) or show any colocalization with p62 (Figure S3f) as commonly observed with the newly formed fibrils (Figures 1c-e and S3g). These findings suggest that the exogenous aSyn PFFs are processed differently compared to the fibrils formed by endogenous aSyn. The absence of N-terminal modifications such as ubiquitination and p62 signal in the internalized aSyn PFFs suggest that modifications at the C-terminus may play an important role in priming N-terminal PTMs and/or the interactome of aSyn aggregates with other proteins in neurons. WB analyses revealed that the internalized PFFs were truncated into four fragments with apparent molecular weights (MWs) between 15 and 12 kDa during the first hours after internalization in the KO neurons (Figure 1j) with complete loss of full-length aSyn (15 kDa indicated by the double red asterisk) over time (Figure S3b) and the appearance of the 12 kDa band (single red asterisk) as the main species within 24 hours (Figure 1j). As in WT neurons (Figure 1h-i), the 12 kDa band was recognized by the Nter and the NAC antibodies but not the C-terminal antibodies targeting residues beyond amino acid 116 (Figure S3h-i). Interestingly, the 12 kDa aSyn is detected in the insoluble fraction of the PFF-treated KO neurons as the dominant species for up to 21 days post-treatment (Figure S3c).

### C-terminal truncation of aSyn PFFs seeds occurs at multiple sites, leading to the accumulation of primary aSyn truncated at residue 114

To more precisely map the cleavage sites of mouse aSyn PFF seeds, aSyn bands from the insoluble fractions were digested by trypsin or Glu-C, and liquid chromatography-tandem mass spectrometry (LC-MS/MS) was performed as described in materials and methods^61^. In line with the WB data in Figure 1, our proteomic analyses demonstrated that while PFFs had an intact N-terminal domain, they were cleaved at several sites within the C-terminal domain: D135, S129, and D119 were detected in the upper band extracted at ∼13-15 kDa, whereas a cleavage occurred at E114 in the band around ∼12 kDa (Figures 1k and S4a-b). LC-MS/MS analyses also established that the main fragment that accumulates as the predominant species in neurons seeded with mouse PFFs ends at residue 114 (1-114, MW =11567.20 Da) (Figure S4c). Interestingly, similar cleavages, including the 1-114 fragment, were identified, using WB and proteomic approaches, for human PFFs added to the KO neurons (Figure S4d-f) or in the human mammalian HeLa cell line (Figure S4g-k). Interestingly, the cleavage sites of the human PFFs were almost identical to those previously identified in aSyn human neuroblastoma cell line seeding model^56^ or in LBs from human brain tissue^2,62^ (Figure S4l-m). Our results suggest that the C-terminal truncation of the internalized aSyn PFFs represents a generic cellular response to the uptake of PFFs that could enhance their seeding activity or is necessary to signal their internalization.

### aSyn PFFs seeds are cleaved in the endo-lysosomal pathway or in the cytosol

To better understand where PFFs cleavage occurs, we next performed confocal imaging analyses using mouse PFFs fluorescently labeled with Atto 488 (PFFs^488^) (Figure S1). Consistent with previous findings^63–65^, we observed that the PFFs were predominantly internalized through the endo-lysosomal pathway and accumulated in late endosomes that were positively stained for Lysosomal-associated membrane protein 1 (LAMP1) marker (Figure 2a). Additional staining using N-ter (1-20 and 34-45) or C-ter (134-138 and 116-131) aSyn antibodies revealed that C-terminal cleavage occurs in the LAMP1-positive organelles (Figures 2b and S5a-b) with ∼15% and ∼45% (Figure S5c) of the PFFs^488^ seeds not being recognized by the C-ter (116-131) antibody at D1 and D3, respectively (Figure S5b-c). This suggests that some of the PFFs are processed in the endo-lysosomal compartments, which contain several proteases that are known to cleave aSyn^66–69^, such as cathepsin^70^ and asparagine endopeptidase^32,70,71^. In line with this hypothesis, the activity of cathepsin B (Figure 2c), but not cathepsin D and L (Figure S6a-b), was significantly increased during the first 6 hours after the addition of the PFF to WT or aSyn KO neurons (Figure S6). Previous studies have also implicated the endo-lysosomal asparagine endopeptidase (AEP) in the cleavage^32,70,71^ of aSyn at residue 103 to generate a shorter C-terminal fragment 1-103. Therefore, we also assessed the activity of this enzyme and whether the PFFs are cleaved at residue 103. The enzymatic activity of AEP was significantly increased in both aSyn KO and WT neurons during the first 24 hours after the addition of the PFFs (Figure S6c-d). Intriguingly, an additional band at ∼10 kDa was detected in the insoluble fraction of the WT (Figure 1i, blue arrow) and KO (Figure S3i, blue arrow) PFFs-treated neurons by SYN-1 antibody but not by the N- or C-terminal antibodies targeting residues, 1-5 and 1-20 or between 108 and 138, respectively (Figure S7a, blue arrow). This ∼10 kDa fragment exhibited similar migration in SDS-PAGE gels as recombinant aSyn 1-103, was recognized by the N103 antibody^71^ (Figure S7b and c) and was detected at a low, albeit similar, level from D1 to D21 (S7a, SYN-1 antibody). These results demonstrate the generation of aSyn 1-103 during the processing of PFF might result from the AEP cleavage as previously described^32,70,71^ in the neurons (Figure S7), and suggest that the 1-103 fragment results from the truncation of a small subset of the PFFs.

**Figure 2.**
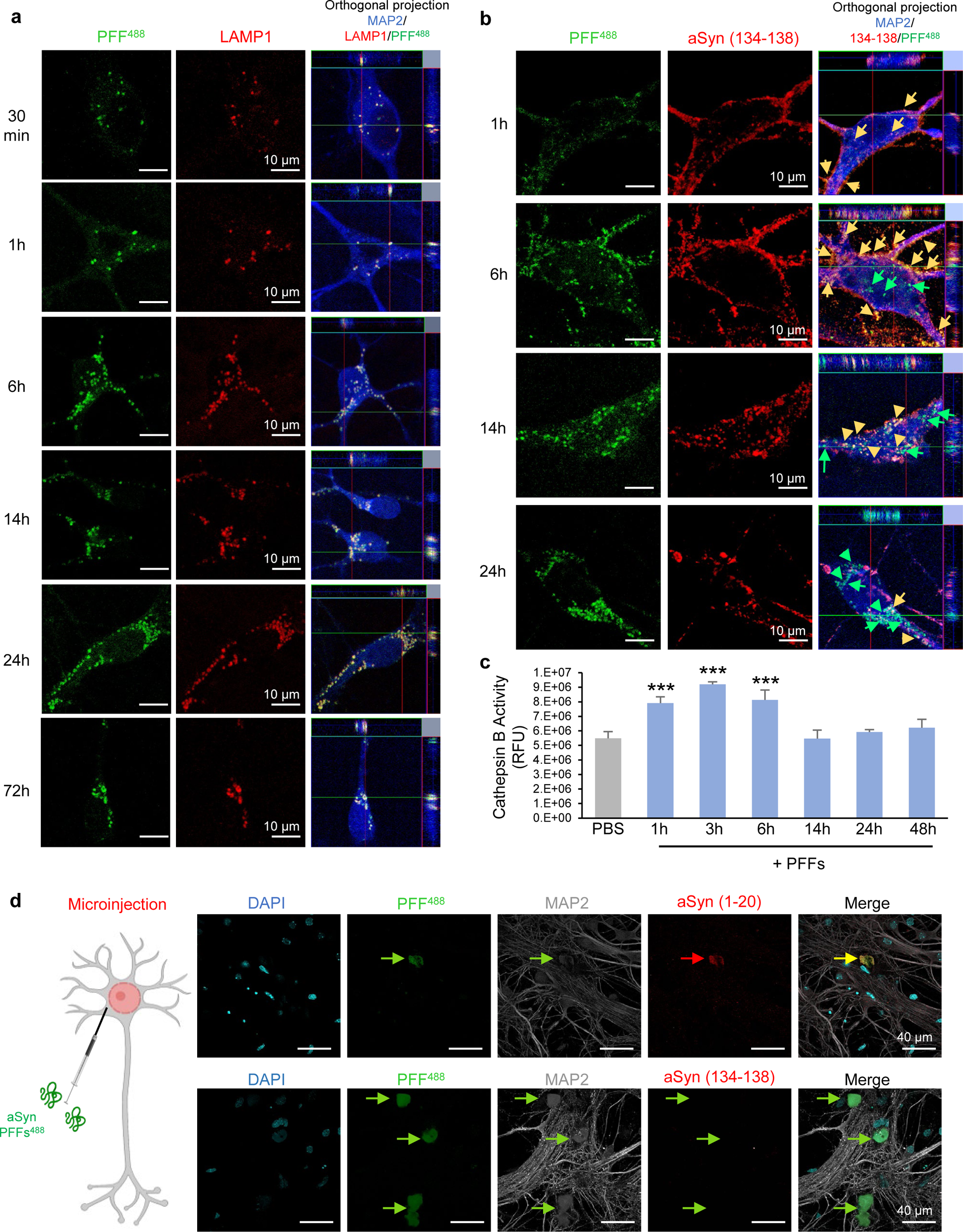
aSyn PFFs seeds are predominantly internalized via the endo-lysosomal route and are processed both in the endolysosomes and the cytosol of the neurons. **a-b.** aSyn KO neurons were treated for up to 72 hours with WT fluorescently labelled PFFs^488^. The internalization and the truncation of the seeds were evaluated by confocal imaging. **a.** One hour after addition to the KO neurons, we observed that most of the intracellular PFFs^488^ were co-stained by an antibody raised against the extremity of aSyn C-terminal domain (epitope: 134-138, yellow arrows). C-terminal truncation of the seeds over time was confirmed by the loss of detection of the seeds by the C-terminal aSyn antibody (134-138, red; green arrows). **b.** The internalization of the seeds via the endo-lysosomal pathway was confirmed by the detection of the fluorescently labelled PFF^488^ seeds in LAMP1-positive (late endosome, red) compartments overtime. **a-b.** Neurons were counterstained with MAP2 antibody and the nucleus with DAPI stain. Scale bars = 10 μm. **c.** Cathepsin B activity was measured in KO neurons treated with 70 nM of WT PFFs seeds for up to 48 hours. Control neurons were treated with Tris buffer. The graphs represent the mean +/-SD of 3 independent experiments. *p<0.01, **p<0.001, ***p<0.0001 (ANOVA followed by Tukey HSD *post-hoc* test, Tris vs. PFFs-treated neurons). **d.** Truncation of aSyn PFFs in the cytosol is confirmed by microinjection. The diagram on the left-hand side shows the experimental approach used to microinject WT PFFs^488^ in KO neurons. Cells were fixed after 24 hours and immunostained using the N-terminus antibody (aSyn 1-20) or C-terminus antibody (aSyn 134-138). Confocal imaging showed that WT PFFs^488^ were detected by the N-terminus antibody (yellow arrows, merge), but not the C-terminus antibody (green arrows, merge). Neurons were counterstained with MAP2 antibody and the nucleus with DAPI stain. Scale bars = 40 μm.

Next, we assessed whether the cleavage of the aSyn PFFs could also occur once they escape from the endocytic pathway^63,72^ or upon their uptake by alternative routes, including receptor-mediated processes^73–75^. Toward this goal, we used a microinjection technique^76^ to deliver fluorescently labelled PFFs^488^ directly into the cytoplasm of individual KO neurons (Figure 2d). 24 hours post-injection, the neurons were fixed and stained with N-terminal (epitope: 1-20) and C-terminal (epitope: 134-138) aSyn antibodies. The colocalization of the aSyn PFFs^488^ signal with aSyn was observed using antibodies that target the N-terminal residues of aSyn (Figure 2d, yellow arrow). An antibody against the extreme C-terminal residues 134-138 failed to detect the PFFs^488^ (Figure 2d, green arrows). Altogether, our data confirm that aSyn PFFs are not only cleaved in the endolysosomal pathway but can also be processed once they reach the cytosol of the neurons.

### PFF cleavage and generation of aSyn truncation occurs in different cell types and rodent models of aSyn pathology formation aSyn

Next, we sought to determine whether aSyn PFFs are subjected to differential cleavage in different cell types and various models of aSyn pathology formation (Figure S8). Towards this goal, we assessed the extent of aSyn truncation in several neuronal seeding models (Figures 3a-d and S9a-c) as well as in a pure primary culture of mouse cortical astrocytes (Figure S9d). Truncation occurred in all the PFFs-treated neurons, including hippocampal or cortical primary neurons from mice (Figures 3a and S9a) or hippocampal, cortical, or striatal primary neurons from rats (Figures 3b and S9b) but not in the astrocytes (Figure S9d). WB analyses clearly showed a similar truncation pattern and efficiency in all the types of neurons with aSyn cleaved into ∼12 kDa fragments. Interestingly, we observed a similar HMW band pattern in the insoluble fraction (∼23, 37, 40, and 50 kDa) in all types of neurons.

**Figure 3.**
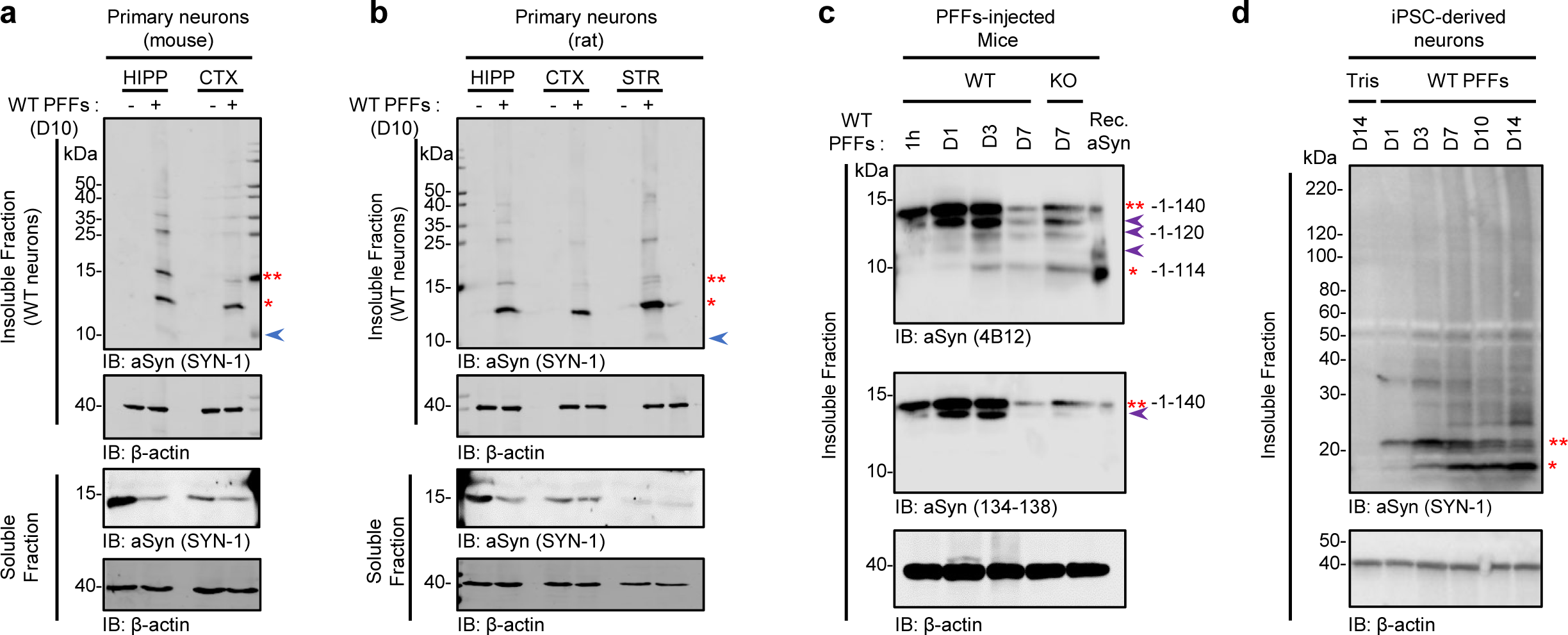
C-terminal truncation is a general phenomenon in aSyn seeding and inclusion formation in cells. WB analyses of the truncation pattern of aSyn in primary neurons (**a-b**), *in vivo* after injection of human PFFs in the striatum of WT and aSyn KO mice (**c**), in iPSC-derived neurons from a healthy control individual transduced with human PFFs (**d**). Hippocampal (HIPP) and cortical (CTX) primary neurons from mice or (**a**) hippocampal (HIPP), cortical (CTX), or striatal (STR) primary neurons from rats (**b**) were treated for 10 days with 70 nM of mouse PFFs. iPSC-derived neurons were treated with 70 nM of human PFFs for 1, 3, 7, 10, and 14 days (**d**). Control neurons were treated with Tris buffer (-). The striatum of WT or aSyn KO mice were dissected after 1 hour or 1, 3, or 7 days after injection with human PFFs^WT^ (**c**). After sequential extractions of the soluble and insoluble fractions, cell lysates were analyzed by immunoblotting. The levels of total aSyn (SYN-1, 4B12, or 134-138 antibodies) (15 kDa, indicated by a double red asterisk; 12 kDa indicated by a single red asterisk or HMW) were estimated by measuring the WB band intensity and normalized to the relative protein levels of actin (Figure S9). Purple arrows indicate the intermediate aSyn-truncated fragments.

Strikingly, a similar truncation pattern was also observed *in vivo* after injecting human PFFs in the striatum of C57BL/6J WT mice or aSyn KO mice (Figures 3c and S9c). To follow the fate of the seeds specifically, we used an antibody raised against human aSyn (clone 4B12). As early as 1 day after injection, four cleaved fragments running at sizes similar to those observed in primary neurons were detected in the insoluble striatal fractions of the rodent brains using an aSyn antibody raised against the epitope 103-108 (clone 4B12), but not with an antibody specific for the C-terminal domain (epitope: 134-138). This is consistent with these fragments resulting from the C-terminal cleavage of aSyn species.

To explore the pathophysiological relevance of our findings further, we next investigated the processing of fibrillar aSyn in human induced pluripotent stem cells (iPSCs) that were differentiated into dopaminergic neurons^77–79^. We used a line derived from a healthy control individual, which showed dense processes and expressed markers consistent with human midbrain neurons (Figures 3d and S9e-g). 24 hours after adding human PFFs to the iPSC-derived human neuronal culture (50 days *in vitro*), aSyn was cleaved into a ∼12 kDa fragment (Figures 3d and S9e), which was similar to our observations of mouse neurons. Altogether, our data demonstrate that the truncation of aSyn fibrils is a general and early event that occurs after the internalization of propagating fibrils and during the formation of intracellular LB-like aSyn inclusions in all neuronal seeding models used.

### Preventing aSyn cleavage at residue 114 does not impact the seeding capacity of PFFs in primary neurons

Given the rapid proteolytic processing of PFFs upon internalization, we initially hypothesized that the C-terminal cleavage of the PFFs might be a prerequisite for the seeding and initiation of endogenous aSyn aggregation in neurons. To test this hypothesis, we used PFFs carrying a single amino acid substitution at residues 114 (E114A) or 115 (D115A) (Figure S1, mouse E114A, and D115A PFFs), which were designed to block C-terminal cleavage at 114. Figure 4a shows that aSyn^E114A^ PFFs but not aSyn^D115A^ PFFs underwent cleavage to generate the 1-114 fragment. Therefore, we used PFFs generated from this mutant to investigate the role of C-terminal cleavage of PFFs in regulating their seeding activity. In aSyn KO neurons, PFFs^D115A^ underwent cleavage only at residues 135, 129 and 119 and the resulting fragments were cleared overtime (Figure 4b).

**Figure 4.**
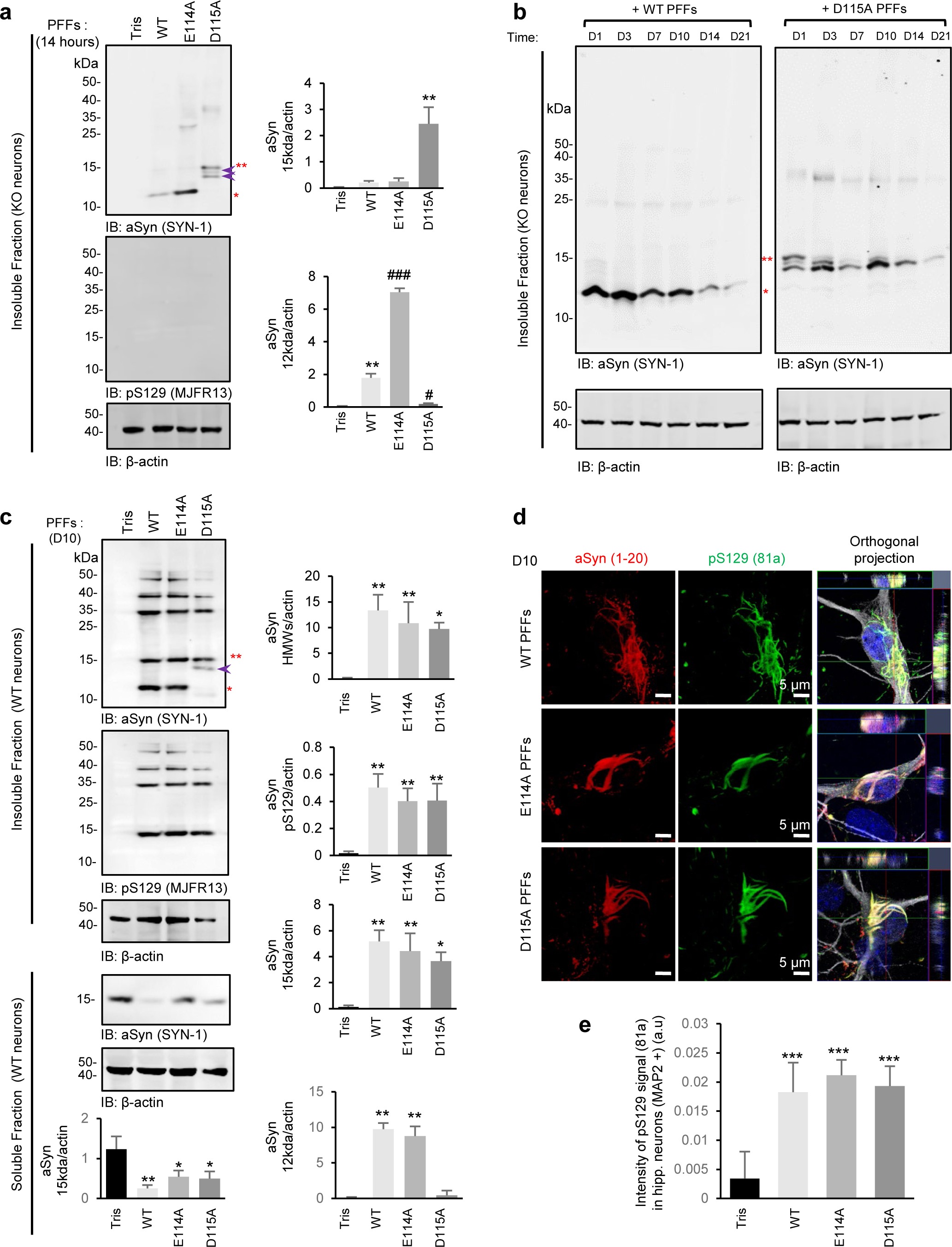
Preventing aSyn cleavage at 114 does not impact the seeding capacity of PFFs in primary neurons. **a-c.** aSyn KO neurons were treated for 14 hours (**a**) or up to 21 days (**b**) or aSyn WT neurons were treated for 10 days (**c**) with PFFs^WT^, PFFs^E114A^, PFFs^D115A^ or with PFFs^1–114^. Neurons were lysed at the indicated time, and the insoluble fractions were analyzed by WB. The levels of total aSyn (SYN-1 antibody) were estimated by measuring the WB band intensity normalized to the relative protein levels of actin. A double red asterisk indicates the 15 kDa species (full-length), and a single red asterisk indicates the C-terminal truncated species running at 12 kDa. The purple arrows indicate the intermediate aSyn-truncated fragments. **d-e**. Newly formed inclusions were detected using pS129 antibody (81a) in WT neurons after 10 days of treatment. Neurons were counterstained with MAP2 antibody, and the nucleus was counterstained with DAPI staining. (**d**) Representative confocal images. Scale bar = 5 μm. (**e**) Quantification of images acquired by a high-throughput wide-field cell imaging system. For each independent experiment, duplicated wells were acquired per condition, and nine fields of view were imaged for each well. Each experiment was reproduced at least 3 times independently. Images were then analyzed using Cell profile software to identify and quantify the level of LB-like inclusions (stained with pS129 antibody, 81a clone) formed in neurons (MAP2-positive cells). The graphs (**a**, **c, e**) represent the mean +/-SD of three independent experiments. p<0.01=*, **p<0.001, ***p<0.0001 (ANOVA followed by Tukey HSD *post-hoc* test, Tris vs. PFFs-treated neurons). ^#^p<0.05, ^##^p<0.005 (ANOVA followed by Tukey HSD *post-hoc* test, PFFs^WT^ vs. mutants PFFs-treated neurons).

Next, we compared the seeding capacity of PFFs^E114A^ and PFFs^D115A^ to that of PFFs^WT^ after 10 days of treatment in WT neurons. PFFs^WT^, PFFs^E114A^, PFFs^D115A^ all induced a similar level of seeding in WT neurons (Figure 4c-e). These findings were confirmed by the WB analyses of the insoluble fractions extracted from WT neurons treated with PFFs^WT^, PFFs^E114A^, or PFFs^D115A^, in which a similar level of pS129 and HMW signals was detected (Figure 4c). Consistent with the data, we observed similar reductions in the levels of soluble proteins in neurons treated with PFFs^E114A^ and PFFs^D115A^ and PFFs^WT^ (Figure 4c). These findings were further confirmed by quantitative ICC. Furthermore, we did not observe significant differences in the pS129 levels (Figure 4e) or the morphology of the newly formed aggregates (Figure 4d) in neurons treated with PFFs^WT^ or with PFFs mutants for 10 days. This is in line with previous studies reporting that the deletion of various aSyn regions other than the NAC domain does not inhibit the formation of seeded aggregates^53,54^.

Our results demonstrate that blocking the C-terminal truncation of the seeds does not prevent the seeding and the recruitment of endogenous aSyn or the formation of aggregates in neurons.

### The newly formed aSyn fibrils are processed differently and undergo more complex PTMs compared to exogenous PFFs

Having established that C-terminal cleavage is not essential for the initial seeding events, we then sought to determine if the newly formed fibrils are also subjected to proteolysis and whether this modification is important for the transition from fibrils to LB-like inclusions^58^. Toward this goal, we used fluorescently labelled PFFs^488^ to enable distinguishing exogenous PFFs from newly formed fibrils. Using ICC and confocal imaging, we observed that the internalized seeds that appear to be incorporated in/or colocalized with the newly formed aggregates represent a minor species (Figure S10a). This suggests that the majority of the fibrils present in the seeded aggregates are derived from newly recruited endogenous aSyn. Next, to discriminate the PFF seeds from the newly formed fibrils by WB analyses, mouse WT neurons were treated with human PFFs for 10 days, and the insoluble fraction was analyzed by WB using a combination of human- and mouse-specific antibodies, 4B12 and Glu105, respectively (Figure S10b). Our data clearly indicated that ∼90% of the ∼12 kDa species detected by WB were composed of the exogenous PFF seeds, while the newly formed aggregates were represented by the HMW bands (∼23, 37, 40, and 50 kDa) but also by the species > 170 kDa and trapped in the stacking gel (Figure S10b). These results and the absence of a truncated mouse aSyn band suggest that the newly formed and post-translationally modified fibrils are more stable and exhibit higher resistance to sodium dodecyl sulfate (SDS) compared to the aSyn PFF seed, which readily disassociates in SDS buffers.

We next used quantitative proteomic analyses (Figures 5a-b and S10c) combined with WB (Figures 1h-i and 5c) and ICC (Figures 5d-g and S10d-e) analyses to determine how the newly formed fibrils are cleaved during the aggregation process in comparison to the exogenous PFFs. At day 10, both the proteomic (Figures 5a-b and S10c) and WB analyses (Figures 1h-i and 5c) showed that the processing of aSyn in WT neurons was, in all respects, similar to that observed for the PFFs added to the KO neurons (Figures 1j and S3a), i.e. presence of aSyn species, that accumulate at ∼12 kDa (Figures 1h-i and 5c) were cleaved after residue 114 and bore an intact N-terminal domain (Figures 1h-i and 5c). In addition, mass spectrometry analyses revealed that the HMW newly formed aggregates were composed of both full-length aSyn and C-terminally cleaved aSyn species (1-114 and 1-119) (Figures 5a-b and S10c). All aSyn species present in the HMWs were also ubiquitinated at multiple lysine residues from D7 to D21 post-treatment (Figure S10c).

**Figure 5.**
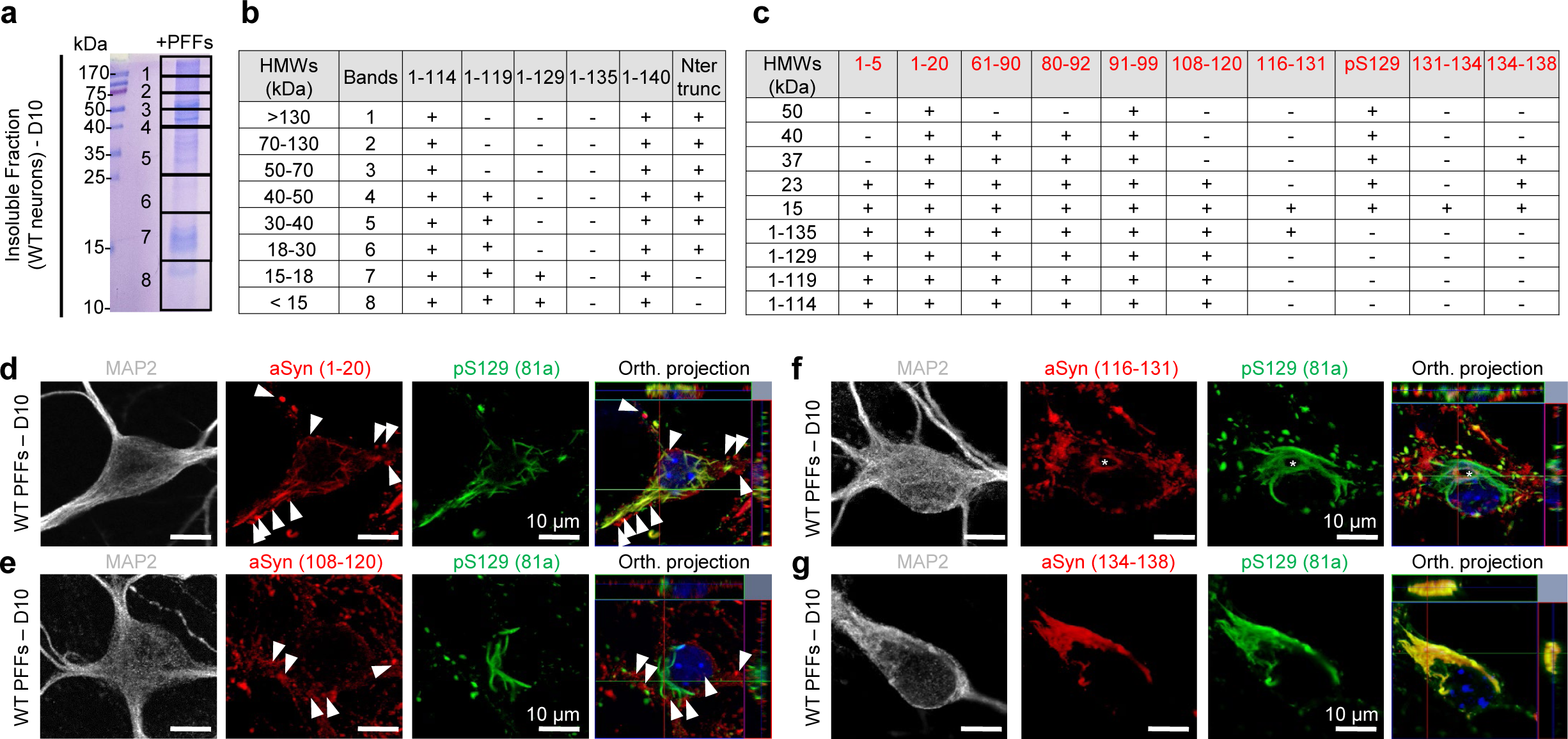
The newly formed aSyn fibrils are processed differently and undergo more complex PTMs compared to exogenous PFFs. **a-g.** Identification of C-terminal truncated fragments by proteomic analysis (**a-b**), WB (**c** and Figure 1h-i) or confocal imaging (**d-g**). WT neurons were treated with 70 nM of mouse PFFs for 10 days. **a-b.** The insoluble fractions of PFFs-treated neurons were separated on a 16% Tricine gel. After Coomassie staining, 8 bands were extracted from the Tricine gel (**a**). Isolated bands were subjected to proteolytic digestion using trypsin for C-terminal truncation identification^132^, followed by LC-MS/MS analysis. **b.** Proteomic analyses showed the presence of the 1-114 and 1-119 C-terminal truncated fragments in the HMW species. **c.** Table summarizing the capacity of NAC domain, N-terminal, and C-terminal antibodies to detect full-length aSyn (15 kDa), the C-terminally cleaved fragment of aSyn (∼12 kDa), and the HMW formed in WT neurons after 10 days of treatment with PFFs (see WB in Figures 1h-i). **d-g**. Antibody mapping of the newly formed inclusions by confocal imaging using pS129 antibody (81a clone) in combination with N-terminal (**d**, epitope 1-20) or C-terminal (**e-g**, respective epitopes 108-120, 116-131, or 134-138) antibodies revealed the presence of aSyn-positive aggregates that were not pS129 positive or only partially phosphorylated at S129 residue (**e-f**). Neurons were counterstained with MAP2 antibody and the nucleus with DAPI stain. The white arrows indicate the sub-populations of aggregates localized near the pS129-positive inclusions, and the white asterisk those inside the pS129-positive filamentous structures. Scale bars = 10 μm.

Next, we compared the immunoreactivity of aSyn seeded aggregates by ICC using a set of antibodies targeting the N- and C-terminal regions of aSyn (Figures 5d-g and S10d-e). After 10 days of treatment, the N-terminal antibody (1-20) and the C-terminal antibodies raised against the 108-120 or the 116-131 region uncovered the presence of aSyn-positive accumulations that were not pS129 positive (Figure 5d-e) or only partially phosphorylated at the S129 residue (Figure 5f). These sub-populations of aggregates were either localized near the pS129-positive inclusions (Figure 5d-e, white arrows) or inside the pS129-positive filamentous structures (Figure 5f, white asterisk). As expected, the C-terminal antibody (aa 134-138) that recognizes only full-length aSyn by WB (Figures 1h-i and 5c) detected the pS129 positive aggregates (Figure 5g). These observations suggest that some of the newly formed aggregates are not yet phosphorylated at S129 or that the detection of pS129-modified fibrils is masked due to the presence of neighbouring modifications.

We have recently developed and validated an expanded aSyn antibodies toolset to profile aSyn pathology in post-mortem human brain tissues and in the neuronal and in vivo seeding model^50^. Using these antibodies, we showed that while the PFFs seeds are exclusively C-terminally truncated, the seeded aggregates contain a diversity of aSyn species phosphorylated mainly at S129 residue but also partially phosphorylated at N-ter and C-ter tyrosine residues (Y39, Y125, Y133, Y136)^50,80–82^, or nitrated at residue Y39^50,83^, or C-terminally truncated^2,45,48,50,62,84,85^. Altogether, our data demonstrate that the newly formed fibrils are processed differently in neurons and exhibit different biochemical properties compared to the exogenous recombinant PFFs, which undergo only C-terminal cleavage but lack phosphorylation, ubiquitination or other PTMs.

### C-terminal truncations of aSyn occur post-fibrillization, induce fibrils lateral association and changes in their interactome

Our previous correlative light electron microscopy (CLEM) studies showed that the process of inclusion formation in seeded neurons is accompanied by the transient lateral association of the newly formed fibrils followed by the sequestration of lipids, organelles, and endomembranes structures into LB-like structures^58^. Given that the highly negatively charged C-terminal domain is exposed and decorates the surfaces of the fibrils, we hypothesized that C-terminal cleavages of the newly formed fibrils could drive their lateral association. We postulated that this post-fibrillization processing might be required for the packing and sequestration of fibrils during LB formation and could explain the accumulation of C-terminal cleavage aSyn in the core of LBs^43,45,86^. Furthermore, the C-terminal domain of aSyn serves as a major interactome hub for aSyn monomers and fibrils. Therefore, removal of this domain is expected to disrupt the interactome of the fibrils, which is expected to regulate the interactions of the fibrils with other cellular proteins and components that co-accumulate with aSyn within LBs.

To investigate the interplay between C-terminal cleavage of fibrils and their interactome, lateral association and LB formation, we first investigated changes in the morphology of the newly formed fibrils over time. We imaged neurons treated for 10, 14, or 21 days with mouse WT PFFs by CLEM (Figure 6). Previously, we showed that at 7 days, newly formed fibrils exist as single long filaments with lengths ranging between 600 nm and up to 2μm^58^. As shown in Figure 4, at D10 (Figure 6a, red arrows) and D14 (Figure 6b, red arrows), the newly formed aSyn fibrils were shorter in length as described previously^58^ and reorganized into tightly packed bundles of fibrils. These clusters of fibrils were not formed by randomly arranged fibrils but rather by fibrils that were closely associated and aligned in parallel. From D14, organelles such as mitochondria and endolysosomal-like vesicles were detected close to the laterally associated bundles of fibrils. At D21, long filamentous-like structures (Figure 6c) and ribbon-like inclusions (Figure 6d) of aSyn newly formed fibrils were still appearing as tightly packed bundles of fibrils that appeared to be closely associated with mitochondria, autophagosomes, and endoplasmic reticulum. Conversely, in neurons in which aSyn aggregates have completely transitioned into inclusions with a round LB-like morphology (Figure 6e), the laterally associated fibrils were no longer observed, and only short aSyn filaments randomly organized were detected in the center of the inclusion. Our data suggest that once fibril fragmentation has reached its minimum size, the lateral association and packing of the newly formed fibrils are no longer sustainable or require fibrils disassembly^87^. Alternatively, other PTMs or cellular proteostasis mechanisms, such as the action of chaperones^88–90^ could induce disruption of fibril-to-fibril association or induce the disaggregation of the fibrils during LB formation and maturation.

**Figure 6.**
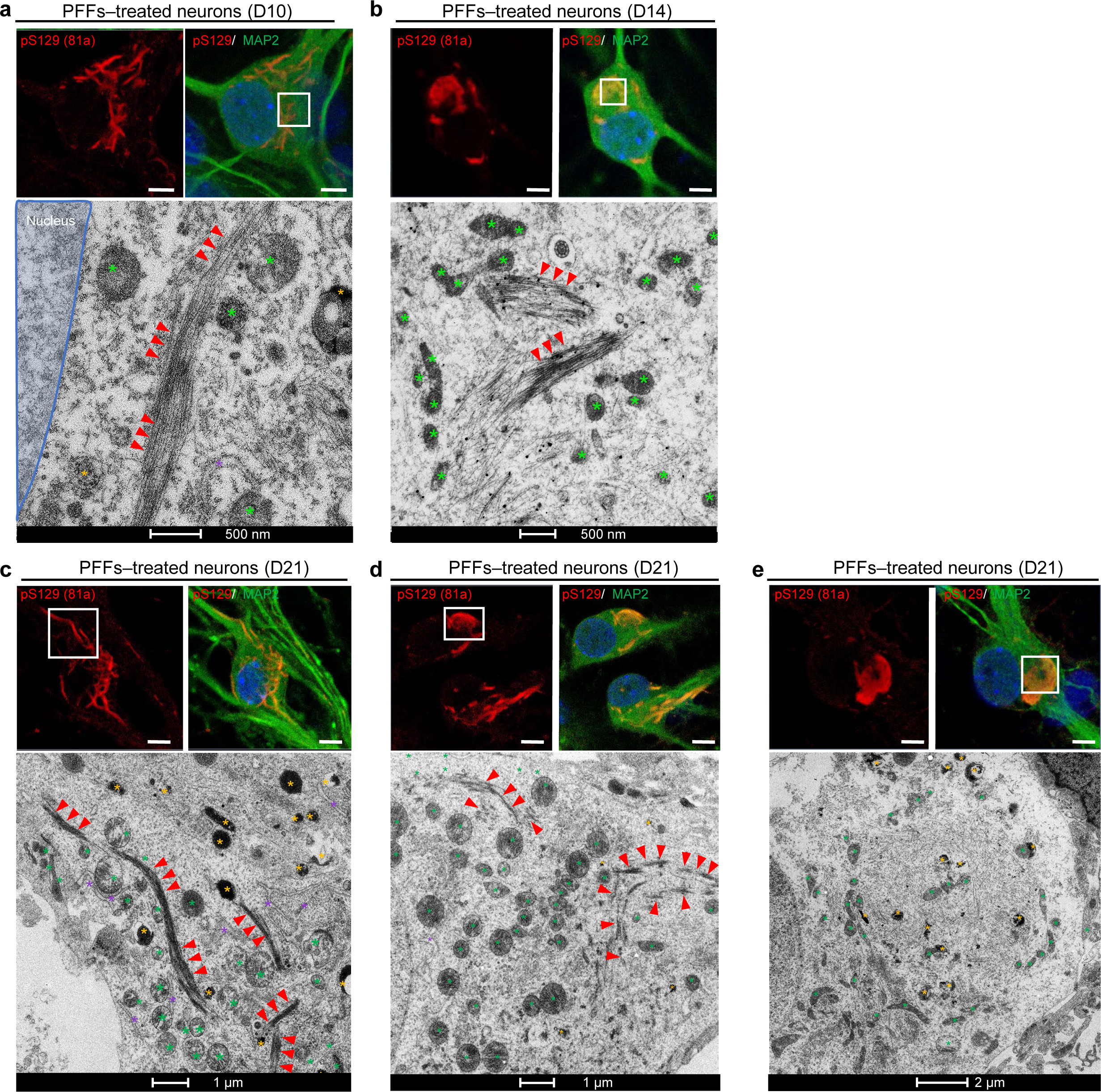
Newly formed fibrils undergo lateral association and fragmentation over time. PFFs were added for 10 (a), 14 (b), and 21 (c-e) days to the extracellular media of hippocampal neurons plated on dishes with alpha-numerical searching grids imprinted on the bottom, which allowed the localization of the cells. At the indicated time, neurons were fixed and imaged by confocal microscopy (top images), and the selected neurons were embedded and cut by an ultramicrotome. Serial sections were examined by EM. Representative newly formed aSyn fibrils are indicated with a red arrow. Autophagolysosomal-like vesicles are indicated by a yellow asterisk, and mitochondrial compartments are indicated by a green asterisk. The nucleus is highlighted in blue. **a, b.** Scale bars = 500 nm; **c-d**. Scale bars = 1 μm; **e.** Scale bar = 2 μm.

Our observation that the newly formed fibrils are composed of a mixture of both full-length (pS129 immunoreactive) and C-terminally truncated aSyn suggests that the latter occurs after aSyn fibrillization. One alternative explanation is that the C-terminally truncated species may represent the early aggregates formed by C-terminally truncated monomers, which then seed the aggregation of the full-length protein. We hypothesized that if the latter is the predominant mechanism, we should not see major changes in the aSyn C-terminal-interactome over time as the number of truncated seeds within the fibrils will not change over time. However, if C-terminal cleavages occur after fibrillization and regulate the transition of fibrils to LB-like inclusions, then we should see a time-dependent decrease in aSyn C-terminal interactome. To test this hypothesis, we first identified the interacting partners of intact recombinant PFFs by incubating cortical mouse brain lysates with PFFs derived from aSyn that was site-specifically biotinylated at its N-terminal residue (Figure S11a-g). Biotinylated fibrils were then specifically pulled down using streptavidin beads, and putative interacting partners were determined by LC-MS/MS analysis (Figure S11i). As expected, ∼80% of the previously reported putative aSyn C-terminal interacting partners were found to bind to full-length PFFs in the pull-down assay (Figure S11i).

Next, we conducted quantitative proteomic studies on the insoluble fraction of PFFs-treated neurons (Figure S11i-j). We monitored the changes in the levels of the 76 previously reported C-terminal aSyn interacting proteins over time (Figure S11i, left-handed column) and determined which C-terminal interacting proteins were lost under conditions where truncations were detected in the neuronal seeding model (Figure S11i, purple boxes). After 14 days of PFF treatment, ∼25% of the proteins identified as putative C-terminal interactors were significantly enriched in the insoluble fraction of the PFFs-treated neurons (Figure S11i-j, green boxes). After 21 days, only 9% of these C-terminal interactors were still present in the insoluble fraction of the PFFs-treated neurons (Figure S11i-j, green boxes). These findings are in agreement with our hypothesis that C-terminal truncations of aSyn occur post-fibrillization and lead to the disruption of the aSyn interactome involving the C-terminal region of the protein. This mostly includes proteins involved in the cytoskeleton architecture (e.g., MAP1B, Tubb3, Tubb4, Tubb5, Tubb6, Myh10 and Rap1a), suggesting that the remodeling of the newly formed fibrils by C-terminal cleavage leads to a loss of physical interactions with the cytoskeleton and other partners.

### C-terminal truncation promotes the lateral association of aSyn fibrils *in vitro*

To test our hypothesis that C-terminal cleavage of fibrils promotes their lateral association, we investigated the impact of C-terminal truncations on the morphology of aSyn fibrils in vitro. First, we assessed the *in vitro* structural properties of the fibrils formed after the incubation of full-length aSyn monomers (1-140) or C-terminal truncated variants corresponding to those detected in our neuronal model and the human brain (1-135, 1-133, 1-124, 1-120, 1-115, or 1-111) at 37°C under agitation conditions (Figure 7a-b). After 6 days, EM imaging showed that the PFFs^1–140^ and PFFs^1–135^ formed predominantly well-dispersed single fibrils. In contrast, PFFs formed by monomers derived from shorter C-terminally truncated fragments (PFFs^1–124^, PFFs^1–120^, PFFs^1–115^, and PFFs^1–111^) exhibited a high tendency to associate laterally and pack together (Figure 7b), forming micrometre-large dense aggregate clumps. The PFFs^1–133^ exhibited an intermediate behavior with less dispersed fibrils compared with PFFs^WT^ (Figure 7b). As expected, single (Δ111-115, Δ120-125 or Δ133-135) or double deletions (Δ111-115Δ133-135) that do not substantially decrease the number of negative charges compared to WT aSyn did not favor the lateral association of these PFFs (Figure 7b). Interestingly, well-dispersed single fibrils were observed when the charge state of PFFs^1–114^ was changed to -14 [comparable to that of the WT protein (-12)] by aggregating the protein at pH 10 (Figure 7c). However, upon the adjustment of the pH to 7.5, the PFFs^1–114^ underwent rapid lateral association and packed together in aggregate clusters. These results demonstrate that the level of lateral association, the size, and the density of the resulting fibrils clumps appeared to depend on the charge state of the C-terminal domain. Monomers with the shortest C-terminal domains (i.e., the lowest number of C-terminal negative charges) exhibit a higher propensity to undergo lateral association and form clusters of densely packed fibrils.

**Figure 7.**
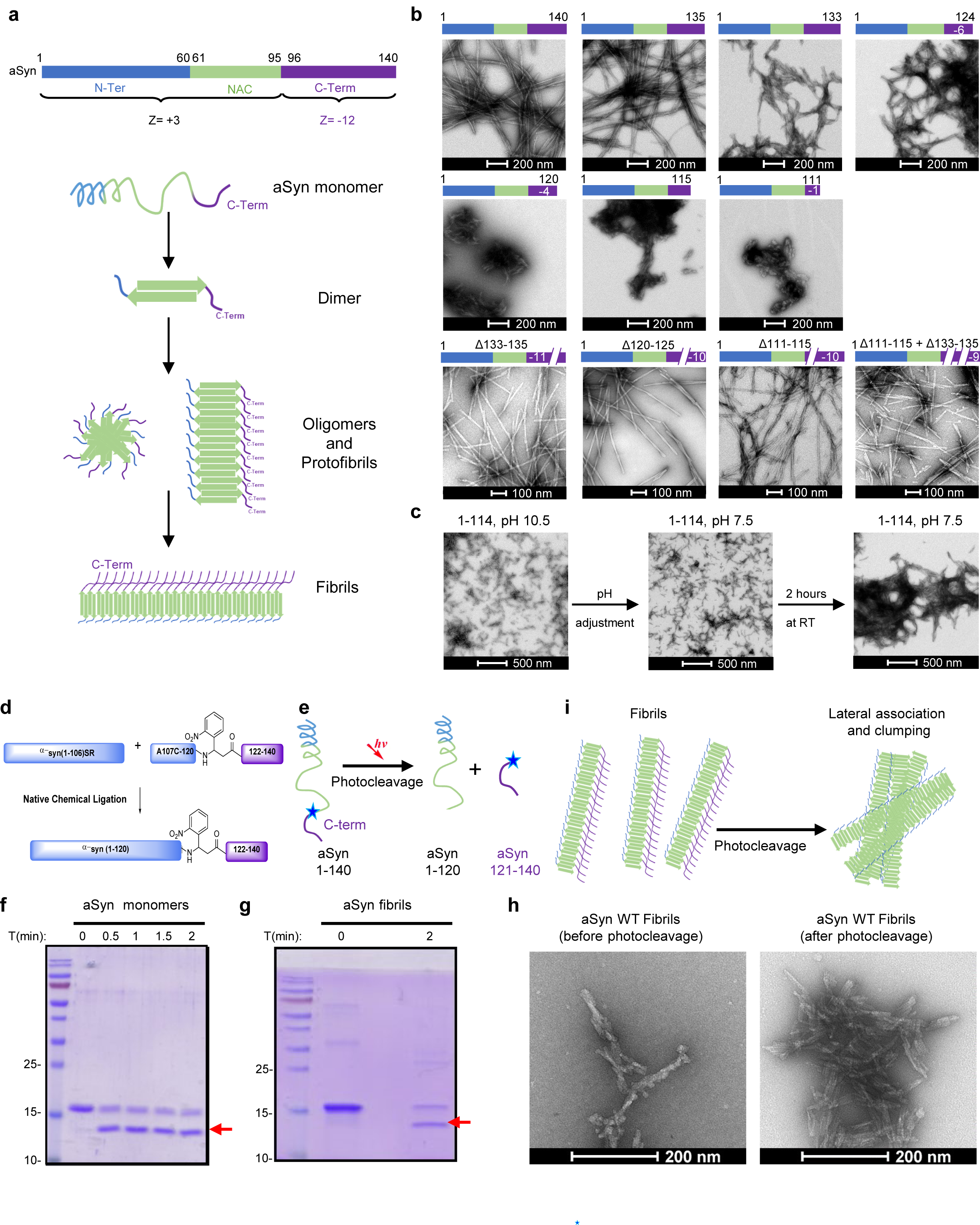
C-terminal truncation promotes the lateral association of aSyn PFFs *in vitro*. **a.** aSyn protein structure. The acidic C-terminal domain (purple, residues: 96-140) is negatively charged (z=-12), while the N-terminal domain (green, residues: 1-60) together with the central hydrophobic core named NAC (red, residues; 61-95) are positively charged. Interaction of unfolded aSyn monomers leads to the formation of 1) dimers that grow into 2) oligomers and protofibrils, which convert to 3) fibrils. During fibrilization, the C-terminal domain is exposed on the surface. **b.** Full-length recombinant aSyn, C-terminal truncated aSyn (1-135, 1-133, 1-124, 1-120, 1-115 or 1-111), and aSyn with single (Δ111-115, Δ120-125 or Δ133-135) and double deletions (Δ111-115Δ133-135) were incubated at 37°C for 6 days under shaking conditions and imaged by EM. We observed that the removal of charges in the C-terminal truncated proteins induces lateral association of the PFFs, resulting in highly packed fibrils. The level of lateral association is stronger for the proteins with a lower number of negative charges in their C-termini. The number of C-terminal charges is indicated on the right side of each diagram. Scale bar = 200 nm. The lateral association is not observed for the PFFs^1–135^, for which the number of negative charges remains comparable to that of the WT protein. Likewise, PFFs with single (Δ111-115, Δ120-125 or Δ133-135) and double deletions (Δ111-115Δ133-135) do not laterally associate, as the number of negative charges remains comparable to that of the WT protein. Scale bars = 100 nm or 200 nm. **c.** Fibrils formation was induced by incubating C-terminally truncated aSyn 1-114 at 37°C and pH 10.5 in 50 mM Tris and 150 mM NaCl under constant agitation at 1000 rpm on an orbital shaker. After 6 days, PFFs were sonicated, and the pH was re-adjusted to 7.5. After 2 hours at room temperature, PFFs laterally associated and formed large, dense aggregate clumps. Scale bars = 500 nm. **d-e.** Semisynthesis (**d**) and strategy (**e**) of photocleavable aSyn at position 120. **f-i.** Photolysis of aSyn monomers (**f**) and PFFs (**g**) results in the rapid generation of C-terminally truncated aSyn. **h**. EM confirmed that photolysis of photocleavable aSyn PFFs enabled the lateral association and clumping of the PFFs, as also depicted in the diagram in **i**. Scale bar = 200 nm.

Given that we postulated that C-terminal cleavages also occur after aSyn fibrillization, we also assessed the effect of site-specific post-fibrillization cleavage on fibril morphology and lateral association with great precision. Towards this goal, we developed a novel semisynthetic form of aSyn with a (2-nitrophenyl)propanoic acid between residues 120 and 122 (aSyn-D121Anp), allowing the temporal regulation of site-specific aSyn cleavage. The incorporation of this unnatural amino acid allows the photocleavage of aSyn, specifically at position 120 (Figure 7d-e). We first verified that photoactivation leads to site-specific cleavage of the monomeric aSyn-D121Anp using ultraviolet (UV) light. As shown in Figure 7f, cleavage of aSyn monomers occurred within a few seconds (Figure 7f). Next, we prepared fibrils derived from aSyn-D121Anp PFFs (Figure 7g) and showed that they could be successfully cleaved following exposure to UV light. As shown in Figure 7h, the site-specific C-terminal photocleavage of full-length aSyn-D121Anp PFFs and removal of the last 20 amino acids led to their tight lateral association and the formation of fibrils that resemble those seen with the C-terminal truncated proteins (Figure 7h-i). Taken together, our findings show that C-terminal cleavages of monomeric or fibrillar aSyn promote the lateral association of fibrils.

### Calpains 1 and 2 play active roles in the remodeling of aSyn fibrils and the formation or maturation of LBs

Having established that aSyn C-terminal cleavage plays a critical role in the lateral association and determined the morphology of the newly formed fibrils and their remodeling, we next sought to investigate the impact of blocking C-terminal cleavage of endogenous aSyn on PFF-mediated seeding and formation of LB-like inclusions. As a first step towards achieving this goal, we investigated which proteases were involved in regulating the C-terminal cleavage of aSyn fibrils.

Several enzymes have been reported as potential proteases that regulate aSyn C-terminal truncations^20,52,91–95^, including cathepsin D^68,96^, neurosin^97^, metalloproteinases^98–100^, caspase 1^29,93,95^, calpain 1^52,91,92^ and calpain 2^92,101^. Among these, it has been shown that calpains 1 and 2 cleave aSyn fibrils predominantly at the amino acids 114 and 122^92^ *in vitro*. Consistent with this report, we confirmed that aSyn fibrils were cleaved by calpain in the 111-115 region (Figure S12a).

To evaluate the potential contribution of calpains 1 and 2 in regulating aSyn seeding and inclusion formation, we first investigated whether these enzymes were activated in the WT neuronal primary cultures in response to treatment with aSyn PFFs. No calpain activity was measured in the extracellular media of the primary culture (Figures 8a and S12b-c). Interestingly, significant activation of calpains 1 and 2 in neurons was observed 6 hours after the addition of aSyn PFFs (Figure 8b). Calpains are calcium-dependent proteases^102^. Therefore, we next measured calcium homeostasis in PFFs-treated neurons. Calcium imaging using Fura-2 measurement showed that cytosolic calcium started to increase as early as 8 hours after the addition of the PFFs to the neurons (Figure 8d-e) when calpain activity also started to rise significantly (Figure 8b). Interestingly, the increase in calpain activity and calcium level correlated well with the extent of aSyn cleavage and the appearance of the ∼12 kDa fragment, which starts to accumulate within the first 6 hours after the addition of PFF seeds to the neurons (Figure 1j). This suggests that the PFFs entry into neurons results in calpains activation with the cleavage of the PFFs as a consequence. Consistent with this hypothesis, pharmacological inhibition of the enzymatic activity of calpain 1 (by ALLN or PD150606) or calpain 2 (by calpain inhibitor IV) in aSyn KO neurons significantly reduced the truncation level of aSyn PFFs in a concentration-dependent manner (Figure S12d). Calpains 1 and 2 activity (Figure 8b) and calcium increase (Figure 8d-e) were concomitantly upregulated in the soluble fraction of PFFs-treated neurons up to D7, suggesting that these enzymes might also be involved in the processing of aSyn monomers when recruited to the seeds during the early stages of the fibrilization.

**Figure 8.**
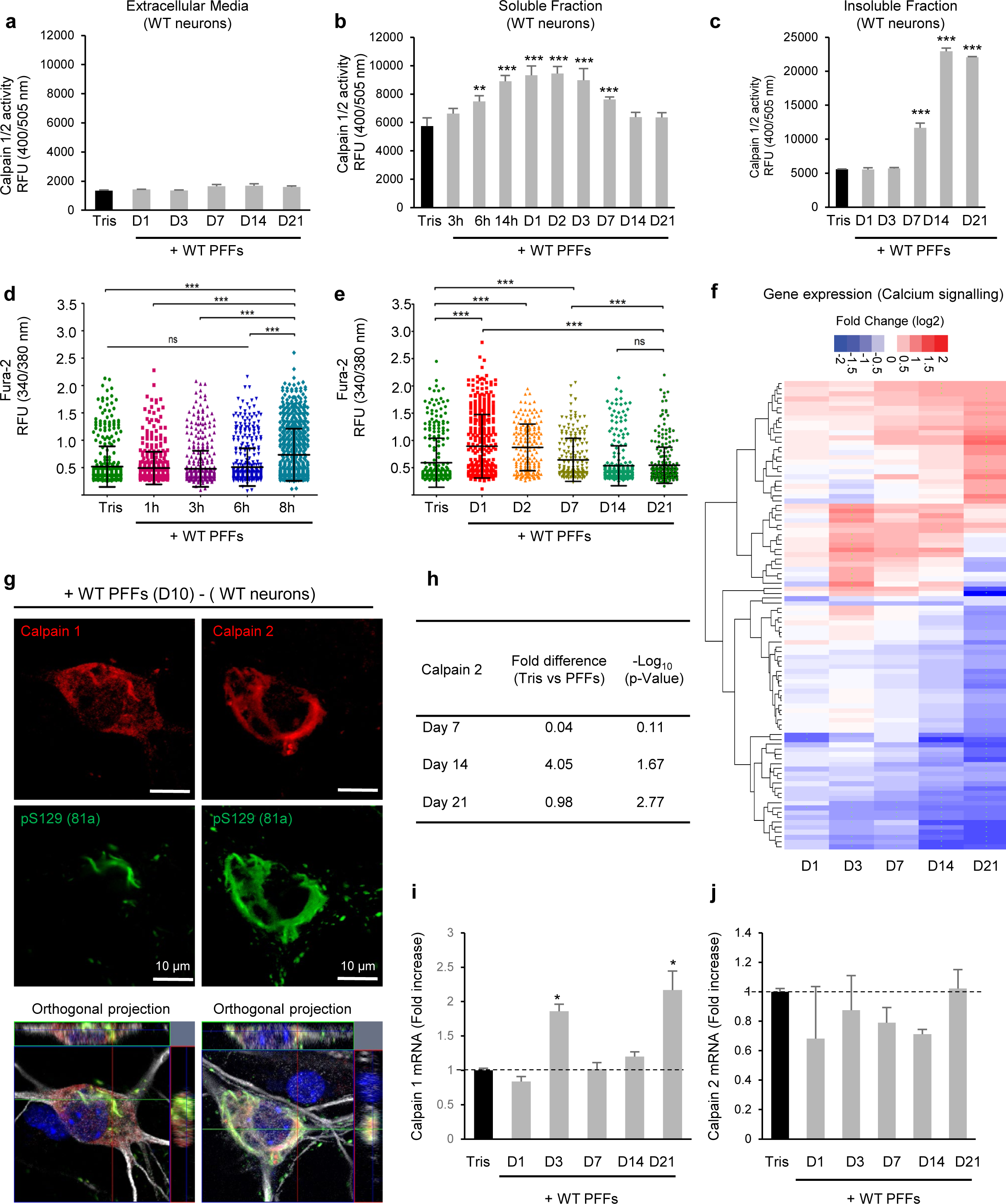
The formation and maturation of the LB-like inclusions induce the activation and the sequestration of calpain 1 and 2. **a-c.** Calpains 1 and 2 are activated in the soluble (**b**) and insoluble fractions (**c**) but not in the extracellular media (**a**) of PFF-treated WT neurons or control neurons (Tris). Activity levels of calpains 1 and 2 were assessed at the indicated times. The graphs represent the mean +/-SD of 3 independent experiments. p<0.001=**, p<0.0001=*** (ANOVA followed by Tukey HSD *post-hoc* test, Tris vs. PFFs-treated neurons). **d-f.** Calcium homeostasis is affected in the PFFs-treated neurons. Intracellular calcium levels were measured in Tris- or PFFs-treated neurons at early time-points (**d**, 1h, 3h, 6h, 8h) or at late time-points (**e**, 1D, 2D, 7D, 14D, 21D) after the addition of the seeds to the neurons. The Fura-2 340/380 ratio was measured in each condition. Each dot represents one neuron subjected for Fura-2 calcium imaging. The data are represented mean+/-SD, *p < 0.05, **p < 0.005, ***p < 0.0005 (ANOVA followed by Tukey HSD *post-hoc* test). **f.** Temporal transcriptomic analysis of the gene expression level in PFFs-treated neurons. Differentially expressed genes were plotted against –log_10_ P-value (t-test, PBS-treated neurons vs. PFFs-treated neurons). Log2 fold changes of the calcium genes expression levels are represented over time. “+” and “-” indicate a significant upregulation or downregulation in the gene expression level. **g.** Confocal imaging confirmed the recruitment of calpains 1 and 2 inside the LB-like inclusions positively stained for pS129-aSyn. Neurons were counterstained with MAP2 antibody and the the nucleus with DAPI staining. Scale bar = 10 μm. **h.** Temporal proteomic analyses showing the enrichment of calpain 2 protein in the insoluble fraction of PFF-treated WT neurons. **i-j.** Transcriptional regulation calpain 1 (**i**) and calpain 2 (**j**) in WT neurons were treated with PFFs for up to 21 days. Quantitative RT-qPCR was performed with primers specific for calpain 1 (**i**) and calpain 2 (**j**) at the indicated time-points after adding PFFs to WT neurons. Results are presented as fold increases in comparison to the respective level in control neurons treated with Tris buffer. mRNA levels were normalized to the relative transcriptional levels of GAPDH and actin housekeeping genes. The graphs represent the mean +/-SD of three independent experiments. *p<0.05 (ANOVA followed by Tukey HSD *post-hoc* test, Tris vs. PFFs-treated neurons).

We next determined whether the cleavage of aSyn mediated by calpains 1 and 2 was only restricted to the cleavage of soluble monomeric aSyn or whether it could also be involved in the truncation of aSyn seeds and/or the newly formed fibrils present in the insoluble fraction. Remarkably, according to WB, a significant increase in calpains 1 and 2 activity was first observed on day 7 in the insoluble fraction of the PFFs-treated neurons, which coincided with the formation of LB-like inclusions in the neurites and cell bodies, as well as the first appearance of HMW aSyn species (Figure 1f). Furthermore, the enzymatic activity of calpains 1 and 2 greatly increased at D14 and D21 post-treatment (Figure 8c). However, no change in calcium level was observed at D14 and D21, suggesting that the activation of the calpains in the insoluble fraction at late stages might result from local calcium increase inside the LB-like inclusions that Fura-2 cannot measure. However, the expression of genes related to calcium signaling pathways was disturbed in PFFs-treated neurons in a time-dependent manner, with most of the significant changes quantified at D14 and D21 (Figure 8f). Thus, altogether our results suggest that adding PFFs to the neurons induces a tight regulation of the calcium homeostasis pathways with the activation of calpains as a consequence and the processing of newly formed aSyn aggregates as an endpoint.

Consistent with this hypothesis, calpains 1 and 2 were detected inside the LB-like inclusions, and their signals perfectly colocalized with pS129 immunoreactivity (Figure 8g). The significant enrichment of calpain 2 level in the seeded-aggregates fraction was confirmed by LC-MS/MS analysis (proteomics dataset, accession no. PXD016850, ProteomeXchange)^58^ (Figure 8h). Finally, gene expression profiling showed that calpain 1 but not calpain 2 mRNA level was markedly upregulated in the early (D3) and late (D21) stages of LB formation and maturation (Figure 8i-j). Altogether, our data support the hypothesis that the calpains play an active role in the remodeling of aSyn fibrils and the formation or maturation of LBs.

### Inhibiting calpain 1 activity accelerates the packaging of the fibrils and their sequestration into small and rounded inclusions highly toxic to the neurons

Having established that calpains play important roles in regulating C-terminal cleavage of aSyn fibrils, we then investigated the extent to which modulating the activity of calpains influences the conversion of aSyn fibrils into inclusions. Calpain inhibitor I or DMSO (negative control) was added to PFFs-treated neurons at D7 (when the newly formed filaments are not yet cleaved^58^ and S12d-e) until D10 (Figure 9a). In PFFs-treated neurons incubated with DMSO, the newly formed fibrils were organized as long and/or compact filamentous structures^54,58,73,103–105^ (Figure 9b-c). Conversely, in the presence of the calpain I inhibitor, the morphology of the seeded aggregates was significantly remodeled into small and numerous rounded inclusions at D10 (Figure 9b-c). This contrasts with the PFFs-treated neurons, where a single large LB-like inclusion is formed overtime. Our data suggest that inhibition of calpain activation could accelerate the conversion of the newly formed fibrils into LB-like inclusions, which are usually observed only after 21 days in PFF-seeded neurons.

**Figure 9.**
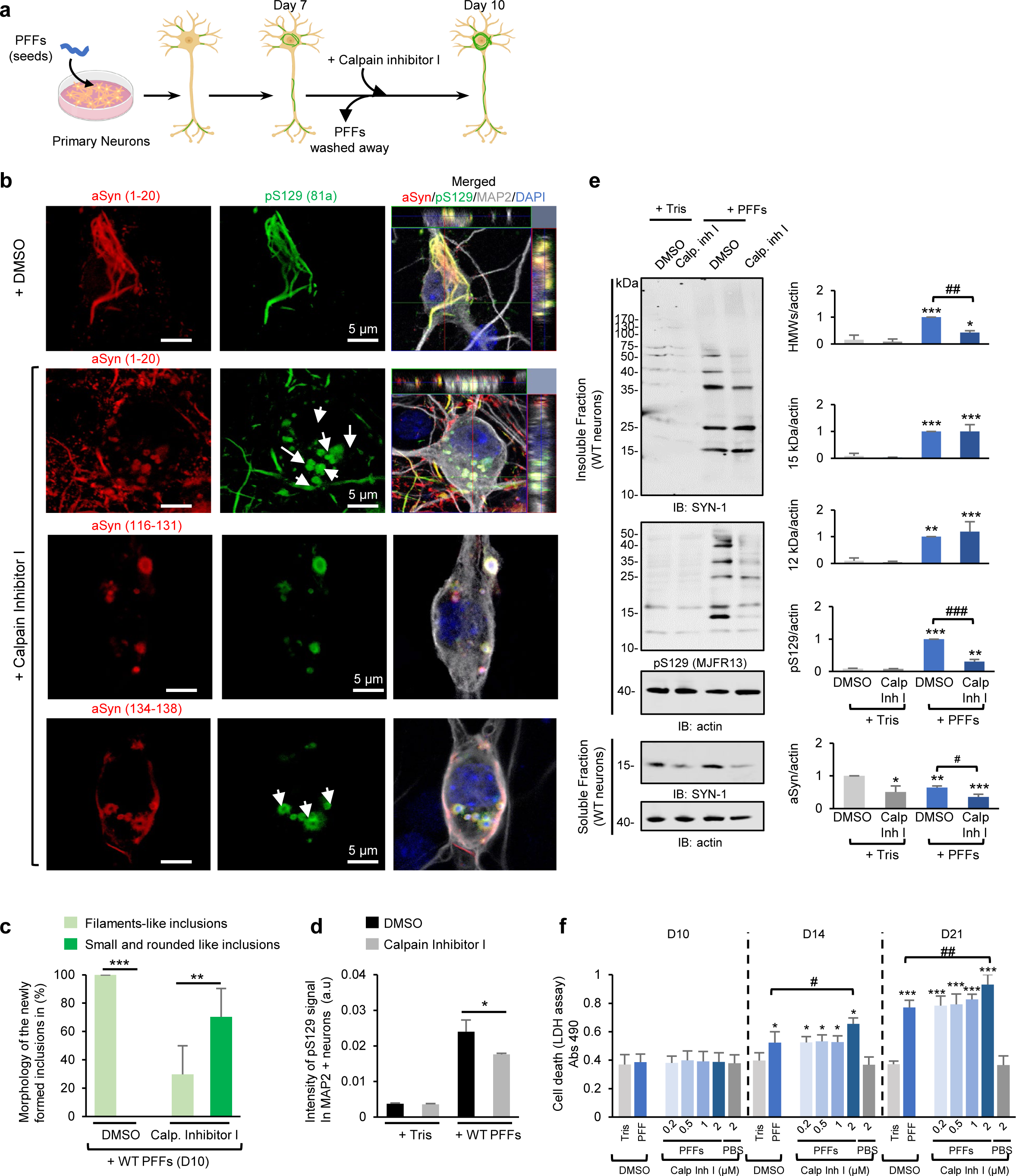
Calpain 1 is involved in the truncation of aSyn in primary neurons. **a.** Experimental design to study the role of calpain 1 in the maturation of the newly formed fibrils. DMSO or calpain inhibitor I was added to PFFs-treated neurons at D7. **b-d.** Level and morphology of the pS129-positive inclusions formed in PFF-treated WT neurons in the presence of DMSO or calpain inhibitor I were assessed after immunostaining using pS129 (81a) antibody in combination with total aSyn N-terminal (1-20) or C-terminal (134-138) antibodies. Neurons were counterstained with MAP2 antibody and the nucleus with DAPI staining. Each experiment was reproduced at least 3 times independently. **b.** Representative images by confocal imaging. Scale bars = 5 μm. **c.** Newly formed aggregates were classified based on their shapes in 2 groups: long filamentous inclusions (**b**, top panel) or small and rounded inclusions (**b**, bottom panel). A minimum of 60 neurons was counted for each condition. **d**. Quantification of images acquired by a high-throughput wide-field cell imaging system. For each independent experiment, duplicated wells were acquired per condition, and nine fields of view were imaged for each well. Each experiment was reproduced at least 3 times independently. Images were then analyzed using the Cell Profile software to identify and quantify the level of LB-like inclusions (stained with pS129 antibody, 81a clone) formed in neurons (MAP2-positive cells). **e.** WB analyses of the insoluble fraction of Tris- or PFFs-treated WT neurons treated with DMSO or calpain inhibitor I between day 7 and day 10. After sequential extractions of the soluble and insoluble fractions, cell lysates were analyzed by immunoblotting. Total aSyn, pS129 and actin were respectively detected by SYN-1, pS129 (MJFR13) and actin antibodies. pS129-aSyn level was estimated by measuring the WB band intensity and normalized to the relative protein levels of actin. **f.** Cell death level was assessed overtime based on lactate dehydrogenase (LDH) release. **c-e, f.** The graphs represent the mean +/- SD of 3 independent experiments. *p<0.05, **p<0.005, ***p<0.0005 (ANOVA followed by Tukey HSD *post-hoc* test, DMSO-Tris-treated neurons vs. the other conditions). ^#^p<0.05, ^##^p<0.005, ^###^p<0.0005 (ANOVA followed by Tukey HSD *post-hoc* test, DMSO-PFFs-treated neurons vs. Calpain inhibitor I-PFFs-treated neurons).

Further characterization of these inclusions using a panel of C-terminal antibodies in combination with pS129 immunostaining confirmed that the C-terminal truncation of the newly formed fibrils was prevented upon treatment with the calpain inhibitor I. This was evidenced by using the C-terminal antibody raised against the 108-120 region that allowed the uncovering of the aSyn-positive aggregates composed of C-terminally cleaved aSyn fibrils that were not pS129 positive (Figure 5e). Conversely, in calpain inhibitor I-treated neurons, all the small and rounded pS129-positive inclusions were positively stained by the aSyn antibody (epitope: 108-120), suggesting that these inclusions are mainly composed of full-length aSyn (Figure 9b). In line with these data, all the pS129-positive inclusions were also positively stained with the C-terminal antibody (epitope: 134-138) (Figure 9b, bottom panel). High-content imaging-based quantification (Figure 9d) and WB analyses (Figure 9e) showed that the pS129 level was significantly lower in the PFF-seeded neurons treated with calpain Inhibitor I. Finally, in the presence of the Calpain inhibitor I, the precocious formation of the round inclusions at D10 induced higher cell death levels in these neurons at D14 and D21 than in those treated with DMSO (Figure 9f). Altogether, our results demonstrate that calpains are key regulators of post-fibrillization C-terminal cleavage of aSyn newly formed and contribute to their efficient packaging and sequestration into LB-like inclusions.

### Implications of post-fibrilization C-terminal truncation for investigating aSyn pathology formation and pathological diversity in synucleinopathies

Although several studies have demonstrated the presence of truncated aSyn aggregates in the brains of patients with PD^2,4,6,22,24,62,84,85^ and DLB^2,4,6,22,23,85,106^, very few studies have explored the role of C-terminal cleavage in MSA^2,6^. To address this knowledge gap, we analyzed tissue homogenates from postmortem brains of MSA patients (Figures 10a and S13a) and confirmed the presence of C-terminally truncated aSyn species, which are detectable with an N-terminal antibody targeting residues 1-20 or a human-specific antibody (clone 4B12) targeting residues 103-108, but not an antibody against residues 134-138. In addition, we stained serial midbrain sections of PD with dementia (PDD) (substantia nigra pars compacta, Figure 10b and S13b for lower magnification), PD with white matter pathology (Figure S13c) and *SNCA* duplication (tegmentum, Figures 10c and S13c for lower magnification) with antibodies against aSyn phosphorylated at residue S129 or the NAC, the N- or the C-terminal domains of the protein. Strikingly, this set of antibodies revealed different types of aSyn inclusions present in the patients’ midbrain tissues. Specifically, while heavier and more diverse pathology was being revealed by SYN-1 (cytoplasmic punctae and extracellular aSyn as well as frequent LBs and LNs), the antibody raised against the last C-terminal amino acids (134-138) displayed rare cytoplasmic inclusions and neuritic pathology. Altogether, these observations suggest that C-terminal antibodies alone may not capture the diversity of aSyn pathology in human brain tissues, which could explain why C-terminally truncated species have always been viewed as a minor component in aSyn pathological aggregates despite their strong representation in proteomic data and WBs^2,4,6,7,22^ or by imaging^43,45,50,51^ when the appropriate antibodies are used.

**Figure 10.**
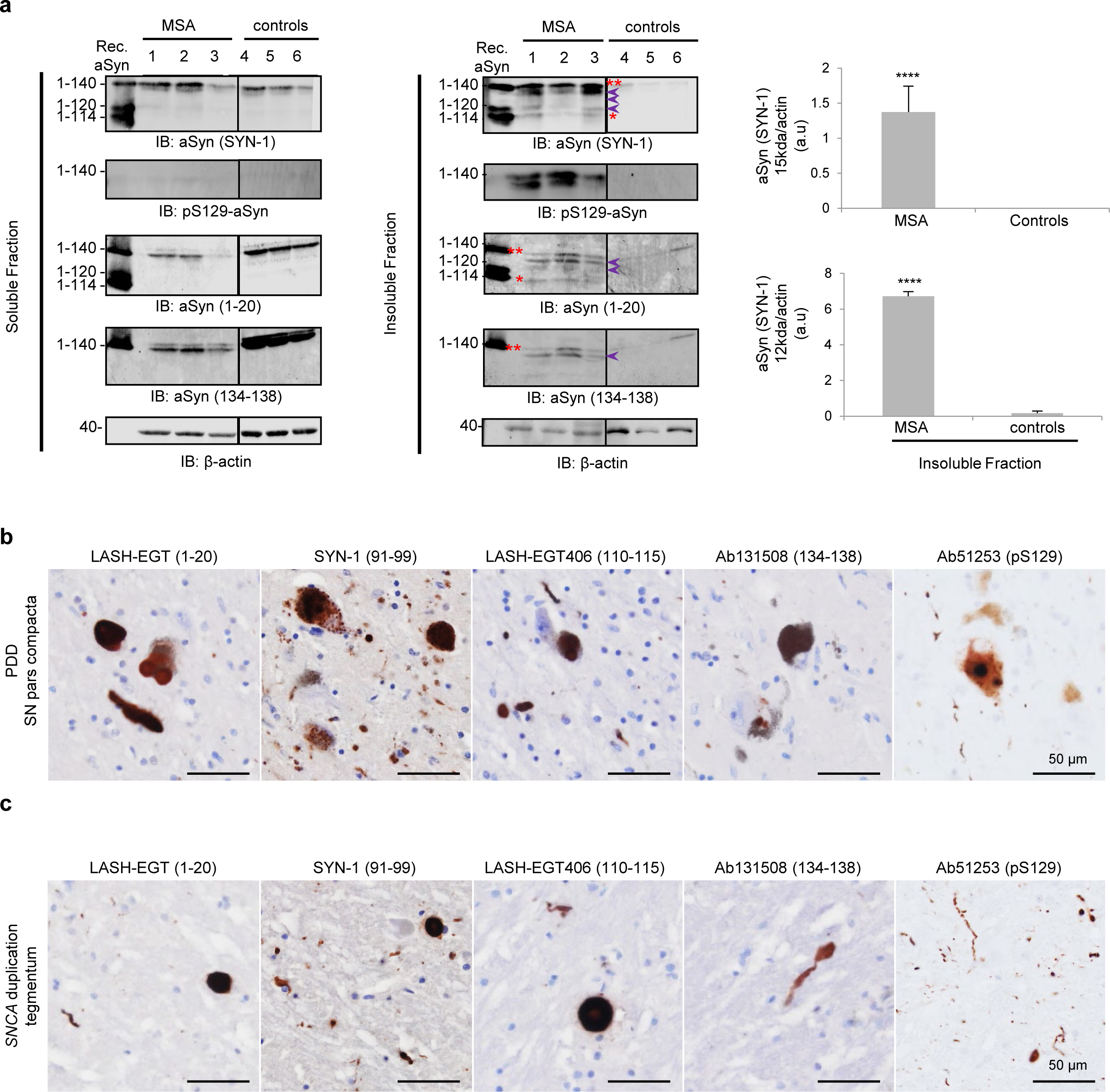
The use of a set of aSyn antibodies captures the morphological spectrum of aSyn pathology in synucleinopathy brain tissues. **a.** WB analyses of the truncation pattern of aSyn in human brain tissue from MSA patients and healthy controls. After sequential extractions of the soluble and insoluble fractions, cell lysates were analyzed by immunoblotting. The levels of total aSyn (1-20, SYN-1 or 134-138 antibodies) or pS129 aSyn were estimated by measuring the WB band intensity and normalized to the relative protein levels of actin. The 15 kDa band is indicated by a double red asterisk, the 12 kDa band by a single red asterisk, and the purple arrows indicate the intermediate aSyn-truncated fragments. **b-c.** Serial sections from the midbrains of PDD (pars compacta) and *SNCA* duplication (tegmentum) cases were stained with aSyn antibodies raised specifically against the N-terminal (epitope: 1-20), the NAC (91-99), the C-terminal (epitopes: 110-115 and 134-138) or pS129 (EP1536Y) regions. Scale bars = 50 μm.

## Discussion

Several aSyn PTMs are consistently associated with LBs and pathological inclusions, suggesting that these modifications are either markers of pathology or play important roles in initiating aSyn misfolding, aggregation, and formation of pathological inclusions inside the neurons. Cryo-EM studies of aSyn fibrils from recombinant proteins and brain-derived fibrils have consistently shown that the C-terminal domain of aSyn remains flexible and accessible to enzyme modifications or cleavage by proteases^107,108^. These observations suggest that post-fibrillization PTMs are likely to occur and influence aSyn fibril structure, interactome and pathogenicity. Consistent with this hypothesis, we recently reported that post-fibrillization nitration induces fibril fragmentation, alters the biochemical surface properties of the fibrils and inhibits aSyn seeding activity^109^. Furthermore, studies from our lab and others have also shown that C-terminal cleavage of aSyn fibrils also occurs post aSyn fibrillization. However, the identity of the enzymes and processes that regulate C-terminal cleavage of fibrils and the role of these modifications in aSyn seeding, LB formation and maturation remain unknown. To address this knowledge gap, we employed a well-established neuronal seeding model that recapitulates all the stages leading to LB formation and neurodegeneration.

First, we showed that the post-fibrilization cleavage of aSyn PFFs is a general phenomenon that occurs within 6-12 hours after their internalization (Figures 1-2) in different models of aSyn pathology formation and spreading, including mammalian cell lines, several types of rodent primary neurons, human iPSC-derived dopaminergic neurons but also *in vivo* in WT and aSyn KO mice (Figure 3). Cleavage of aSyn PFFs has been consistently observed after their internalization into the cells^31,57,58,70^. A recent report^43^ also confirmed our original findings^57^, showing that aSyn PFFs seeds are mainly cleaved at residue 114 (Figures 1 and S4). Next, we investigated the enzymes involved in the cleavage of aSyn PFF upon their internalization in neurons. Our results confirmed previous findings demonstrating that cathepsins^67,70^ and the asparagine endopeptidase^70^ are involved in the processing of the PFFs. Furthermore, we showed that calpains 1 and 2 are also involved in regulating the C-terminal cleavage of both exogenous aSyn PFFs (Figure S12) and newly formed fibrils (Figure 9), suggesting an active role of these enzymes in the processing and/or remodeling of aSyn fibrils and possibly their conversion into LBs.

Interestingly, despite the systematic and rapid proteolytic processing of the PFFs upon internalization into neurons, blocking the C-terminal truncation of the seeds (e.g., PFFs carrying point mutations that prevent their cleavage) did not prevent or significantly alter the seeding and the recruitment of endogenous aSyn or the formation of fibrils and LB-like inclusions in neurons (Figure 4). Thus, C-terminal cleavage of the PFFs is not a prerequisite for the initiation of aSyn seeding and aggregation in neurons. In line with our findings, the addition of C-terminally truncated PFFs did not accelerate nor increase the seeding level in HEK cells (1-110^31^, 1-120^31^ or 1-130^31^), primary neurons (1-114^57^, 1-110^31^, 1-120^31,54^ or 1-130^31^) or PFFs-injected mice (1-110^110,111^, 1-115^112^, 1-119^112^, 1-120^31,113^, 1-122 ^112^, 1-125^112^, 1-129^112^, 1-130^110^) suggesting that the deletion of various aSyn regions other than the NAC domain does not prevent the initiation of the seeding in cells and does not inhibit the formation of the LB-like structures^31,53,54,57^. Only one report suggested that human PFFs lacking the last 30 amino acids have higher seeding activity in SH-SY5Y cells overexpressing mouse WT aSyn^31^. Surprisingly, in this study, the aSyn 1-120 PFFs, shown to have higher seeding activity *in vitro*, had reduced seeding in cells. Our in vitro aggregation results suggest that the high propensity of the C-terminally truncated aSyn species to laterally associate could hinder their uptake and/or reduce the surface of fibrils that catalyze secondary nucleation events, thus resulting in reduced seeding activity. Altogether these reports demonstrate that fibrils generated from truncated monomers or fibrils that undergo post-fibrillization C-terminal cleavage retain their seeding activity in neurons and can induce spreading of the pathology *in vivo*. Furthermore, our results suggest that cleavage of the internalized aSyn PFF seeds may serve other functional or signalling purposes but is not essential for regulating aSyn seeding in neurons.

Interestingly, we observed that the exogenous aSyn PFF seeds and the newly formed aSyn fibrils are processed differently by the neurons and possess distinct biochemical properties, including PTMs (Figure S14). Internalized PFFs do not undergo N-terminal cleavage or phosphorylation on pS129 residue, nor do they colocalize with p62 or ubiquitin, in contrast to the newly formed fibrils showing all these attributes. These observations suggest that C-terminal PTMs may be required to prime N-terminal PTMs or that aSyn PTMs occur during the process of fibril growth and LB formation, and not simply in response to the presence of fibrils. Consistent with this hypothesis, the addition of aSyn PFFs to the aSyn KO neurons does not induce toxicity^58^. These findings, combined with our studies on the internalization of aSyn PFFs in aSyn KO neurons, suggest that the mere presence of fibrils in the absence of the formation of new fibrils by the endogenous protein is not toxic to neurons.

### Post-fibrilization truncation of aSyn impacts the structural and physical properties of the fibrils

It is reasonable to speculate that the PTMs on the surface of aSyn fibrils could significantly influence their interactome, toxicity and maturation into LB-like inclusions ^15,20,109^. In line with this hypothesis, our present work clearly demonstrates that post-fibrillization C-terminal truncation plays an important role in regulating the processing and structural reorganization of the newly formed fibrils from their lateral assembly into higher-order aggregates and to the formation and maturation of LBs^58^ (Figures 6-7). More specifically, we show that the newly formed aSyn fibrils in the seeding model undergo three major changes over time: 1) increased lateral association (Figures 6-7); 2) fragmentation as evidenced by the significant decrease in fibril length over time^58^ resulting in short aSyn filaments randomly organized in the center of the LB-like inclusions at later stages (Figure 6); and 3) loss of C-terminal interacting proteins over time, in particular during the formation of LB-like inclusions (Figure S11). Based on these observations, we hypothesize that aSyn C-terminal truncations could also occur as post-fibrillization events that play important roles in regulating the interactome of the newly formed fibrils and trigger their lateral association, which favors their packaging into LB-like inclusions. Consistent with this hypothesis, site-specific C-terminal photocleavage of full-length PFFs and removal of the last 20 amino acids led to their rapid and tight lateral association. Several studies have also shown that treatment of PFFs with proteases (e.g, calpain 1^114^, trypsin^115^, cathepsin B or D^67,70^), that target the C-terminus of aSyn decreased fibril width^67,114,115^ or height^67,114,115^ and promoted their lateral association^67,114,115^. Interestingly, some of these proteases, such as the calpain 1^52^ and the proteasome^116^, are found sequestered in the *bona fide* LBs in human brain tissues. In addition, the truncated fragments seen upon cleavage of fibrils *in vitro* or in cells are similar to those detected *in vivo*, in the isolated LBs or in the insoluble fractions extracted from _PD2,4,6,22,24,25,33,40-42,52,71,85,117,118, DLB2,4,23,25,71,85,106,117,119, and MSA2,6,57,118,120 patients brain_ tissues. These observations suggest that specific enzymes can cleave the surface-exposed regions of aSyn fibrils, thereby significantly affecting the interactome and structural properties of the fibrils.

### Implications for investigating aSyn pathology formation and pathological diversity in synucleinopathies

The abundance of C-terminally truncated aSyn species within pathological aggregates in the brain has significant implications for the detection, quantification and targeting of aSyn pathologies in the brain and preclinical models of aSyn pathology formation and spreading^121,122^. First, our work underscores the limitation of relying on pS129 antibodies as the primary tools to assess and quantify aSyn aggregates and pathology formation, especially since recent studies have also shown that level and types of C-terminally truncated aSyn varies in aSyn pathologies in different brain regions^47,48^. The use of multiple antibodies targeting pS129 aSyn species and the N- and C-terminal regions of the aSyn is essential to capture the diversity of aSyn pathology in PD brain tissues and to more precisely map the distribution of aSyn species in aSyn pathologies^51,123^ (Figures 10 and S13). This conclusion is supported by recent studies from our group and others using different libraries of aSyn antibodies targeting different aSyn PTMs and epitopes to profile aSyn pathology in different brain regions and synucleinopahties^45,47,48,51,123,124^. Second, we show that C-terminal truncations play a critical role in the aSyn inclusion formation and maturation processes. Third, C-terminal truncations significantly alter the interactome of aSyn fibrils and may influence their clearance, interactions with organelles, and pathogenic properties (Figure S15). Finally, C-terminus targeting therapeutic antibodies may not engage all aSyn pathological aggregates, in particular astrocytic aSyn aggregates, which lack the C-terminal domain of the protein. That being said, it should be emphasized that truncated aSyn species are always found associated with full-length phosphorylated aSyn, which is consistent with post-fibrillization cleavage as the major driver for the generation of these species. The only aSyn inclusions that are composed primarily of C-terminally truncated aSyn species are astrocytic aSyn inclusions, although these inclusions appear to be non-fibrillar and are not positive for any known LB markers^51,123^. Further studies using antibodies that are specific for the different truncated forms of aSyn^51,110,123^ are essential to determine the true abundance and distribution of aSyn truncated aggregates in different brain regions and whether they are secreted and play important roles in aSyn pathology spreading. These studies are essential to determine whether inhibiting or promoting aSyn C-terminal cleavage at specific stages of LB formation represents a viable therapeutic strategy for treating PD and synucleinopathies (Figure S15).

### Conclusions

Although several studies have shown that the C-terminal cleavage of aSyn occurs under both physiological^23,24^ and pathological^4,6,22,25^ conditions, studies in cell-free systems^26–32^ and in cellular and animal models^25,29–39^ of synucleinopathies have consistently shown that C-terminal truncations of aSyn monomers accelerate aSyn aggregation and pathology formation. Previous studies have consistently shown that C-terminal truncations at the monomer level accelerate aSyn fibrillization and could contribute to initiating aSyn fibrillization in the brain. Although these findings suggest that inhibiting this process could protect against aSyn toxicity, it remains unclear whether C-terminal cleavage is the dominant mechanism for triggering aSyn aggregation, as several other biochemical and cellular processes have also been implicated. The work presented here demonstrates that C-terminal truncations are not limited to monomeric aSyn but can also occur at the level of the fibrils and provides strong evidence that post-fibrillization C-terminal cleavages act as master regulators of fibril processing, PTMs, morphology (lateral association) and the formation and maturation of the LBs. Further studies are required to dissect the role and impact of C-terminal cleavages coupled with other post-fibrillization PTMs on aSyn pathology formation, spreading, and neuronal loss in different brain regions. This work, along with recent studies from our group, highlights the crucial importance of focusing on the pattern of PTMs on aSyn fibrils. These findings demonstrate that it is the specific pattern of PTMs, rather than merely the presence of the fibrils themselves, that plays key roles in determining their pathogenicity. Finally, this is also crucial as many of the current diagnostic immunoassays^125^ and therapeutic strategies targeting aSyn pathological aggregates are based on targeting the C-terminal domain of the protein which harbors the majority of disease-associated PTMs, which also influence aSyn fibril dynamics, structure and interactome. Thus underscoring the importance of accounting for PTMs in the development of aSyn diagnostic and therapeutics aimed at targeting the diversity of aSyn pathology

### Limitation of this study

One major limitation of this study is the use of unmodified fibrils derived from recombinant proteins, whereas aSyn fibrils formed in neurons are extensively modified at multiple N- and C-terminal residues. It is possible that these modifications could alter the kinetics and extent of aSyn uptake, processing, clearance and seeding activity. Recent studies by Marotta et al.^126^ showed that aSyn fibrils derived from PDD and AD brains, when added to neuronal cultures, result in predominantly cell body pathology, whereas unmodified recombinant aSyn fibrils give rise to predominantly neuritic pathology. These observations underscore the role of PTMs and other Lewy pathology-associated cofactors in aSyn seeding, pathology formation, and spreading. Therefore, future studies should systematically examine how PTMs found in aSyn aggregates isolated from different synucleinopathies brains influence aSyn uptake, seeding, and LB formation and maturation. Given the challenges in accessing brain tissues and the small amount of material that could be isolated from postmortem human brains, we propose the use of neuronal models of seeding as an alternative source of native modified aSyn fibrils, since fibrils produced in these models recapitulate the PTM pattern found in LBs.

## Materials and Methods

### Antibodies and compounds

Information and RRID of the primary antibodies used in this study are listed in Figure S2. Tables include their clone names, catalogue numbers, vendors and respective epitopes.

### Expression and purification of human and mouse aSyn

pT7-7 plasmids were used for the expression of recombinant human and mouse aSyn in *E.coli*. Human and mouse wild type (WT) aSyn or mouse aSyn mutants (1-135, 1-133, 1-124, 1-120, 1-115, 1-114, 1-111 or 1-101, E114A, D115A, Δ111-115, Δ120-125, Δ133-135, Δ111-115Δ133-135) were expressed and purified using anion exchange chromatography (AEC) followed by reverse-phase High-Performance Liquid Chromatography (HPLC). Recombinant aSyn proteins were fully characterized as described previously by Fauvet et al^11^. For aSyn 1-114, the AEC purification was replaced by a cation exchange chromatography due to the lack of the negatively charged C-terminus. The protein was then further purified and characterized similarly to the other proteins.

### Preparation of WT, deletion mutants and C-terminally truncated PFFs for in vitro studies

Lyophilized full-length, C-terminally truncated (1-135, 1-133, 1-124, 1-120, 1-115, 1-114 or 1-111) single residue deletion (Δ111-115, Δ120-125 or Δ133-135) or double residue deletion (Δ111-115Δ133-135) aSyn proteins were dissolved in 50 mM Tris, 150 mM NaCl. The pH was re-adjusted to 7.5 using a 1M sodium hydroxide solution, and the proteins were subsequently filtered through 100 kDa MW-cut-off filters. The concentration of aSyn in solution was determined using its UV absorption at 280 nm, while, due to the low number of remaining tyrosine residues, the concentration of the C-terminally truncated aSyn mutants and aSyn deleted mutants were determined using the Pierce^TM^ bicinchoninic acid (BCA) Protein Assay kit (Thermo Scientific). Fibril formation was induced by incubating the proteins at a final concentration of 20 µM at 37°C and pH 7.5 (unless stated otherwise) in 50 mM Tris and 150 mM NaCl under constant agitation at 1000 rpm on an orbital shaker for 6 days.

### Preparation of aSyn PFFs for cell treatments

Monomeric human or mouse WT, or biotinylated aSyn monomers were dissolved in 500 μL of Tris buffer (50 mM Tris, 150 mM NaCl, pH 7.5) filtered using a 100 kDa filter (Millipore, Switzerland) at a final concentration of 1-2 mg/mL. Proteins were incubated under constant orbital agitation at 1000 rpm (PEQLAB Biotechnologie GMBH) at 37°C for 5 days^127^ leading to the formation of aSyn fibrils. These fibrils were sonicated with a fine probe 4 times for 5 sec at an amplitude of 20%, (Sonic Vibra Cell, Blanc Labo, Switzerland). Their biophysical and structural properties were then assessed by EM, ThT fluorescence, SDS-PAGE and Coomassie blue staining^128^. For long-term storage, sonicated PFFs were snap-frozen in liquid nitrogen and kept at -80°C.

### Semisynthesis, aggregation and photolysis of aSyn photocleavable at residue 120

The semisynthesis of aSyn photocleavable at residue 120 (aSyn D121Anp) was carried out as previously described^14^. Briefly, we first dissolved 1-106-SR into 6M guanidine and 200 mM Na_2_PO4 supplemented with 30 mM TCEP buffer. Then 1.5 equivalent of A107C-140 D121Anp peptides was added to the reaction mixture. Next, the native chemical ligation was performed with shaking at 1000 rpm at 37°C for 3 hours. When the ligation was complete, as monitored by mass spectroscopy and SDS-PAGE, the protein was purified as previously described^14^. To prepare monomeric aSyn D121Anp, 100 µg of the protein was dissolved in phosphate buffered saline (PBS). For the preparation of fibrils, 500 µg of the protein was dissolved in 100 µL of PBS, and the fibrils were formed by incubating the protein under constant orbital agitation at 1000 rpm (Peqlab, Thriller, Germany) at 37°C for 5 days^127^. The photocleavage of both the fibrils and monomer was performed using a Panasonic UP50, 200 W mercury Xenon lamp, bandpass filter 300-410 lamp with 100% power.

### Electron microscopy (EM)

3.5 μl of sonicated or unsonicated fibrils (final concentration 10 μM) were applied onto glow-discharged formvar/carbon-coated 200-mesh copper grids (Electron Microscopy Sciences, United Kingdom) for 1 min. The grids were blotted off before the two washes with ultrapure water. The grids were then incubated with a staining solution of uranyl formate 0.7% (w/V) for 30 sec. The grids were blotted and dried. Samples were imaged using a Tecnai Spirit BioTWIN operated at 80 kV and equipped with a LaB6 filament and a 4K x 4K FEI Eagle CCD camera.

### Primary culture of hippocampal neurons and treatment with mouse aSyn fibrils

Primary hippocampal neurons were prepared from P0 pups from WT mice (C57BL/6JRj, Janvier) or aSyn KO mice (C57BL/6J OlaHsd, Envigo) and cultured as previously described^58^. The neurons were seeded in 6 wells plates or onto coverslips (CS) (VWR, Switzerland) previously coated with poly-L-lysine 0.1% w/v in water (Brunschwig, Switzerland) at a density of 300,000 cells/mL. After 5 days in culture, the WT or aSyn KO neurons were treated with extracellular aSyn fibrils to a final concentration of 70 nM as previously described^58^. All procedures were approved by the Swiss Federal Veterinary Office (authorization number VD 3392).

### Cell lysis and WB analyses of primary hippocampal neurons

After aSyn PFFs treatment, primary hippocampal neurons were lysed as previously described^54,58,59^. Briefly, treated neurons were lysed in 1% Triton X-100/ Tris-buffered saline (TBS) (50 mM Tris, 150 mM NaCl, pH 7.5) supplemented with protease inhibitor cocktail (Roche, Switzerland), 1 mM phenylmethane sulfonyl fluoride (PMSF) and phosphatase inhibitor cocktail 2 and 3 (Sigma-Aldrich, Switzerland). Cell lysates were sonicated 10 times with a fine probe for 0.5 sec pulse at an amplitude of 20% (Sonic Vibra Cell, Blanc Labo, Switzerland), and then incubated for 30 mins on ice. The cell lysates were then centrifuged at 100,000 g for 30 min at 4°C. The supernatant (soluble fraction) was collected while the pellet was resuspended in 1% Triton X-100/TBS. The pellet was then sonicated as described above and centrifuged for 30 min at 100,000 g. The supernatant was discarded, whereas the pellet (insoluble fraction) was resuspended in 2% SDS/TBS supplemented with protease inhibitor cocktail (Roche, Switzerland), 1 mM PMSF and phosphatase inhibitor cocktail 2 and 3 (Sigma-Aldrich, Switzerland). The insoluble fraction was then sonicated using a fine probe 15 times at 0.5 sec pulse at an amplitude of 20%. The protein concentration of the soluble and insoluble fractions was quantified using the BCA protein assay. Laemmli buffer (4% SDS, 40% glycerol, 0.05% bromophenol blue, 0.252 M Tris-HCl pH 6.8 and 5% β-mercaptoethanol) was then added to the soluble and insoluble fractions.

### Microinjection of PFFs in primary hippocampal neurons

Primary aSyn KO hippocampal neurons were isolated and cultured according to the methods described above. Mouse Alexa Fluor^488^ labeled aSyn PFFs were prepared as described in the methods above. aSyn PFFs were diluted to 9.2 µM (0.13 µg/µl) in distilled H_2_O. Microinjection was performed using INJECT+MATIC Sàrl injector (Geneva, Switzerland) under Leica light transmission microscope. A pre-pulled glass microinjection needle was loaded with 2 µl of fibrils, and fibrils were injected at 2 times units/1 pressure unit/neuron (approximately 6.5 ng in 50 nl/neuron). Cells were incubated for 24 hours post-injection. Cells were fixed using 4% formaldehyde solution (Sigma-Aldrich, Switzerland) for 20 min, washed twice in 1X PBS, then permeabilized and blocked in 1% saponin and 5% bovine serum albumin (BSA) in PBS with 0.01% NaN_3_ for 4 hours at room temperature (RT), then stained with specific anti-aSyn antibodies (epitopes: 1-20 or 134-138) and anti-MAP2 overnight at 4 °C. Cells were washed twice in PBS and incubated in secondary antibody and DAPI stain to visualize nuclei for 2 hours at RT. Secondary antibodies coupled with Alexa Fluor dyes (Thermo Fisher Scientific, USA) were applied for 1 hour at RT. Cells were washed twice in PBS, and coverslips were mounted onto microscopy slides using Mowiol mounting medium (Sigma-Aldrich, Switzerland). The cells were examined with a confocal laser-scanning microscope (LSM 700, Carl Zeiss Microscopy, Germany) with a 40x objective and analyzed using Zen software (RRID:SCR_013672).

### Immunocytochemistry (ICC)

HeLa cells or primary hippocampal neurons treated with PFFs were washed twice with PBS, before being fixed in 4% PFA for 20 min at RT. Cells were then immunostained as previously described^58^. The antibodies used are indicated in the corresponding legend section of each figure. The source and dilution of each antibody can be found in Figure S2. The cells were examined with a confocal laser-scanning microscope (LSM 700, Carl Zeiss Microscopy, Germany) with a 40x objective and analyzed using Zen software (RRID:SCR_013672).

### Quantitative high-throughput wide-field cell imaging screening (HTS)

After aSyn PFFs treatment, primary hippocampal neurons plated in a clear black bottom 96 well plates (BD, Switzerland) were washed twice with PBS, fixed in 4% PFA for 20 min at RT and then immunostained as described above. Images were acquired using the Nikon 10X/ 0.45, Plan Apo, CFI/60 of the IN Cell Analyzer 2200 (GE Healthcare, Switzerland), a high-throughput imaging system equipped with a high-resolution 16-bits sCMOS camera (2048×2048 pixels), using a binning of 2×2. For each independent experiment, duplicated wells were acquired per condition, and nine fields of view were imaged for each well. Each experiment was reproduced at least 3 times independently. Images were then analyzed using Cell profiler 3.0.0 software (RRID:SCR_007358) for identifying and quantifying the level of LB-like inclusions (stained with pS129 antibody) formed in neurons MAP2-positive cells as previously described^58^.

### Correlative Light Electron Microscopy (CLEM)

Mouse primary hippocampal neurons were grown on 35 mm dishes with alphanumeric searching grids etched to the bottom glass (MatTek Corporation, Ashland, MA, USA) and treated with WT PFFs. At the indicated time point, cells were fixed for 2 hours with 1% glutaraldehyde and 2.0% PFA in 0.1 M phosphate buffer (PB) at pH 7.4. After washing with PBS, ICC was performed (for more details, see the corresponding section in the Materials and Methods). Neurons with LB-like inclusions (positively stained for pS129) were selected with fluorescence confocal microscope (LSM700, Carl Zeiss Microscopy, Germany) for ultrastructural analysis. The precise position of the selected neuron was recorded using the alpha-numeric grid etched on the dish bottom. The cells were then fixed further with 2.5% glutaraldehyde and 2.0% paraformaldehyde in 0.1 M PB at pH 7.4 for another 2 hours. After washing 5 times for 5 min each with 0.1 M cacodylate buffer at pH 7.4, cells were postfixed with 1% osmium tetroxide in the same buffer for 1 hour and washed with double-distilled water before contrasting with 1% uranyl acetate stain for 1 hour. The cells were then dehydrated in increasing concentrations of alcohol (2 × 50%, 1 × 70%, 1 × 90%, 1 × 95% and 2 × 100%) for 3 min each wash. Dehydrated cells were infiltrated with Durcupan resin diluted with absolute ethanol at 1 : 2 for 30 min, at 1 : 1 for 30 min and 2 : 1 for 30 min and twice with pure Durcupan (Electron Microscopy Sciences, Hatfield, PA, USA) for 30 min each. After 2 hours of incubation in fresh Durcupan resin, the dishes were transferred into a 65°C oven for the resin to polymerize overnight. Once the resin had hardened, the glass CS on the bottom of the dish was removed by repeated immersion in hot (60°C) water, followed by liquid nitrogen. The cell of interest was then located using the alphanumeric coordinates previously recorded, and a razor blade was used to cut this region away from the rest of the resin. This piece was then glued to a resin block with acrylic glue, trimmed with a glass knife using an ultramicrotome (Leica Ultracut UCT, Leica Microsystems), and then ultrathin sections (50–60 nm) were cut serially from the face with a diamond knife (Diatome, Biel, Switzerland) and collected onto 2 mm single-slot copper grids coated with formvar plastic support film. Sections were contrasted with uranyl acetate and lead citrate and imaged with a transmission electron microscope (Tecnai Spirit EM, FEI, The Netherlands) operating at 80 kV acceleration voltage and equipped with a digital camera (FEI Eagle, FEI).

### Identification of N- and C-terminal truncation sites by quantitative proteomic analyses

After aSyn WT PFF treatment, primary hippocampal neurons (treated for 14 or 21 days) were lysed as previously described^54,58,59^ and separated by 16% Tricine gel, which was then stained with Coomassie Safestain (Life Technologies, USA). Each gel lane was entirely sliced, and proteins were in-gel digested as previously described^129^, skipping the reduction and alkylation procedure. Peptides were desalted on stageTips^130^ and dried under a vacuum concentrator. For LC-MS/MS analysis, resuspended peptides were separated by reversed-phase chromatography on a Dionex Ultimate 3000 RSLC nano UPLC system connected in line with an Orbitrap Lumos (Thermo Fisher Scientific, Waltham, USA). Raw data were processed using Mascot (Matrix Science, Boston, USA), MS-Amanda^131^ and SEQUEST in Proteome Discoverer v.1.4 (RRID:SCR_014477) against the Uniprot Mouse protein database. Data were further processed and inspected in Scaffold4 (Scaffold Proteome Software, RRID:SCR_014345), and spectra of interest were manually validated.

### Fura-2 calcium imaging

Primary hippocampal neurons treated with Tris and PFFs at various time points were incubated in growth media with 1 μM Fura-2 AM (Abcam, UK) for 30 minutes, followed by without Fura-2 AM for 30 minutes. After this, fluorescent images were collected by excitation at 340 nm and 380 nm, emission at 510 nm for about 4-5 minutes. At the end of each condition, ionomycin was added as a positive control to verify an increase in intracellular calcium levels. Fura-2 AM calcium imaging analysis was performed using an Olympus cellSens Software (RRID:SCR_014551) ratio analysis tool as per instruction manual to collect Fura-2 340/380 ratio values.

### Intra-striatal stereotaxic injection of mouse and human aSyn PFFs into mice brain

Animal experiments were performed according to the guidelines of the European Directive 2010/63/EU and Belgian legislation. The ethical committee for animal experimentation from UCB Biopharma SPRL (LA1220040 and LA2220363) approved the experimental protocol. All surgical procedures were done with wild-type male mice from C57Bl/6J and male aSyn KO mice (The Jackson Laboratory; #023837). Surgeries were performed under general anesthesia using a mixture of 50 mg/kg of ketamine (Nimatek, Eurovet Animal Health B.V.) and 0.5 mg/kg of medetomidine (Domitor, Orion Corporation) injected intraperitoneally. In addition, 2.5 mg/kg atipamezole (Antisedan, Orion Corporation) was given to support awakening. The recombinant purified full-length human PFFs were thawed and sonicated at RT by probe sonication (Q500 sonicator from Qsonica; 20 kHz; 65% power, for 30 pulses of 1 sec ON, 1 sec OFF). C57BL/6 wild-type and aSyn KO mice were anesthetized and stereotactically injected into the right striatum (coordinates: anterior-posterior, 0.2 mm; mediolateral, -2 mm; dorsoventral, −3.2 mm relative to bregma and dural surface) using a glass capillary attached to a 25 µl Hamilton microsyringe. 5 µg of human PFF (2.5 µg/µl in sterile PBS) were administered at a constant rate of 0.2 µl per minute, and the needle was left in place for an additional 2.5-min period before its slow retraction. Mice were sacrificed at 0, 1, 3 and 7 days post-injection. After anesthesia, the animals were perfused through the ascending aorta with a mixed solution (30 ml/mouse) made of PBS and heparin (10 U/ml, 4°C). Brains were removed from the skull, and the right striatum was dissected out. Brain tissues were snap-frozen in nitrogen and stored at -80°C.

### Cell lysis and WB analyses of human PFFs-injected mice brains and human brain tissues

The soluble and insoluble fractions of mouse and human brain tissues, for biochemistry analyses, were prepared according to previously described protocols^54,59^. Briefly, tissues were homogenized in TBS-T lysis buffer containing 50 mM Tris-HCl, 1% Triton X-100, 150 mM NaCl, and a cocktail of protease and phosphatase inhibitors (Roche, Switzerland), and were sonicated at 20 % power for 20 s using Q500 sonicator (Qsonica, USA). Homogenates were centrifuged at 100,000 g for 30 min at 4 °C. The supernatant corresponded to the soluble fraction. The pellet was resuspended in TBS-T lysis buffer, sonicated at 20% power for 20 s, and then centrifuged at 100,000 g for 30 min at 4°C. The supernatant was discarded. The insoluble fraction was prepared by resuspending the pellet in TBS-T lysis buffer containing 2% SDS, and then sonicated at 20% power for 40 s. Protein concentration was determined using the BCA assay. 20 µg of total protein was loaded in Tris-glycine 16% gels (Novex Thermo, USA) and migrated at 100 V. Gels were transferred onto Immobilon polyvinylidene diffluoride membranes (Merck-Millipore, Germany) at 25 V for 30 min using Trans-Blot Turbo (Bio-Rad Laboratories, USA). Membranes were blocked for 1 hour with Odyssey blocking buffer (LiCor, USA) and then incubated overnight at 4°C with different primary antibodies diluted in the same blocking buffer. After rinsing with TBS-Tween 0.1%, membranes were incubated with IRDye® conjugated secondary antibodies (1:5,000; LiCor, USA) for 1 hour at RT, and visualization was performed by fluorescence using Odyssey CLx from LiCor. Signal intensity was quantified using Image Studio 3.1 from LiCor (RRID:SCR_013715). Primary antibodies were directed to human aSyn epitope 103-108 (4B12, 1:1,000, Thermo FisherScientific, USA), mouse aSyn (D37A6, 1:1,000, Cell Signaling Technology, USA), aSyn epitope 1-20 (1:750, homemade), aSyn epitope 91-99 (clone 42, SYN-1, 1:1,000, Becton Dickinson, USA), aSynuclein epitope 134-138 (1:1,500, Abcam, UK), phospho-serine 129 aSyn (1:1,500, Abcam, UK), actin (1:3,000, Cell Signaling Technologies, USA).

### Human brain samples

Case demographics are presented in Figure S13a. Frozen human brain samples from MSA patients were obtained from the Netherlands Brain Bank (NBB), Netherlands Institute for Neuroscience (Amsterdam, the Netherlands; open access: www.brainbank.nl) after approval of the project by the Netherlands Brain Bank ethical committee. All material was collected from donors for or from whom a written informed consent for brain autopsy and the use of material and clinical information for research purposes had been obtained by the Netherlands Brain Bank. The PDD and *SNCA* duplication cases selected from the Queen Square Brain Bank (QSBB) at the University College London (UCL, UK), Institute of Neurology, were collected following protocols approved by the London Multicentre Research Ethics Committee. Written informed consent was obtained from all donors before brain collection. The samples were stored under a license approved by the Human Tissue Authority (HTA; license number 12198). Ethical approval for the research was granted by the National Research Ethics Service (NRES) Committee London Central.

### Immunohistochemistry on postmortem human tissue

Eight-micrometer-thick paraffin-embedded sections were cut sequentially. For selected primary antibodies (Figure S2a) immunohistochemical staining was performed as described previously^51,123^. Briefly, sections were deparaffinized, treated with 80% formic acid for 10 min and/or citrate buffer at pH 6.0 under pressure for 10 min at 121 °C, and treated with hydrogen peroxide 3%. Blocking was carried out for 30 min in fetal bovine serum (10%). Sections were incubated in primary antibodies overnight at 4 °C (Figure S2) and in secondary antibody-horseradish peroxidase complex (REAL EnVision detection kit, Dako #K5007) for 1 h at RT. Visualisation was carried out with 3,3’-diaminobenzidine (DAB) and counterstaining with hematoxylin. Sections were imaged using Olympus VS120 (RRID:SCR_018411) and analysed on QuPath (RRID:SCR_018257).

### Statistical analyses

The level of total aSyn (15 kDa, 12 kDa or HMW) or pS129-aSyn were estimated by measuring the WB band intensity using Image J software (U.S. National Institutes of Health, Maryland, USA; RRID:SCR_001935) and normalized to the relative protein levels of actin. All the experiments were independently repeated 3 times. The statistical analyses were performed using Student’s *t*-test or ANOVA test followed by Tukey-Kramer *post-hoc* test using KaleidaGraph (RRID:SCR_014980). The data were regarded as statistically significant at p<0.05.

## Acknowledgments

This work was supported by funding from EPFL and UCB (H.A.L), (A.L.M), (F.A), (N.M), (N.A.B), (A.C), (G.L), (S.J) and (S.N). R.W.-M and S.V were supported by the Joint Program for Neurodegenerative Diseases (JPND) from the UK Medical Research Council (R.W.-M.). J.H and C.S were supported by the Multiple System Atrophy Trust; the Multiple System Atrophy Coalition; Fund Sophia, managed by the King Baudouin Foundation; and Karin & Sten Mortstedt CBD Solutions. Queen Square Brain Bank for Neurological Disorders is supported by the Reta Lila Weston Institute for Neurological Studies and the Medical Research Council UK. This research was supported in part by the National Institute for Health Research University College London Hospitals Biomedical Research Centre. J.Y.L and C.H were supported by the NNSF-81430025, Swedish Research Council, EU-JPND (aSynProtec, REfrAME) and EU-ITN (Syndegen).

We acknowledge Elena Gasparotto, Jérémy Campos, Yllza Jasiqi and Lorène Aeschbach for their valuable technical assistance respectively: Elena, Lorène and Yllza for their continuous support with the preparation of the primary culture, Elena for the cloning of the mutated aSyn, Jérémy for the expression and purification of WT or mutants aSyn proteins.

We are grateful to Dr. Arne Seitz and his staff at the Bio-imaging Core Facility (EPFL) for their technical support. We are grateful to the individuals and their families for consenting to donate to QSBB and NBB.

## Author contributions

H.A.L conceived and supervised the study. H.A.L and A.L.M.M designed all the experiments and wrote the paper. A.L.M.M performed and analyzed the experiments shown in Figures 1-2 a-c, 3 a-b, 4, 5, 6, 8 a-c, f-j, 9 and Figures S2, S3, S4 a-f, i-m, S6, S7 b-c, S8, S9 a-b, d, S10, S12 d-e, S14 and S15. M.F.A designed, performed and analyzed the experiments shown in Figures 10 b-c and S1, S4 g-k, S12 b-c and S13. N.M designed, performed and analyzed the experiments shown in Figures S11. N.A.B and A.C designed, performed and analyzed the experiments shown in Figure 7. A.C produced and characterized aSyn-biotin PFFs shown in Figure S1 and used in Figure S11. G.L, S.J and S.N designed, performed and analyzed the experiments shown respectively in Figures 2 d, 8 d-e and S7 a. S.V and R.W.M designed Figures 3 d and S9 e-g. S.V performed and analyzed the experiment shown in Figures 3 d and S9 e-g. R.H performed and analyzed the LC-MS/MS of the experiments shown in Figures 1 I, 5 a-b, S4 and S10 c. J.H and C.S contributed to the analysis of Figures 10b-c and S13 a-c. J.Y.L and C.H performed and analyzed the experiment shown in Figure S13 c. M.C prepared the samples for CLEM analysis and acquired EM images in Figure 6. G.K supervised the experiments shown in Figure 6 and contributed to the interpretation of the data. G.M.C designed and analyzed the experiments shown in Figures 3 c, 10 a and S9 c. L.W performed and analyzed the experiments shown in Figures 3 c, 10 a and S9 c. A.M designed and performed the experiments shown in Figures 3 c. P.D and M.C supervised the study performed in Figures 3 c, 10 a and S9 c. All authors reviewed and contributed to the writing.

