## Supplemental Figures for "Dissecting the differential role of C-terminal truncations in the regulation of aSyn pathology formation and the biogenesis of Lewy bodies"

^1^Laboratory of Molecular and Chemical Biology of Neurodegeneration, Brain Mind Institute, Ecole Polytechnique Fédérale de Lausanne (EPFL), 1015 Lausanne, Switzerland. ^2^Department of Physiology, Anatomy and Genetics, University of Oxford, Oxford, OX1 3QX, UK. ^3^Queen Square Brain Bank for Neurological Disorders, Department of Molecular Neuroscience, UCL Institute of Neurology, London, UK. ^4^Neural Plasticity and Repair Unit, Wallenberg Neuroscience Center, Department of Experimental Medical Science, Lund University, 22184 Lund, Sweden. ^5^Institute of Health Sciences, China Medical University,110122 Shenyang, China. ^6^Proteomic Core Facility and Technology Platform, EPFL, Lausanne 1015, Switzerland. ^7^BioEM Core Facility and Technology Platform, EPFL, Lausanne 1015, Switzerland. ^8^UCB Biopharma SPRL, Chemin du Foriest B-1420 Braine-l'Alleud, Belgium.

\# These authors contributed equally

*To whom correspondence should be addressed: Laboratory of Molecular and Chemical Biology of Neurodegeneration, Brain Mind Institute, Ecole Polytechnique Fédérale de Lausanne, 1015 Lausanne. Tel: +41216939691, Fax: +41216939665,

**Running title:**

C-terminal truncation: a master regulator of LB biogenesis.

**Keywords:** alpha‐synuclein (aSyn), Parkinson’s disease (PD), Lewy body (LB), aggregation, post‐translational modification (PTM), truncation, phosphorylation

**Material and Methods**

***Preparation of fluorescently labelled mouse aSyn PFFs***

aSyn mouse WT PFFs were diluted at a concentration of 250 μM in a final volume of 500 μl of PBS. The pH was adjusted to 7.5. One equivalent of Atto^488^ maleimide (Atto-Tec, Switzerland) was added and incubated at 4 °C overnight. The labelled PFFs were then ultracentrifuged at 100,000 g for 1 hour at 4 °C. The supernatant was collected, and the pellet resuspended in PBS. This wash step was repeated until the dye in excess was removed. PFFs were loaded onto a SDS-PAGE gel and labelling was confirmed by scanning the gel using Typhoon FLA 7000 (GE Healthcare Life Sciences, Switzerland) with respective excitation and an emission wavelength of 400 and 505 nm. Labelled fibrils were then fragmented via sonication for 4 times at 5 sec at an amplitude of 20%, (Sonic Vibra Cell, Blanc Labo, Switzerland). PFFs were then snap-frozen in liquid nitrogen and kept at -80 °C for long-term storage. The biophysical and structural properties of the fibrils were then assessed by EM, SDS-PAGE/ Coomassie staining and thioflavin T assays.

***Preparation of biotinylated aSyn monomers***

Recombinant FL M1C aSyn (10 mg) was placed in 1.0 mL of degassed buffer (200 mM Tris, 6.0 M Gdn at pH 6.5). To make sure that cysteines were in the reduced form, the protein was treated with 1.0 equivalent of Tris(2-carboxyethyl)phosphine (TCEP) and incubated at 37°C for 20 min. The reaction was monitored by LC-MS to confirm that dimers were reduced to monomers. Then 3.0 equivalent of biotin maleimide (10 µL of 200 mM solution in dimethylformamide) was added and stirred at 37°C for 60 min at pH 6.5. The progression of the labeling was monitored by electrospray ionization (ESI) LC/MS, and after complete consumption of the starting M1C aSyn protein, the reaction was quenched by lowering the pH from 6.5 to 5. The reaction mixture was cooled down to 5°C and purified by C8 semi-preparative column chromatography using a 20%-80% acetonitrile gradient over 60 min using 3.0 mL/min flow rate. The purity of the fractions was verified by ultra performance liquid chromatography (UPLC), and the biotin-labeled aSyn was lyophilized. The purity and identity of the final product were confirmed by UPLC, SDS-PAGE and ESI-LC/MS analyses.

**HeLa cell culture, plasmid transfection and treatment with human aSyn PFFs**

HeLa cells were cultured in Dulbecco's modified Eagle's medium (DMEM), high glucose, pyruvate and GlutaMAX™ supplemented with 10% FBS and 10 μg/mL penicillin and streptomycin in a humidified incubator, 5% CO2, 37°C. According to the manufacturer's protocol, HeLa cells were transfected with a pAAV vector coding for human aSyn using Lipofectamine 2000 transfectant reagent (Life Technologies, Switzerland). 24 hours post-transfection, human aSyn fibrils were added to the cells at a final concentration of 500 nM.

***Cell lysis and WB analyses of HeLa cells***

HeLa cells were lysed in radioimmunoprecipitation assay (RIPA) buffer (150 mM sodium chloride, 50 mM Tris pH 8.0, 1% NP-40, 0.5% deoxycholate, 0.1% SDS) supplemented with protease inhibitor cocktail (cOmplete^TM^, Roche, Switzerland), 1 mM PMSF and phosphatase inhibitor cocktail 2 and 3 (Sigma-Aldrich, Switzerland). Cell lysates were cleared by centrifugation at 4°C for 15 min at 13’000 rpm. The pellet (insoluble fraction) was resuspended in SDS/TBS supplemented with protease inhibitor cocktail (cOmplete^TM^, Roche, Switzerland), 1 mM PMSF and phosphatase inhibitor cocktail 2 and 3 (Sigma-Aldrich, Switzerland) and sonicated using a fine probe 15 times at 0.5 sec pulse at an amplitude of 20%. BCA protein assay was performed to quantify the protein concentration in the soluble and insoluble fractions before the addition of Laemmli buffer 4x. Proteins from the soluble and the insoluble fractions were then separated on a 16% Tricine gel, transferred onto a nitrocellulose membrane (Thermo Fisher Scientific, Switzerland) with a semi-dry system (Bio-Rad, UK) and immunostained as previously described^1^.

***iPSC-derived neurons and treatment with human aSyn PFFs***

Induced pluripotent stem cells were derived from dermal fibroblasts. The line used in this study stems from a healthy control female of 78 years old and was reprogrammed using the CytoTune-iPS Sendai Reprogramming kit (Invitrogen). The iPSC line has been characterized in previous studies as SFC856-03-04 (control 4)^2^^,^^3^. The iPSCs were kept at 37ºC, 5% CO_2_ in feeder-free culture conditions with daily changes of mTeSR™1 (StemCell Technologies, UK), on hESC-qualified Matrigel-coated plates (BD biosciences, UK) and passaged as single cells using TrypLE Express incubation (Life Technologies, UK) and ROCK inhibitor (10 µM Y-27632) (Bio-Techne, Tocris, UK). IPSCs were differentiated into dopaminergic neurons using a floor-plate-based culture established by Kriks *et al.*^4^ and modified as previously described^5^^,^^6^. The cells were seeded in 6-well plates coated with Geltrex (Life Technologies, UK) and grown to confluency. Patterning was achieved using the medium with differentiation and neurotrophic factors, as listed below. The medium was fully changed every two days, with half changing every other day until the day *in vitro* 20 (DIV20). The cells were then dissociated with StemPro Accutase (Life Technologies, UK) and re-plated onto Geltrex-coated 12 well plates (Corning, UK) in an even monolayer of 3x10^5^ cells/cm^2^ in medium containing ROCK inhibitor (10 µM Y-27632). Cultures were treated with 1 µg/ml mitomycin C (Bio-Techne, Tocris, UK) in neurobasal (NB) medium for 1 hour to remove proliferating cells and washed with NB medium before adding fresh NB medium. After a full medium change 3 days later to remove dead cells, medium was half changed every 2-3 days until DIV50 when 70 nM of PFFs were added to neurons 14, 10, 7, 3 and 1 days before fractionation analyses at DIV64. For WB analyses, cultures were washed in phosphate-buffered saline (PBS), detached by scraping into fresh 1% Triton X-100/ Tris-buffered saline (TBS) (50 mM Tris, 150 mM NaCl, pH 7.5) with protease (cOmplete^TM^, Roche, Switzerland) and phosphatase inhibitors (PhosSTOP^TM^, Roche, Switzerland) and subjected to a previously described fractionation protocol^7^. After sonication using a fine probe, 0.5-sec pulse at an amplitude of 20%, 10 times (UP50H, Hielscher, Germany), cell lysates were incubated on ice for 30 min and centrifuged (Optima TLX ultracentrifuge, TLA-110 rotor, Beckman Coulter, USA) at 100,000 g for 30 min at 4°C and the supernatant (soluble fraction) collected. The pellet (insoluble fraction) was resuspended in 2% sodium dodecyl sulfate (SDS)/TBS supplemented with protease (cOmplete^TM^, Roche, Switzerland) and phosphatase inhibitors (PhosSTOP^TM^, Roche, Switzerland) and sonicated using a fine probe (0.5-sec pulse at an amplitude of 20%, 15 times). 10 ug of protein from each fraction was loaded on Criterion™ TGX™ precast gel 4-15% (BioRad) and transferred onto Trans-Blot® Turbo™ polyvinylidene difluoride (PVDF) membranes (BioRad). Membranes were blocked in 5% milk in PBS containing 0.1% Tween for 30 min and incubated overnight in blocking solution with a total aSyn antibody (SYN-1, BD biosciences; 1:500). The secondary anti-mouse HRP (BioRad; 1:5000) and loading control β-actin-HRP (Abcam, 1:50 000) were diluted in blocking solution and applied for 1 hour at room temperature (RT). The membrane was exposed to Immobilon Western Chemiluminescent HRP Substrate and developed with the ChemiDoc^TM^ System (BioRad). Protein levels were measured with the Image Lab Software (RRID:SCR_014210) (BioRad) and analyzed with GraphPad Prism (RRID:SCR_002798).

***Medium and reagents used during differentiation of iPSC cells into dopaminergic neurons***

| **Medium** | **Product name** | **Supplier** | **Catalogue no** | **Working** | **Time added** |
| --- | --- | --- | --- | --- | --- |
| **KO DMEM KSR** | Knockout DMEM | Life Technologies | 10829018 | n/a | 100% DIV 0-4  75% DIV 5-6  50% DIV 7-8  25% DIV 9-10 |
|  | KnockOut Serum  Replacement | Life Technologies | 10828010 | n/a |  |
|  | L-Glutamine | Life Technologies | 25030-024 | 2mM |  |
|  | MEM NEAA | Life Technologies | 11140-050 | 1X |  |
|  | β-mercaptoethanol | Gibco | 21985023 | 10µM |  |
| **NNB** | Neurobasal Medium | Life Technologies | 21103049 | n/a | 25% DIV 5-6  50% DIV 7-8  75% DIV 9-10 |
|  | N2 Supplement | Life Technologies | 17502048 | 0.5X |  |
|  | B-27® Supplement  w/o Vit A | Life Technologies | 12587010 | 0.5X |  |
|  | L-Glutamine | Life Technologies | 25030-024 | 2mM |  |
| **NB** | Neurobasal Medium | Life Technologies | 21103049 | n/a | DIV11- |
|  | B-27® Supplement  w/o Vit A | Life Technologies | 12587010 | 1X |  |
|  | L-Glutamine | Life Technologies | 25030-024 | 2mM |  |
| **Growth factors** | SB431542 | Bio-techne (Tocris) | 1614 | 10µM | DIV0-4 |
|  | LDN193189 | Sigma | SML0559-5MG | 100nM | DIV0-10 |
|  | Sonic Hedgehog C24II  high activity | Bio-techne (Tocris) | 1845-SH-500 | 100ng/mL | DIV1-6 |
|  | Purmorphamine | Bio-techne (Tocris) | 4551/10 | 2µM | DIV1-6 |
|  | FGF8a | Stratech | 16124-HNAE-SIB | 100ng/µL | DIV1-6 |
|  | CHIR99021 | Bio-techne (Tocris) | 4423 | 3µM | DIV3-12 |
|  | TGFβ3 | Peprotech | 100-36E | 1ng/mL | DIV13- |
|  | DAPT | Abcam | ab120633 | 10µM | DIV13- |
|  | db-cAMP | Sigma | D0627-1g | 0.5mM | DIV13- |
|  | GDNF | Peprotech | 450-10 | 20ng/µL | DIV13- |
|  | BDNF | Peprotech | 450-02 | 20ng/µL | DIV13- |
|  | Ascorbic acid | Sigma | A4544-25G | 0.2mM | DIV13- |

***Cathepsin B, D and L*** ***activity assay in primary neurons***

Cathepsin B, D and L activity were measured in KO and WT neurons treated for up to 48 hours with PBS (negative control) or mouse WT PFFs at 70 nM using fluorescence-based assays (Abcam, UK) as per the manufacturer's instructions. Briefly, the primary neurons were lysed and incubated at 37°C for 2 hours protected from light with the cathepsin-B substrate sequence RR labelled with AFC (amino-4-trifluoromethyl coumarin) or with the cathepsin-D substrate sequence GKPILFFRLK(Dnp)-D-R-NH2, labelled with MCA (7-methoxycoumarin-4-acetic acid) or with the preferred cathepsin-L substrate sequence FR labelled with AFC. The released AFC or MCA was quantified using Tecan Infinite M200 Pro plate reader (Tecan, Maennedorf, Switzerland). The following wavelengths were used: for cathepsin B assay (Ex/Em = 400/505 nm), cathepsin D assay (Ex/Em = 328/460 nm) and cathepsin L (Ex/Em = 400/505 nm).

***AEP activity assay in primary neurons***

AEP activity was measured in KO and WT neurons treated for up to 21 days with PBS (negative control) or mouse WT PFFs at a final concentration of 70 nM, as previously reported^8^. Briefly, the primary cell cultures were lysed in the assay buffer (20 mM citric acid, 60 mM Na_2_HPO_4_, 1 mM EDTA, 0.1% CHAPS and 1 mM DTT, pH 6.0), which contains 20 μM AEP substrate Z-Ala-Ala-Asn coupled to the AMC (7-Amino-4-methylcoumarin) fluorescent dye (Bachem, Switzerland) and incubated at 37°C for one hour. AMC released was measured using the Tecan Infinite M200 Pro plate reader (Tecan, Maennedorf, Switzerland) with respective excitation and an emission wavelength of 380 and 460 nm.

***Calpain 1 and calpain 2 activity assay in primary neurons***

Using calpain activity assay kit (Abcam, UK), calpain activity was measured in primary culture treated with Tris buffer (negative control) or with aSyn WT PFFs for 3 hours and up to 21 days. At each indicated time-points, 100 μl of the extracellular media was collected before harvesting the neurons. Collected extracellular media and neurons were then lysed in the extraction buffer provided in the kit and centrifuged at 4°C at 13 000 rpm for 5 min. Soluble fractions were collected and handled following the manufacturer's instructions. Pellets from the lysed neurons were resuspended in the extraction buffer provided in the kit and mild sonication for 10 sec at 20% amplitude (Sonic Vibra Cell, Blanc Labo, Switzerland) was performed to ensure the complete dispersion of the pellet. Calpain activity assay was then performed following the supplier's instructions. Fluorescein emission was quantified using Tecan Infinite M200 Pro plate reader (Tecan, Männedorf, Switzerland) with respective excitation and an emission wavelength of 400 and 505 nm.

***Calpain 1 cleavage of aSyn PFFs* in vitro**

Cleavage of aSyn PFFs was performed as described previously^9^. Briefly, 0.15 U of active calpain 1 recombinant protein (Abcam, UK) was added to aSyn PFFs diluted to a final concentration of 50 μM in the reaction buffer containing 40mM 4-(2-hydroxyethyl)-1-piperazineethanesulfonic acid (HEPES) at pH 7.5 and 5 mM dithiothreitol (DTT) at 37 °C. The reaction was initiated by the addition of CaCl_2_ (1 mM final). Samples were collected before the addition of CaCl_2_ (time 0) or 2, 5, 15 or 30 mins after calpain 1 activation. Cleavage of aSyn by calpain 1 was assessed by Coomassie staining after separation of the samples onto a 16.5% SDS-PAGE gels.

***Quantitative real-time RT-PCR and transcriptomic analyses***

Primary hippocampal neurons were treated with WT PFFs or PBS buffer for 7, 14, and 21 days. Total RNA was isolated using the RNeasy Mini Kit (Qiagen, Switzerland) according to the manufacturer's protocol. The concentration of each sample was measured using NanoDrop (NanoDrop Technologies, Wilmington, DE, USA), and the purity was confirmed using the ratios at 260/280 nm and 260/230 nm.

*Quantitative real-time RT-PCR*

2 µg of RNA was used to synthesize cDNA using the High-Capacity RNA-to-cDNA Kit (Life Technologies, USA) following the manufacturer's instructions. To quantify aSyn, calpain 1, calpain 2 mRNAs levels, we used the SYBR green PCR master mix (Pack Power SYBR Green PCR mix, Life Technologies, USA). qRT/RT-PCR assay was performed using the following primers synthesized by Microsynth (Balgach, Switzerland):

| **Gene** | **Forward sequence (5'-3’)** | **Reverse sequence (5'-3’)** |
| --- | --- | --- |
| **SNCA** | AATGTTGGAGGAGCAGTGGT | GGCATGTCTTCCAGGATTCC |
| **calpain 1** | GAAAGGACCCTGGAGTGACA | TCCGGTGTAAGGTTGCAGAT |
| **calpain 2** | GTTGGTGAAAGGACATGCGT | TCAGGTTGCAGATCTCCAGG |
| **β-actin** | TTGTGATGGACTCCGGAGAC | TGATGTCACGCACGATTTCC |
| **GAPDH** | AACGACCCCTTCATTGACCT | TGGAAGATGGTGATGGGCTT |

40 cycles of amplification were then performed in an ABI Prism 7900 (Applied Biosystem, Foster City, USA) using 384-wells plate, which simultaneously allowed the analysis of the genes of interest and the housekeeping genes (β-actin and GAPDH) that serve as references for the normalization step. For each independent experiment, triplicated wells were acquired per condition and each experiment was reproduced at least 3 times independently. To quantify the expression level of the genes of interest in the different conditions tested, the comparative 2^-ΔΔCT^ method was used where ΔΔCT = ΔCT(target gene)−ΔCT(reference gene) and ΔCT = CT(target gene)−CT(reference gene). Results were expressed as the fold change relative to control neurons (2^-ΔΔCT^). geNorm method (RRID:SCR_006763, <https://genorm.cmgg.be/>) was performed to assess the most stable reference gene that should be used to normalize the gene expression^10^.

*Temporal transcriptomic analyses*

Libraries for mRNA-seq were prepared according to manufacturer's instructions with the TruSeq stranded mRNA kit (Illumina, USA) starting from 300 ng of good-quality total RNAs (RNA quality scores >8.9 on the TapeStation 4200). Libraries were subsequently loaded at 1.44 pM on two High Output flow cells (Illumina, USA) and sequenced in a NextSeq 500 instrument (Illumina, USA) according to manufacturer instructions, yielding 75 nt paired-end reads.

RNA-seq reads were aligned to the mm10 mouse reference genome (GRCm38.p5) with [STAR](https://www.ncbi.nlm.nih.gov/pubmed/23104886) (version 2.5)^11^. An average of 52 million of uniquely mapped reads per sample was then assigned to genes ([gencode primary assembly v16](https://www.gencodegenes.org/mouse/release_M16.html)) using [HTSeq-count](https://htseq.readthedocs.io/en/release_0.10.0/count.html) (RRID:SCR_011867, version 0.9)^12^. The analyses was focused on protein-coding and antisense RNA genes only, representing a total of 23,057 studied genes. Lowly expressed genes (CPM < 5) and genes present in less than three samples were excluded prior to further processing. Normalization and differential expression analyses of the remaining 11,767 genes were done with the R package DEseq2 (SARTools, RRID:SCR_016533, version 1.2)^13^, using the internal DEseq and the results methods. Genes with an absolute log2 fold-change greater than 1 and an FDR less than 0.01 were considered as significantly differentially expressed. Table 2 [combined_DEseq2_FC1_FDR_0_01_PFatLeastOnce.xls] contains the final results. Tables 3-4 contain respectively analyses of mitochondrial and synaptic genes significantly enriched.

***Quantification of LDH release in primary neurons***

Using CytoTox 96® Non-Radioactive Cytotoxicity Assay (Promega, Switzerland), released lactic acid dehydrogenase (LDH) in culture supernatants was measured following the manufacturer's instructions. After a 30 min coupled enzymatic reaction, which results in the conversion of a tetrazolium salt (INT) into a red formazan product, the amount of color formed, that is proportional to the number of damaged cells, was measured using Tecan Infinite M200 Pro plate reader (Tecan, Maennedorf, Switzerland) at a wavelength of 490 nm.

**Supplemental information - Figures titles and Legends**

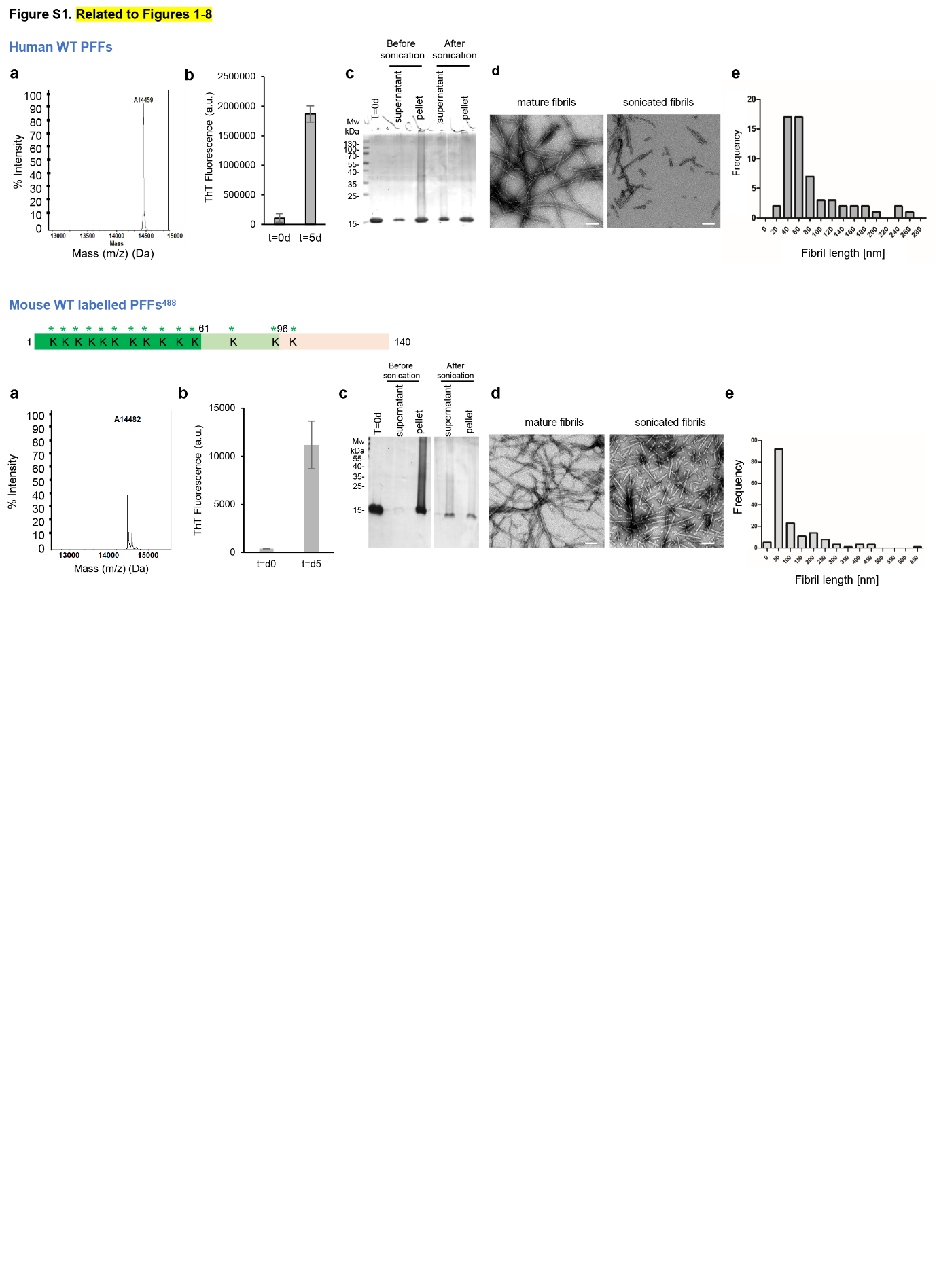
**Figure S1**

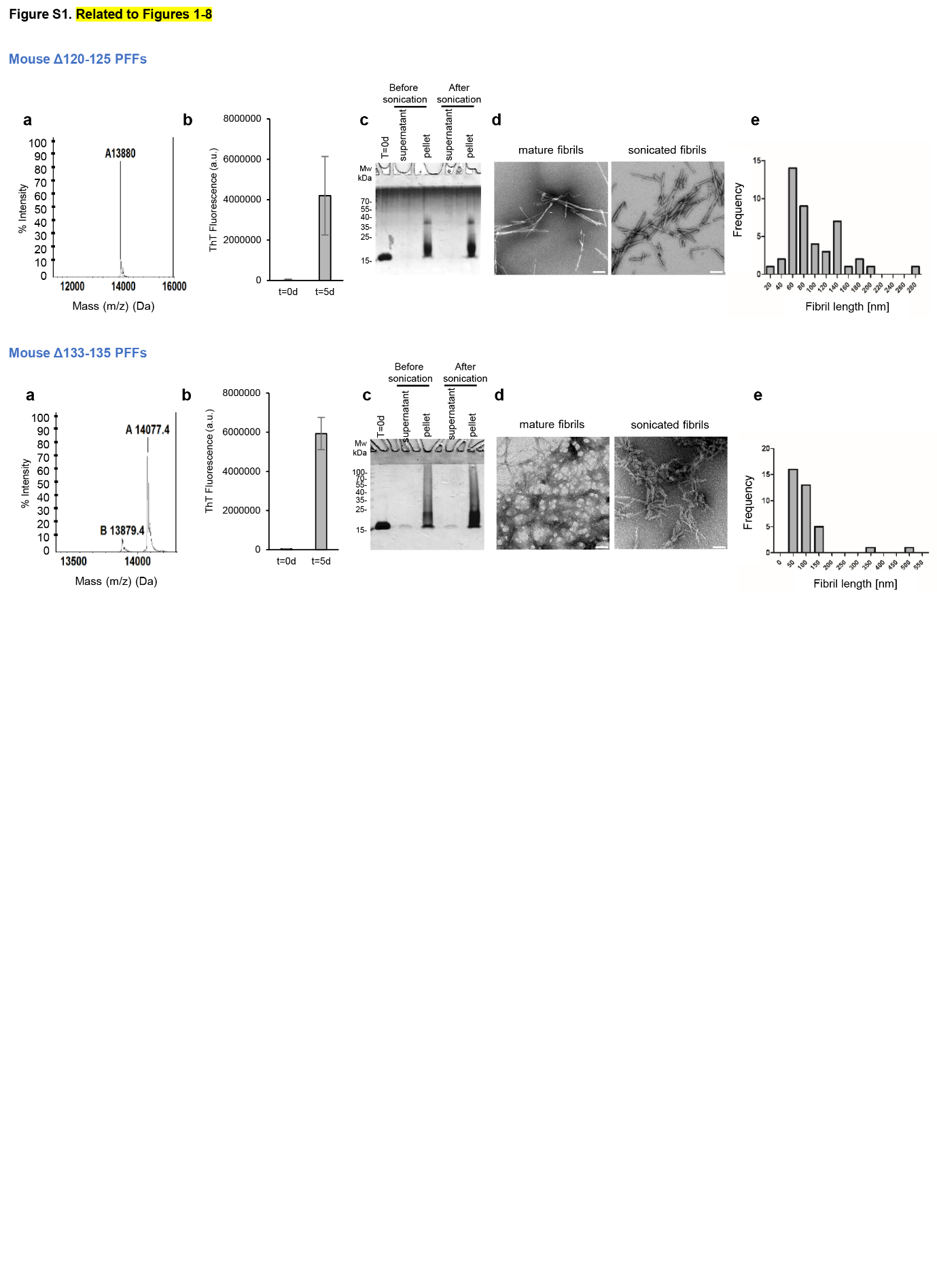

**Figure S1**

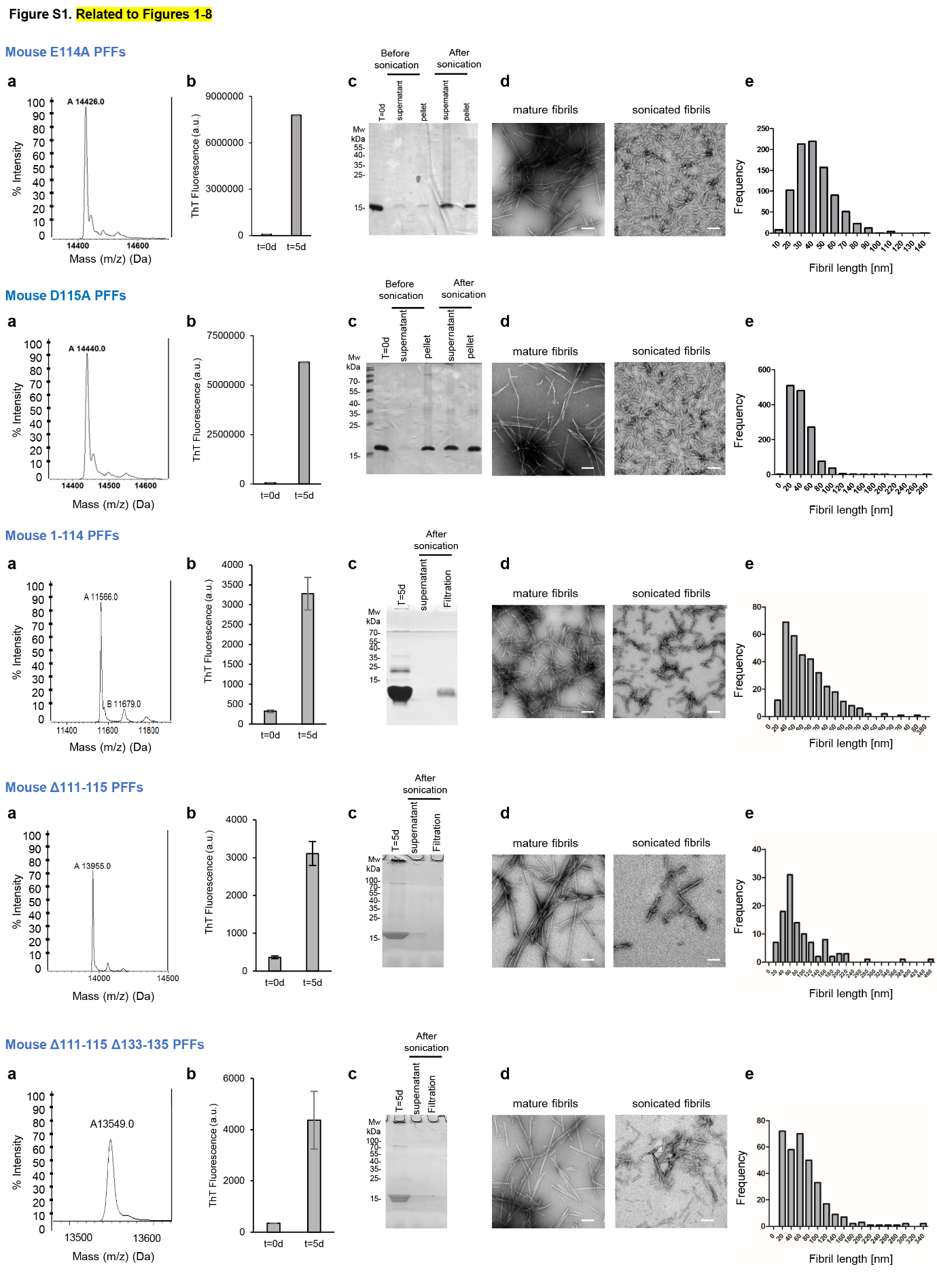

**Figure S1. Preparation and characterization of recombinant monomeric and preformed fibril (PFF) aSyn species (related to Figures 1 to 9). a.** Purity and characterization of aSyn monomers. Human aSyn and mouse aSyn mutants, including E114A, D115A, 1-114, Δ111-115, Δ120-125, Δ133-135 and Δ111-115-Δ133-135 were produced in *E. coli* and purified by anion exchange chromatography and size-exclusion chromatography, followed by a final chromatographic step using reverse-phase HPLC, as previously described^14^. The purity of recombinant monomeric aSyn after purification was assessed by ESI-LC/MS, which showed the expected mass. The recombinant mouse aSyn used in this publication has been previously described in Mahul-Mellier et al., 2020^15^. **b-e.** Purity and characterization of aSyn PFFs. aSyn PFFs were formed by incubation of monomeric aSyn for 5 days at 37°C under constant agitation at 1000 rpm. (**b**). After sonication, PFF formation was assessed by ThT fluorometry. All data represent the average ± SD (n=3). (**c**). The purity of aSyn PFFs was verified by SDS-PAGE gel and Coomassie blue staining. After sonication, PFF preparations were centrifuged, and the presence of the PFFs was verified in the pellet fraction, while the absence of monomer release after the sonication step was assessed in the supernatant fraction or after filtration through a 100 kDa filter (filtration). (**d-e**). aSyn PFFs were characterized by transmission electron microscopy (TEM) imaging. (**d**) Representative images of negatively stained aSyn PFFs before and after sonication. All aSyn PFFs showed the characteristic rigid non-branched fibrillar morphology. Scale bars = 100 nm. (**e**) The average length of the PFFs after sonication. Mouse aSyn PFFs used in this publication has been previously described in Mahul-Mellier et al., 2020^15^.

**Figure S2**

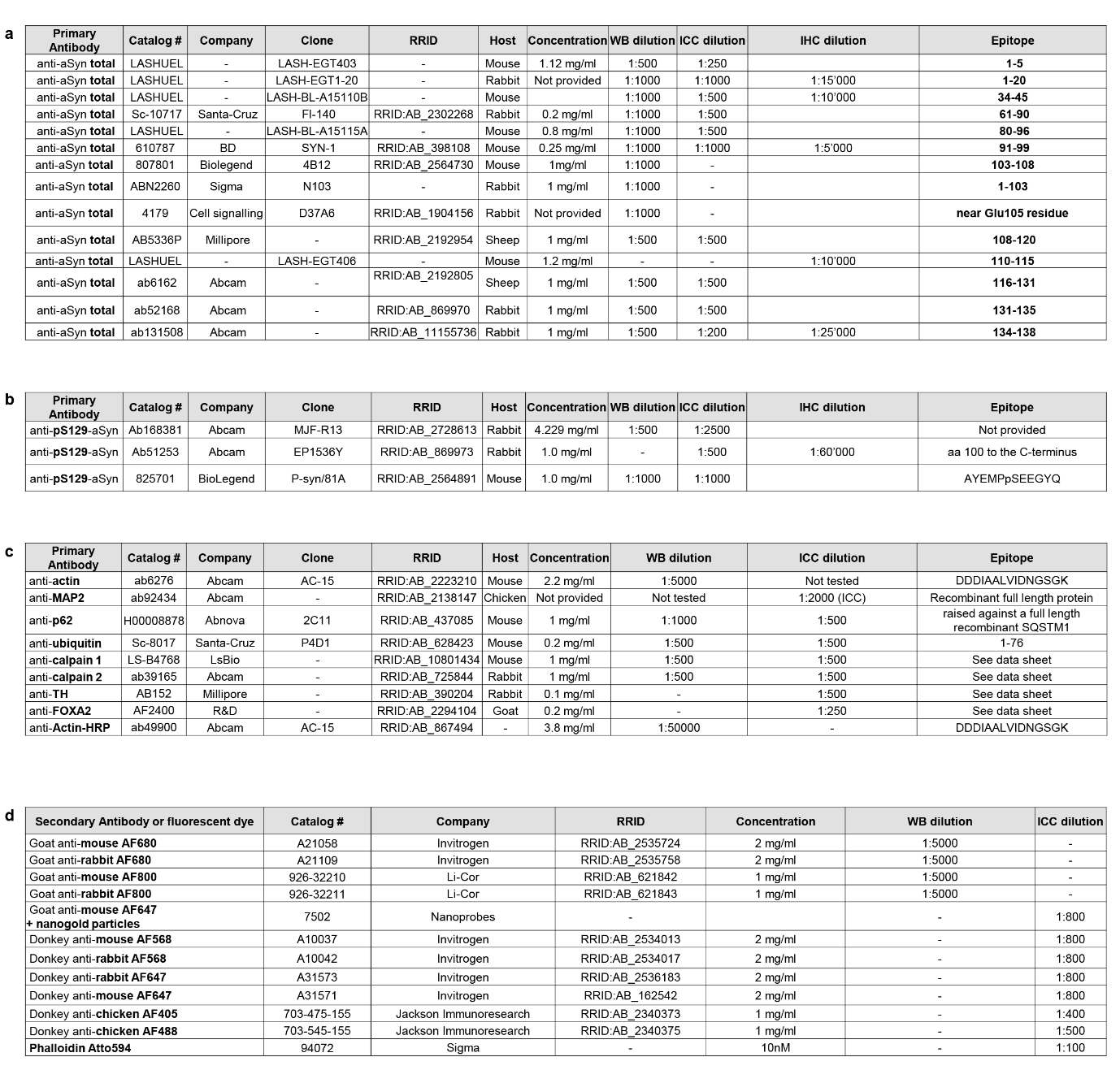

**Figure S2. List of the antibodies used in this study (related to Figures 1 to 10)**

**a**. Antibodies used for the detection of total aSyn.

For selected primary antibodies, immunohistochemical staining was performed as previously described in Figures 10 and S13b-c^16^^,^^17^.

**b**. Antibodies used for the detection of aSyn phosphorylated on S129 residue.

**c**. Other antibodies used in the study.

**d**. Secondary antibodies used for immunoblotting or confocal imaging.

**Figure S3**

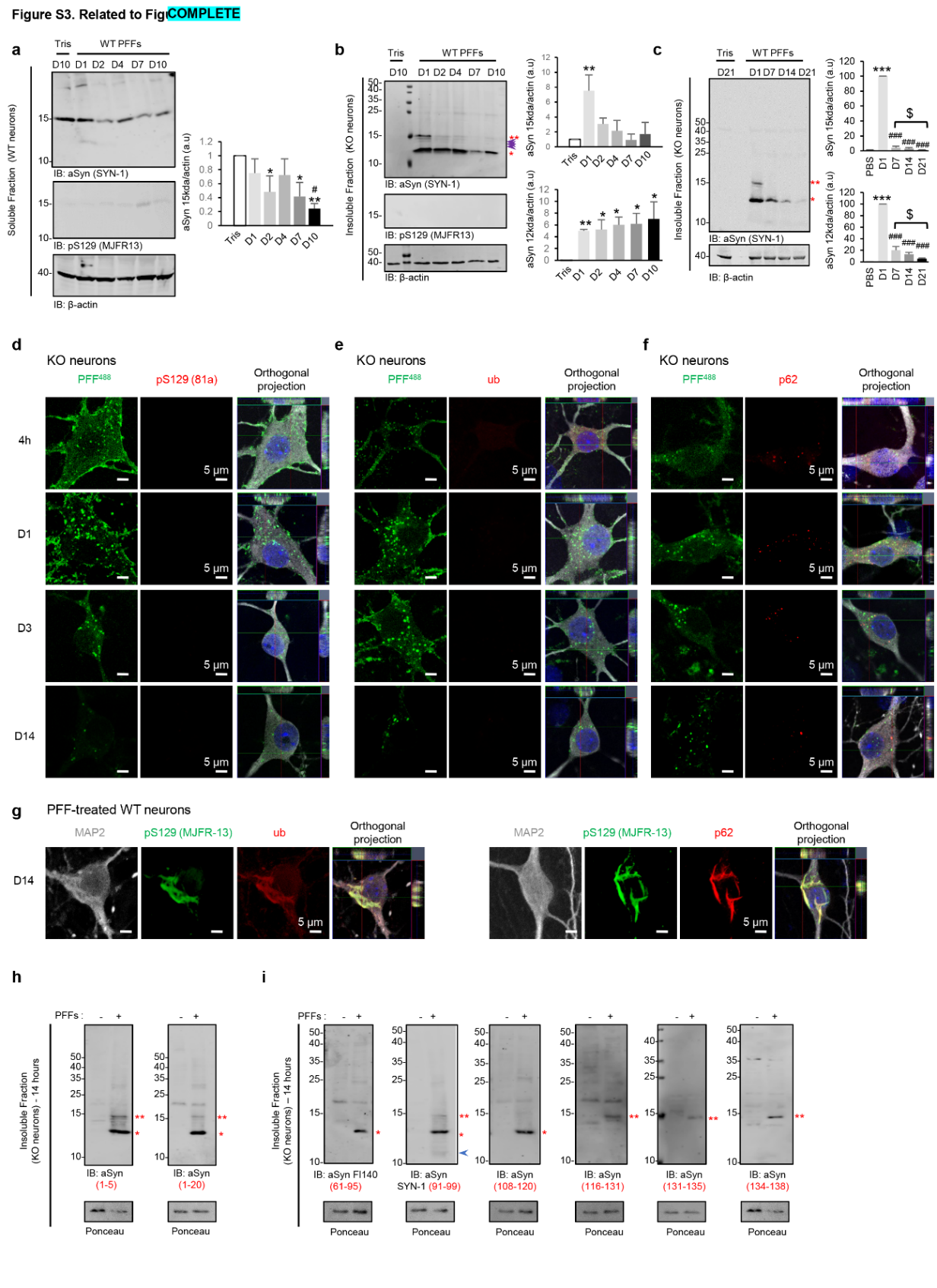

**Figure S3.** **Truncation of aSyn PFF seeds is an early event during the aSyn** **seeding process after internalization (related to Figure 1). a-c, h-i.** WB analyses of soluble fractions from PFF-treated WT neurons (**a**, related to **Figure 1f**) and insoluble fractions from PFF-treated aSyn KO neurons over time (b-c, h-i, related to Figure 1h-i). Control neurons were treated with Tris buffer. Cell lysates were immunoblotted after sequential extractions. Total aSyn, pS129, and actin were detected using SYN-1, pS129 (MJFR13), and actin antibodies, respectively. Levels of total aSyn (15 kDa, double red asterisk; 12 kDa, single red asterisk or HMW) or pS129-aSyn were normalized to actin. Purple arrows indicate intermediate truncated aSyn fragments below the full-length aSyn band. These fragments appeared within the first hour but did not increase over time, suggesting rapid cleavage to generate the 12 kDa fragment (see **Figures 1f, j**). In KO neurons, aSyn PFF seeds were fully cleaved after 24 hours. In WT neurons, intact aSyn at 15 kDa was reduced in the first four days but increased after day 7, coinciding with HMW bands and pS129 signal (**Figure 1f)**, indicating endogenous protein aggregation. Graphs represent the mean +/- SD of 3 independent experiments. a-c. *p<0.01, **p<0.001, ***p<0.0001 (ANOVA with Tukey HSD post-hoc test, Tris vs. PFF-treated neurons). a. #p<0.01 (ANOVA with Tukey HSD, PFF-treated D10 vs. D7, D4, or D1). c. ###p<0.0001 (ANOVA with Tukey HSD, PFF-treated D1 vs. D7, D14, or D21). c. $p<0.01, $$p<0.001 (ANOVA with Tukey HSD, PFF-treated D7 vs. D14 or D21). **d-g**. KO neurons were treated for up to 14 days with WT PFFs^488^ (d-f) and WT neurons with WT PFFs (**g**). Neurons were fixed and immunostained with pS129 (**d**, 81a; **g**, MJFR13), ubiquitin (**e, g** left panel), or p62 (**f, g** right panel) antibodies, counterstained with MAP2, and nuclei stained with DAPI. Scale bars = 5 μm.

**Figure S4**

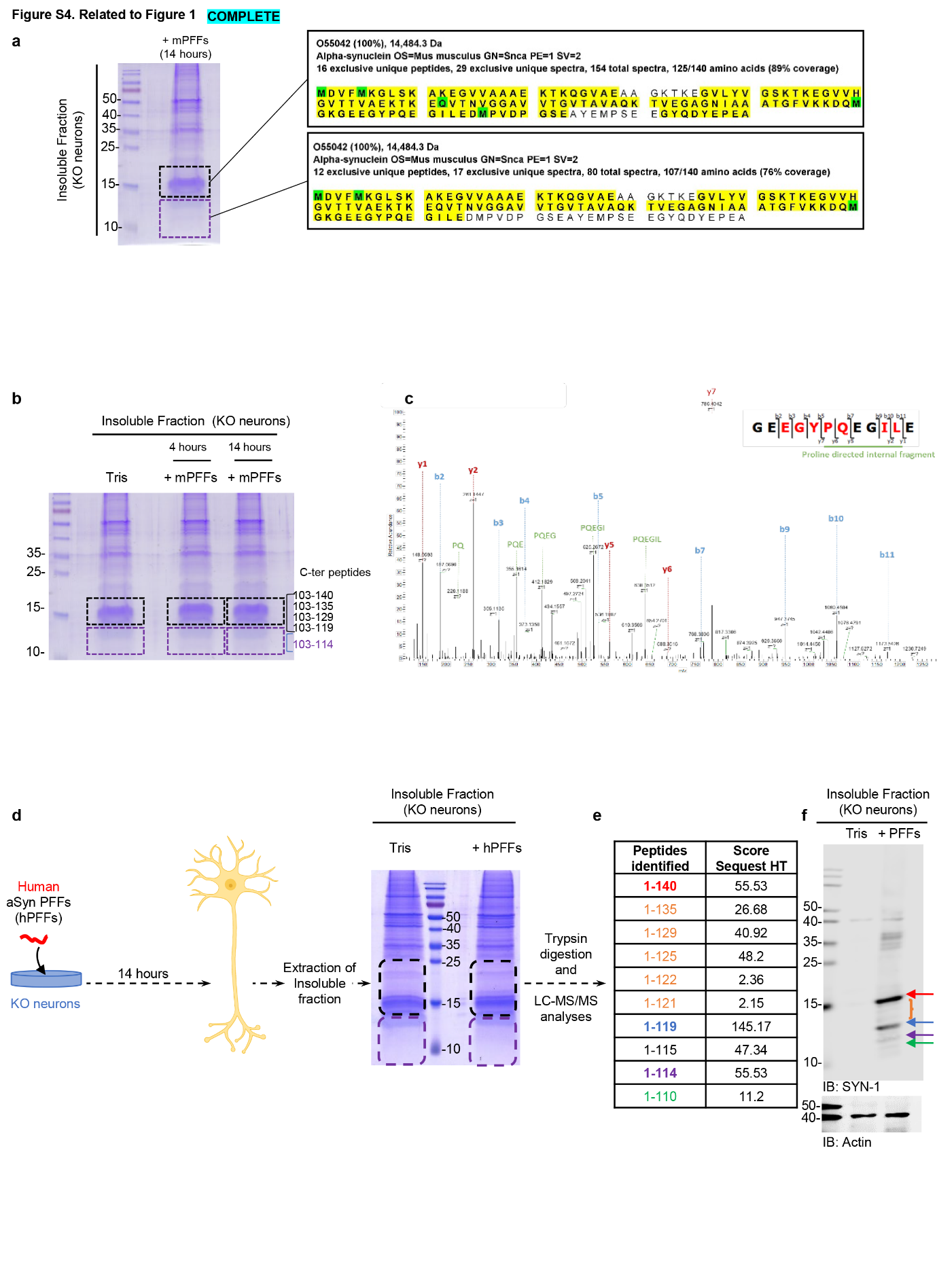

**Figure S4**

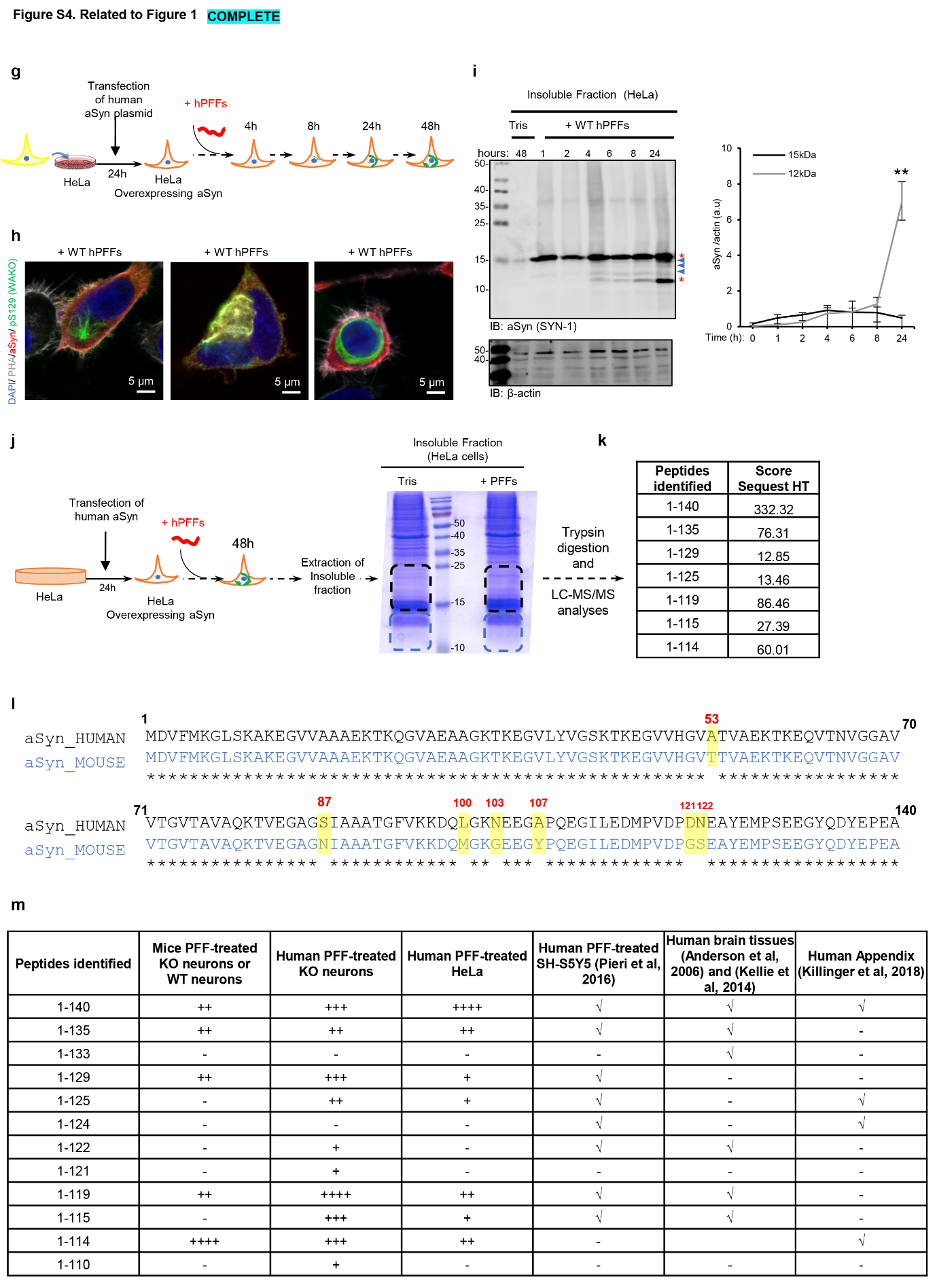

**Figure S4.** **C-terminal truncation of aSyn PFF seeds occurs primarily at residue 114 (related to Figures 1 and 3).** **a-k.** Insoluble fractions of aSyn KO primary neurons treated with 70 nM of mouse PFFs for 4 or 14 hours (**b-c**) or with 70 nM of human PFFs for 14 hours (**d-f**) were separated on a 16% Tricine gel. After Coomassie staining, two bands at ~15 (indicated by a black dashed box) and 12 kDa (indicated by a purple dashed box) were extracted from 16% Tricine gels (**a**, **b, d**). Isolated bands were selected based on the size of the proteolytic fragments observed by WB (**Figures 1j-k**) and subjected to proteolytic digestion followed by LC-MS/MS analyses. Glu-C was used to analyze N-terminal truncation^18^, which enables detection by LC-MS/MS of an intact peptide corresponding to the first 13 amino acids of aSyn (**a**). To analyze C-terminal truncations, aSyn was digested using trypsin, which cannot cleave the C-terminal domain, thus allowing detection of an intact peptide corresponding to the 103-140 region by LC-MS/MS^18^ (**c, e**). Although the N-terminal truncation has been reported previously for cell culture seeding models^19^^,^^20^, our proteomic analyses showed that the N-terminal domain of aSyn PFFs was intact in both the 15 and 12 kDa bands sliced from the tricine gel. These findings are consistent with our observation that antibodies raised against N-terminal residues 1-5 or 1-20 were still able to detect aSyn-cleaved products (~12 kDa) in the insoluble fraction of KO neurons **(Figure S3h)**. Our proteomic analyses also revealed that the C-terminal domain of mouse PFFs was cleaved at residues Asp-135, Ser-129, and Asp-119 (**b**) with a predominant site of cleavage at residue Glu-114 (**C**). This resulted in the formation of three fragments (1-135, 1-129, and 1-119) detected in the upper band excised in (**b**) and one main fragment (1-114) found in the lower band excised in (**b**). **d-f.** LC-MS/MS analyses revealed that internalized human PFFs in KO neurons were cleaved into 9 fragments, including the 1-114 fragment. Identified C-terminal fragments are listed in the table in **e**. **g-k.** Truncation of aSyn in HeLa cells overexpressing human aSyn. **g.** aSyn seeding model is based on the addition of human PFFs to HeLa cells overexpressing human aSyn. **h.** ICC analyses of LB-like inclusions that formed at 2 days after adding human PFFs (500 nM) to HeLa cells overexpressing human aSyn. After 48 hours of treatment, LB-like aSyn inclusions positively stained for pS129-aSyn were detected in the cytosol of HeLa cells by ICC. The panel shows examples of the aggregate heterogeneity detected using pS129 (MJF-R13 or WAKO) and total aSyn (SYN-1 or FL-140) antibodies. HeLa cells were stained with labelled Phalloidin^565^ (actin probe) and the nucleus was stained with DAPI staining. Scale bars = 5 μm. **i.** WB analyses of the soluble and insoluble fraction extracted from HeLa cells treated with Tris of human PFFs. As early as 4 hours after the delivery of the human PFFs, truncated aSyn fragments similar to those found in the neuronal seeding models were detected in the insoluble fraction of HeLa cells with a gradual increase in the amount of the main truncated form of aSyn (~12 kDa) over time. **j-k.** Proteomic analyses showed that human PFFs were also cleaved at the C-terminus after 14 hours in HeLa cells overexpressing aSyn. 6 fragments were detected, including the 1-114 fragment. Identified C-terminal fragments are listed in the table in **k**. The cleavage sites of the human PFFs were almost identical to those detected in the human PFF-treated KO neurons **(Figure S4e)** and in the human neuroblastoma cell line seeding model^19^ **(Figure S4m).** **i.** Amino acid alignment of human and mouse aSyn. The differences in amino acids are highlighted in yellow. **m.** Cleavage of mouse or human PFFs in cellular models and human tissues. The cleavage sites identified in mouse aSyn were close to those detected in the LBs from human brain tissue (Asp-115, Asp-119, Asn-122, Tyr-133, and Asp-135)^21^^,^^22^ or in an aSyn neuroblastoma cell line seeding model (Asp-115, Asp-119, Asn-122, Ala-124, Tyr-125, Ser-129, Tyr-133, and Tyr-135)^19^. This suggests the involvement of specific proteases in cellular responses to aSyn PFFs. Human and mouse aSyn proteins differ by only 7 amino acids, of which 5 are in the C-terminal region (L100M, N103G, A107Y, G121D, S122N, see **i**) and are in close proximity to the cleavage sites that we identified. This could explain the differences observed in the cleavage sites between PD models.

**Figure S5**

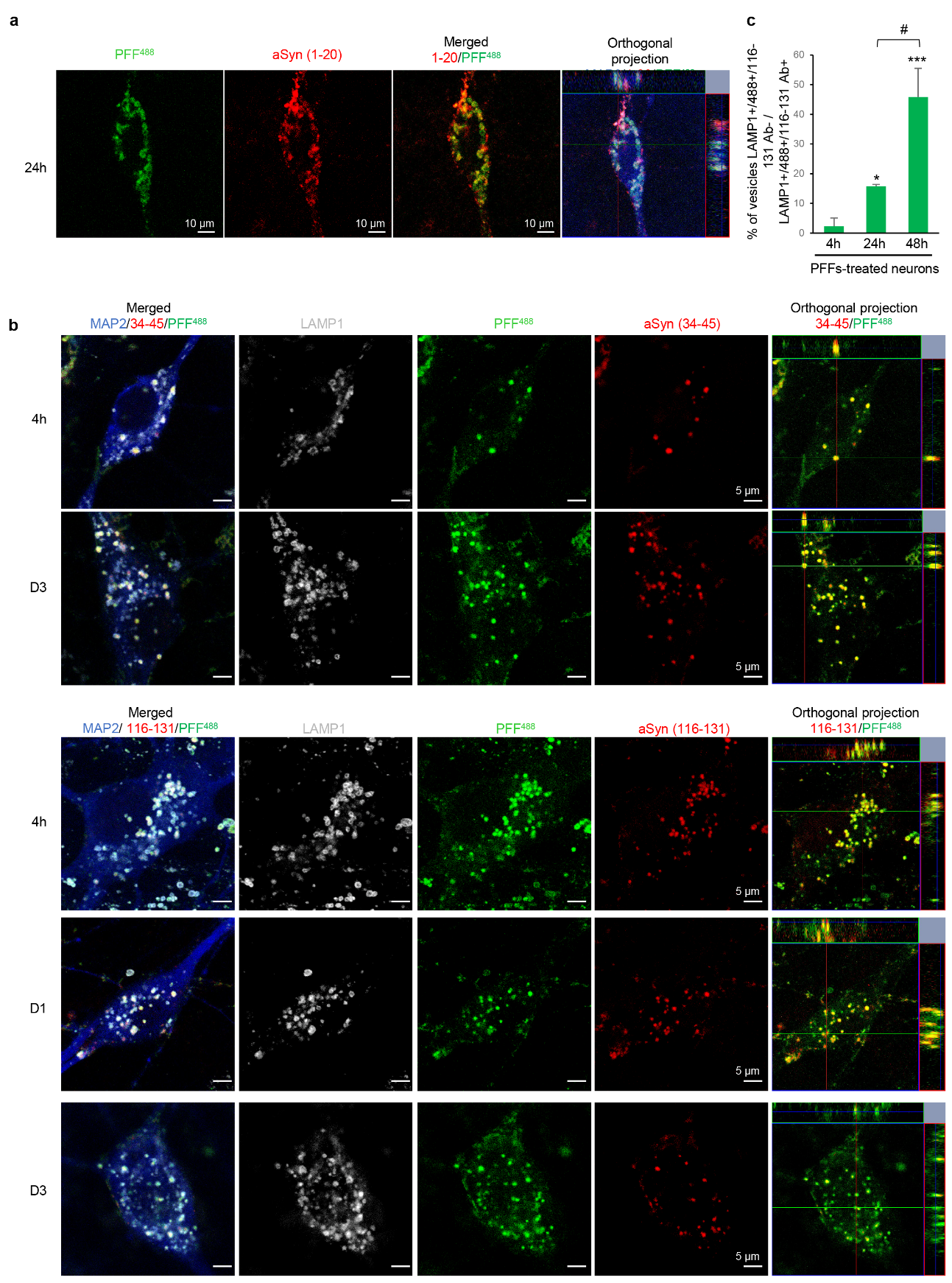

**Figure S5.** **aSyn PFF seeds are predominantly internalized via the endo-lysosomal route and are processed both in the endolysosomes and the cytosol of the neurons (related to Figure 2). a-b.** KO neurons were treated for up to 3 days with fluorescently labelled mouse WT PFFs^488^. **a.** Fluorescently labelled PFF^488^ seeds were detected by the N-terminal aSyn antibody (1-20) upon internalization. **b.** C-terminal truncation of the seeds in LAMP1-positive (late endosome, grey) compartments overtime was confirmed by the loss of detection of the seeds (green) by the C-terminal aSyn antibody (116-131, red) while N-terminal antibody (34-45, red) was still able to detect the seeds. **a-b.** Neurons were counterstained with MAP2 antibody and the nucleus with DAPI stain. Scale bars = 10 μm. **c.** C-terminal truncation of the seeds evidenced by the loss of detection of the PFFs (green) by the C-terminal aSyn antibody (116-131, red) in LAMP1-positive (late endosome, grey) compartments overtime was quantified by the counting of the number of LAMP1^+^/PFFs^+^/Ab 116-131^+^ objects vs. LAMP1^+^/PFFs^+^/Ab 116-131^-^ objects. An average of 180 vesicles were counted. The graphs represent the mean +/- SD of 3 independent experiments. p<0.01=*, p<0.001=** (ANOVA followed by Tukey HSD *post-hoc* test, 4h vs. other time-points). p<0.01=# (ANOVA followed by Tukey HSD *post-hoc* test, 24 h vs. 48 h).

**Figure S6**

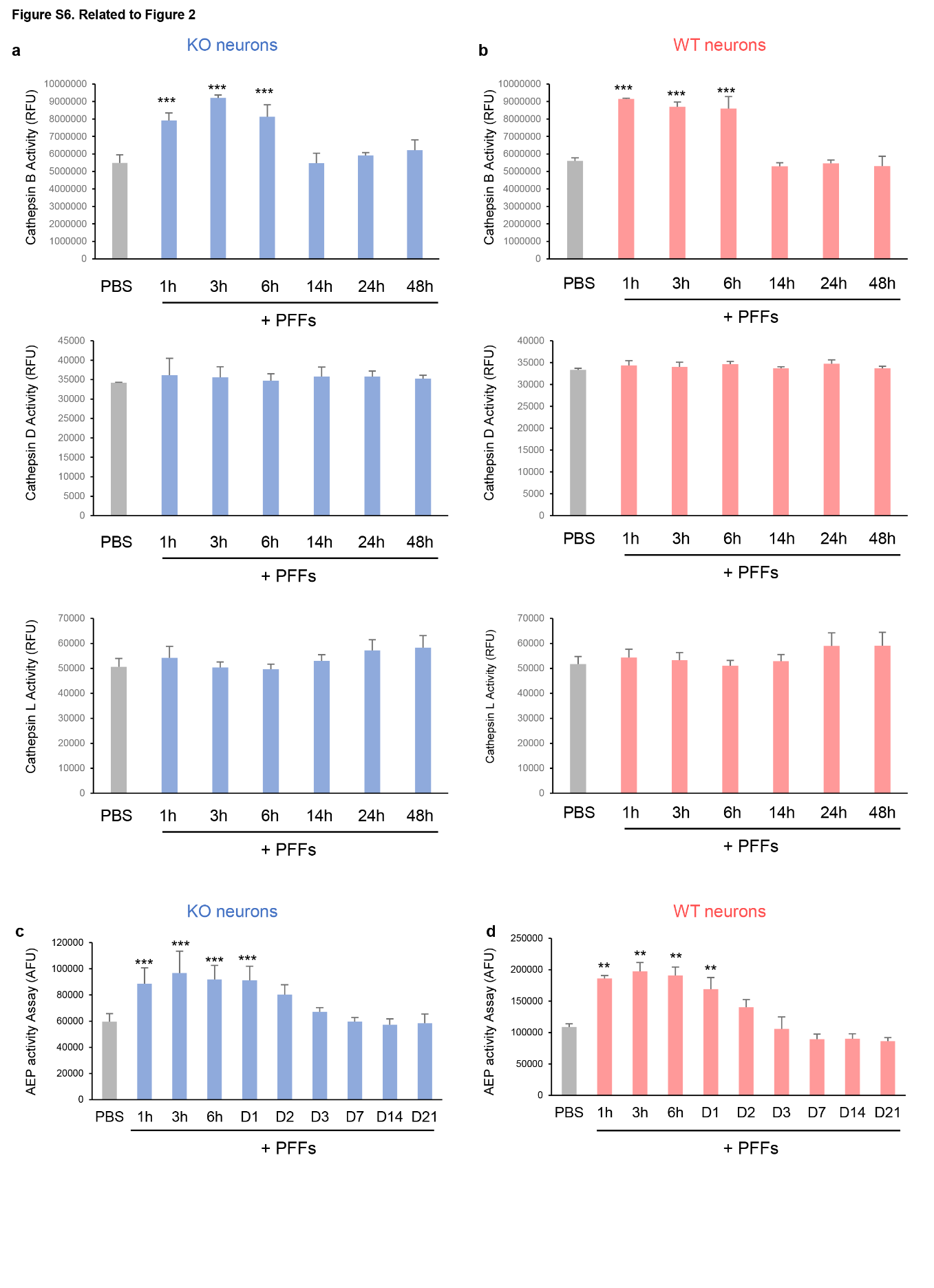

**Figure S6.** **Enzymatic activity of cathepsin B, D, L and AEP in PFF-treated WT and KO neurons overtime (related to Figure 2).**

**a-b.** Cathepsin B, D and L activity were measured in aSyn KO neurons (**a**) or WT neurons (**b**) treated with 70 nM of mouse WT PFF seeds up to 48 hours. Control neurons were treated with Tris buffer. The graphs represent the mean +/- SD of 3 independent experiments. p<0.0001=*** (ANOVA followed by Tukey HSD *post-hoc* test, Tris vs. PFF-treated neurons). Our data suggest that C-terminal truncation of the PFFs could occur in the endo-lysosomal compartments as they contain cathepsin enzymes that have been previously shown to be involved in the degradation of aSyn^23-26^ via proteolytic cleavages at multiple sites^27^. **c-d.** The AEP activity was measured in aSyn KO neurons (**c**) or WT neurons (**d**) treated with 70 nM of WT PFF seeds up to 21 days. Control neurons were treated with Tris buffer. The graphs represent the mean +/- SD of 3 independent experiments. p<0.001=**, p<0.0001=*** (ANOVA followed by Tukey HSD *post-hoc* test, Tris vs. PFF-treated neurons overtime).

**Figure S7**

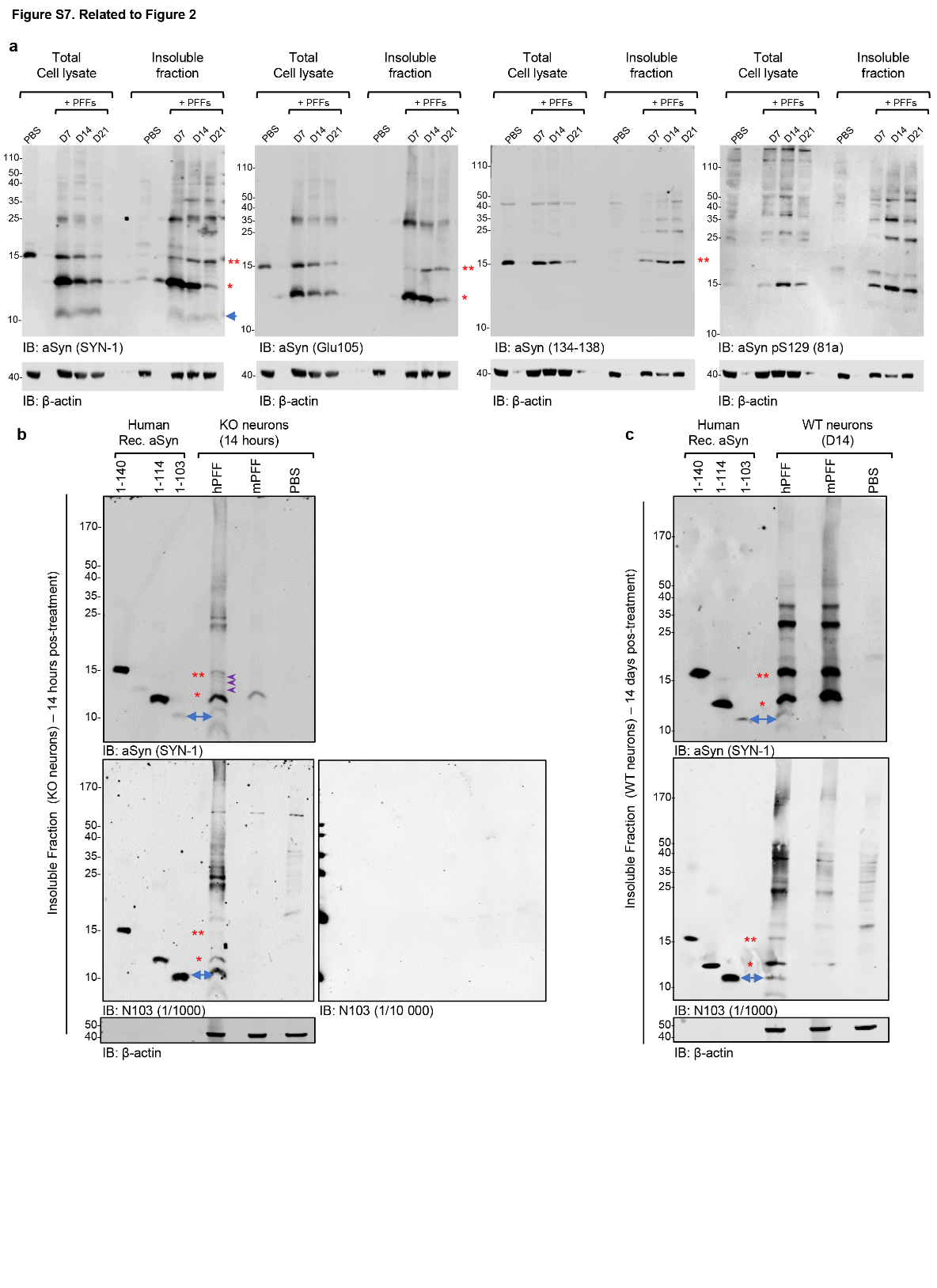

**Figure S7.** **Identification of the 10 kDa aSyn species formed in PFF-treated neurons (related to Figure 2).**

**a.** WB analyses of the insoluble fractions of the PFF-treated WT neurons at D7, D14 and D21. Mouse aSyn PFFs were added at a final concentration of 70 nM. Control neurons were treated with PBS buffer. Total aSyn was detected by SYN-1 (epitope: 91-99), Glu105 (epitope: near 103 a.a) or the C-terminal (epitope: 134-138) antibodies. pS129 level was probed with MJFR13 antibody. The total loading level was assessed using the actin antibody. Full-length aSyn (15 kDa) is indicated by a double red asterisk, the C-terminally truncated species (12 kDa) by a single red asterisk. The purple arrows indicate the intermediate truncated aSyn fragments and the blue arrow the 10 kDa species. **b-c.** To further identify the smallest C-terminal fragment, insoluble fractions of WT and KO neurons treated with either human or mouse PFFs were probed with the N103 antibody^8^ recently shown as specific to the human 1-103 fragment. WB analyses of the insoluble fractions of the PFF-treated KO neurons (**b**) or WT neurons (**c**) treated respectively for 14 hours or 14 days (D14) with human or mouse aSyn PFFs. Control neurons were treated with PBS buffer. Mouse aSyn PFFs were added at a final concentration of 70 nM. Human aSyn PFFs that are less efficiently internalized by the KO neurons and induce less seeding in WT neurons were added at a final concentration of 250 nM in order to detect a signal at a similar intensity than in the mouse PFF-treated neurons. 40 ng of monomeric recombinant human aSyn [full length (1-140) or C-terminally truncated species (1-114 and 1-103)] were loaded onto the gel in (**b-c**) to validate the specificity of the N103 antibody toward the 1-103 fragment and assess the size of the truncated fragments accurately. In our hands, this antibody recognized both full-length and C-terminally truncated fragments (i.e: 1-114 and 1-103). As advised by Zhang et al^8^, the immunoblot was first performed with the N103 antibody at a concentration of 1/10’000. Due to the absence of a signal at this concentration, we used the N103 antibody at 1/1000. At this dilution, the N103 antibody recognizes both the full-length and the 1-114 and 1-103 C-terminal truncated recombinant proteins. It also recognizes in the neuronal seeding model the full length (15 kDa indicated by the double red asterisk), the 12 kDa species indicated by the single red asterisk and, as expected, the 1-103 fragment (blue arrow).

**Figure S8**

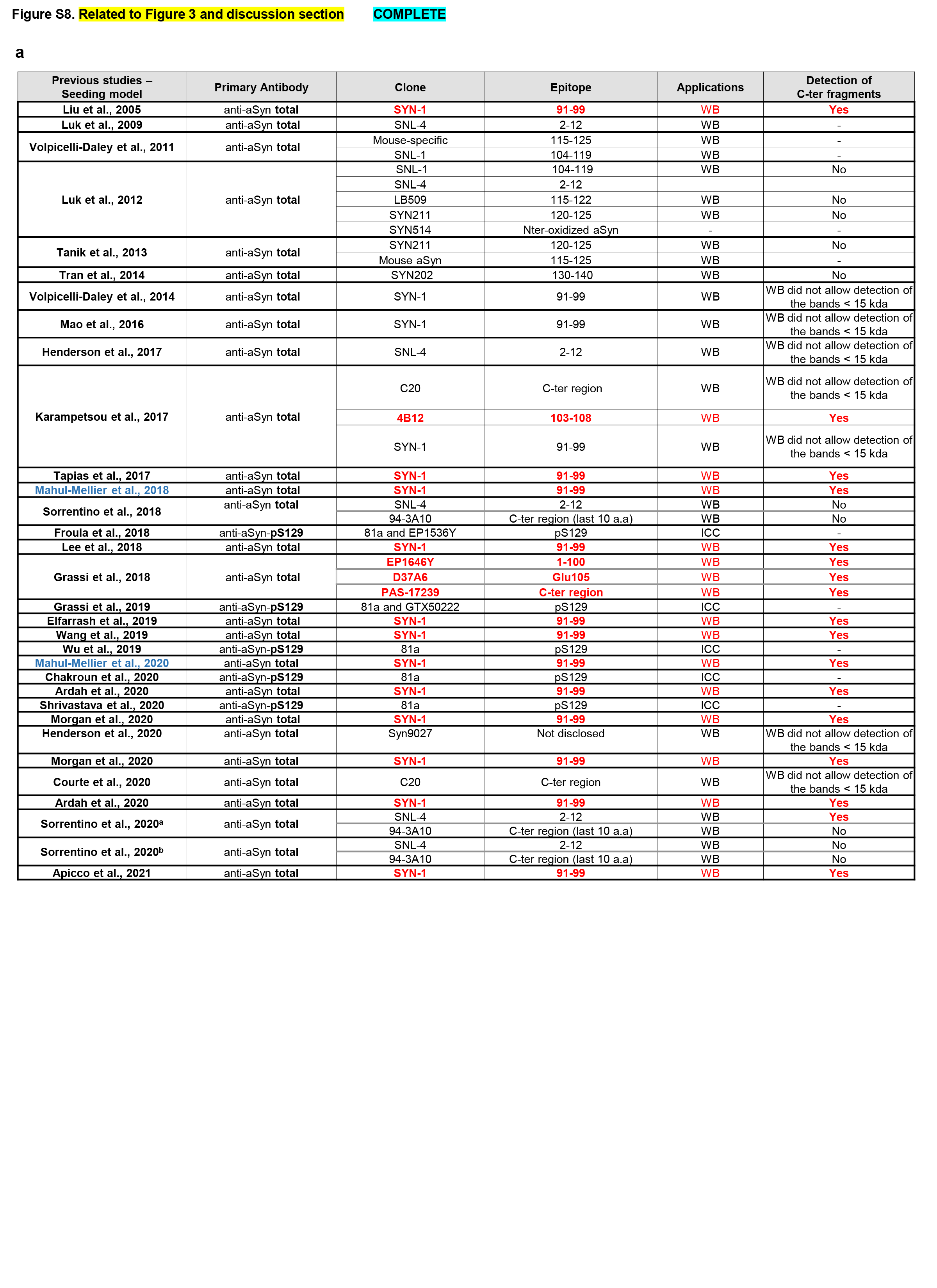

**Figure S8**

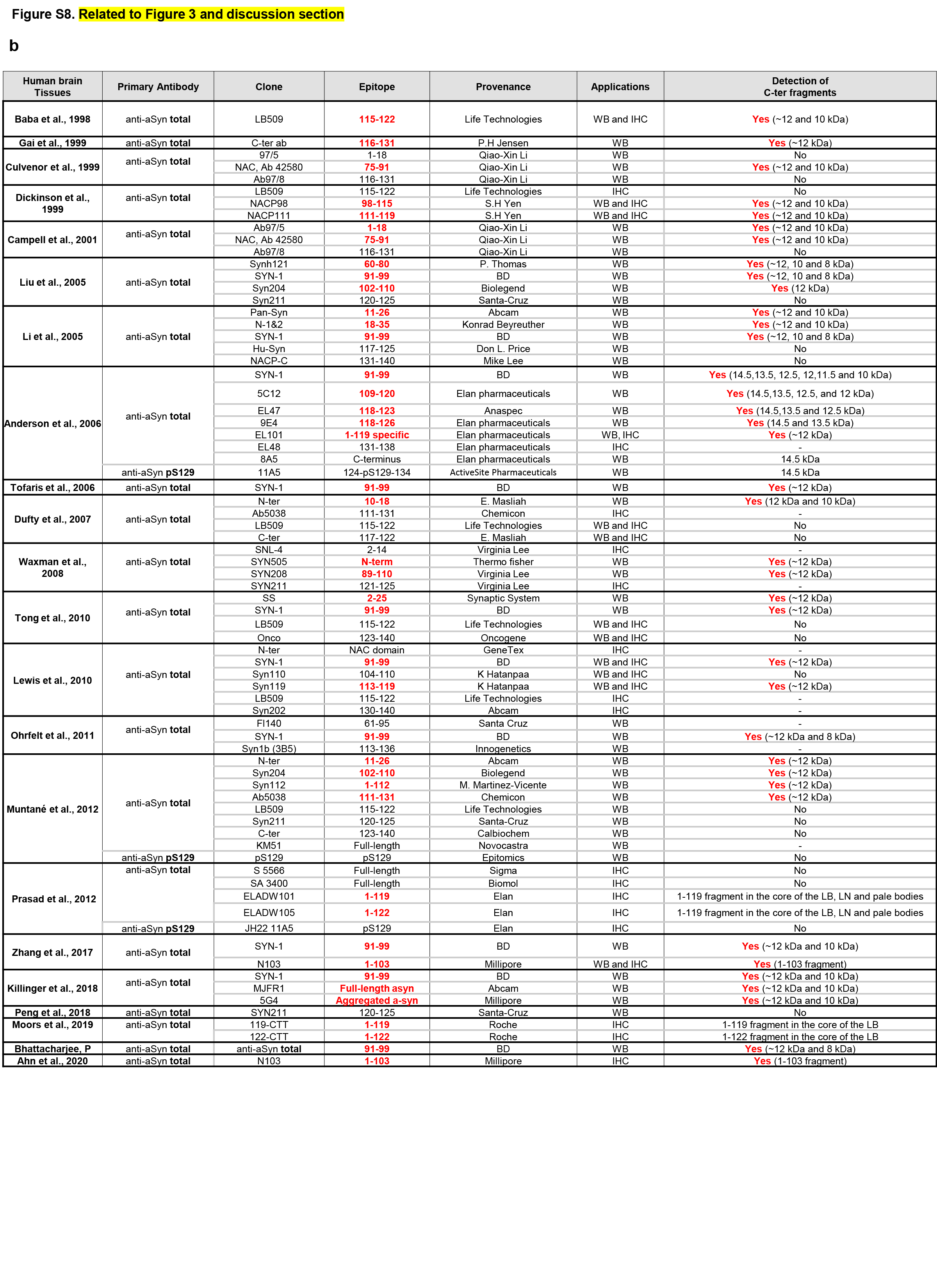

**Figure S8. Detection of total aSyn and truncated aSyn species in studies based on the neuronal seeding model or human synucleinopathy brain tissues (related to Figure 3). a**. Table showing the antibodies used to detect total aSyn by WB approaches in studies using the seeding model in primary neuronal cultures. **b**. Table showing the antibodies used to detect total aSyn in LB inclusions from human synucleinopathy brain tissues by WB or IHC approaches.

**Figure S9**

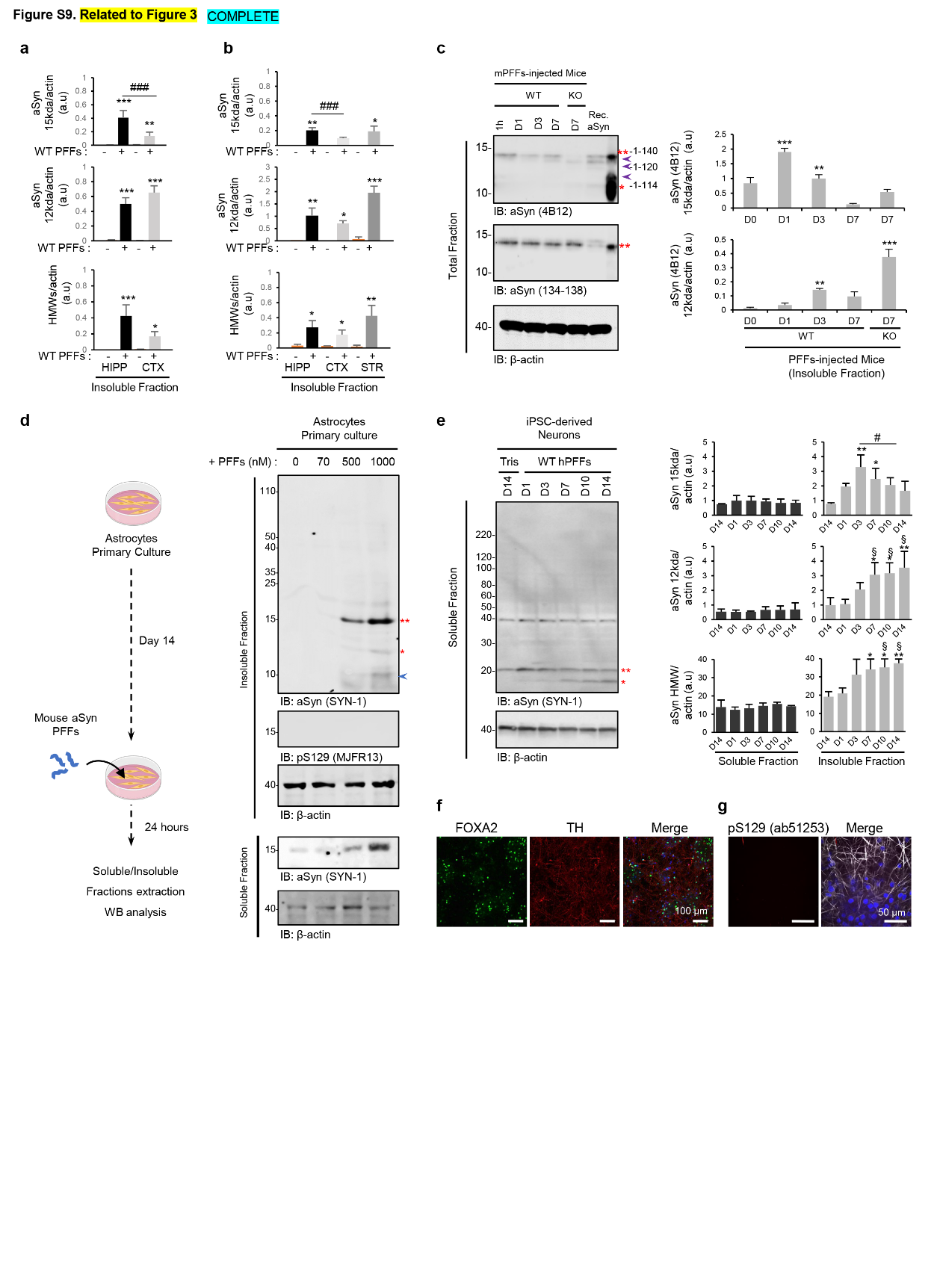

**Figure S9. C-terminal truncation is a general phenomenon in aSyn seeding and inclusion formation in cells (related to Figure 3).** Investigation of the truncation pattern of aSyn in rodents’ primary neurons (**a-b**), in mouse cortical astrocyte cultures (**c**), *in vivo* after injection of human PFFs in the striatum of WT and KO mice (**d**), in iPSC-derived neurons from a healthy control individual transduced with human PFFs (**e-g**). **a-b.** Quantification of aSyn levels in the insoluble fractions of hippocampal (HIPP) and cortical (CTX) primary neurons from mice (**a**) or hippocampal (HIPP), cortical (CTX), or striatal (STR) primary neurons from rats (**b**) treated for 10 days with 70 nM of mouse PFFs. The levels of aSyn were normalized to the relative protein levels of actin (**Figure 3 a-b**). The graphs represent the mean +/- SD of 3 independent experiments. *p<0.01, **p<0.001, ***p<0.0001 (ANOVA followed by Tukey HSD *post-hoc* test, Tris vs. PFF-treated neurons). ^###^p<0.0001 (ANOVA followed by Tukey HSD *post-hoc* test, hippocampal vs. cortical neurons). **c.** Insoluble and soluble fractions extracted from mouse cortical astrocyte cultures treated for 24 hours with increasing concentrations of PFFs (70 to 1000 nm). Control neurons were treated with Tris buffer (-). **d.** WB analyses of the total cell fraction extracted from the striatum from WT or aSyn KO mice dissected after 1 hour or 1, 3, or 7 days after injection with human PFFs. The levels of total aSyn (4B12 or 134-138 antibodies) (15 kDa, indicated by a double red asterisk; 12 kDa indicated by a single red asterisk or HMW) were normalized to the relative protein levels of actin. Purple arrows indicate the intermediate truncated aSyn fragments. The graphs represent the mean +/- SD of 3 independent experiments. **p<0.001, ***p<0.0001 (ANOVA followed by Tukey HSD *post-hoc* test, Tris vs. PFF-treated neurons). **e-g.** Truncation of aSyn in human iPSC-derived dopaminergic neurons. **e.** WB analyses of the soluble fraction extracted from iPSC-derived neurons were treated with 70 nM of human PFFs for 1, 3, 7, 10, and 14 days. The levels aSyn were normalized to the relative protein levels of actin. Purple arrows indicate the intermediate truncated aSyn fragments. The graphs represent the mean +/- SD of 3 independent experiments. **e.** p<0.01=§ (ANOVA followed by Tukey HSD *post-hoc* test, D1 vs. D7, D10 or D14, iPSC-derived neurons). p<0.01=# (ANOVA followed by Tukey HSD *post-hoc* test, D3 vs. D14, iPSC-derived neurons). **f-g.** Expression of neuronal midbrain markers in iPSC-derived neuronal cultures using an established methodology^4-6^. iPSC-derived neuronal cultures from healthy control were fixed and stained for proteins expressed in human midbrain dopaminergic neurons. Tyrosine hydroxylase (TH) and FOXA2 are expressed at DIV47 of neuronal differentiation. Scale bars = 100 μm. **g.** Phosphorylation of aggregated aSyn is a rare event in iPSC-derived neurons and the levels were not measurable by WB. ICC analyses of the inclusions formed at 16 days after adding human PFFs (70 nM) to iPSC-derived neurons. The panel shows an example of a phosphorylated aggregate detected using a pS129 (ab51253) antibody and counterstained with MAP2 and DAPI. Scale bars = 50 μm.

**Figure S10**

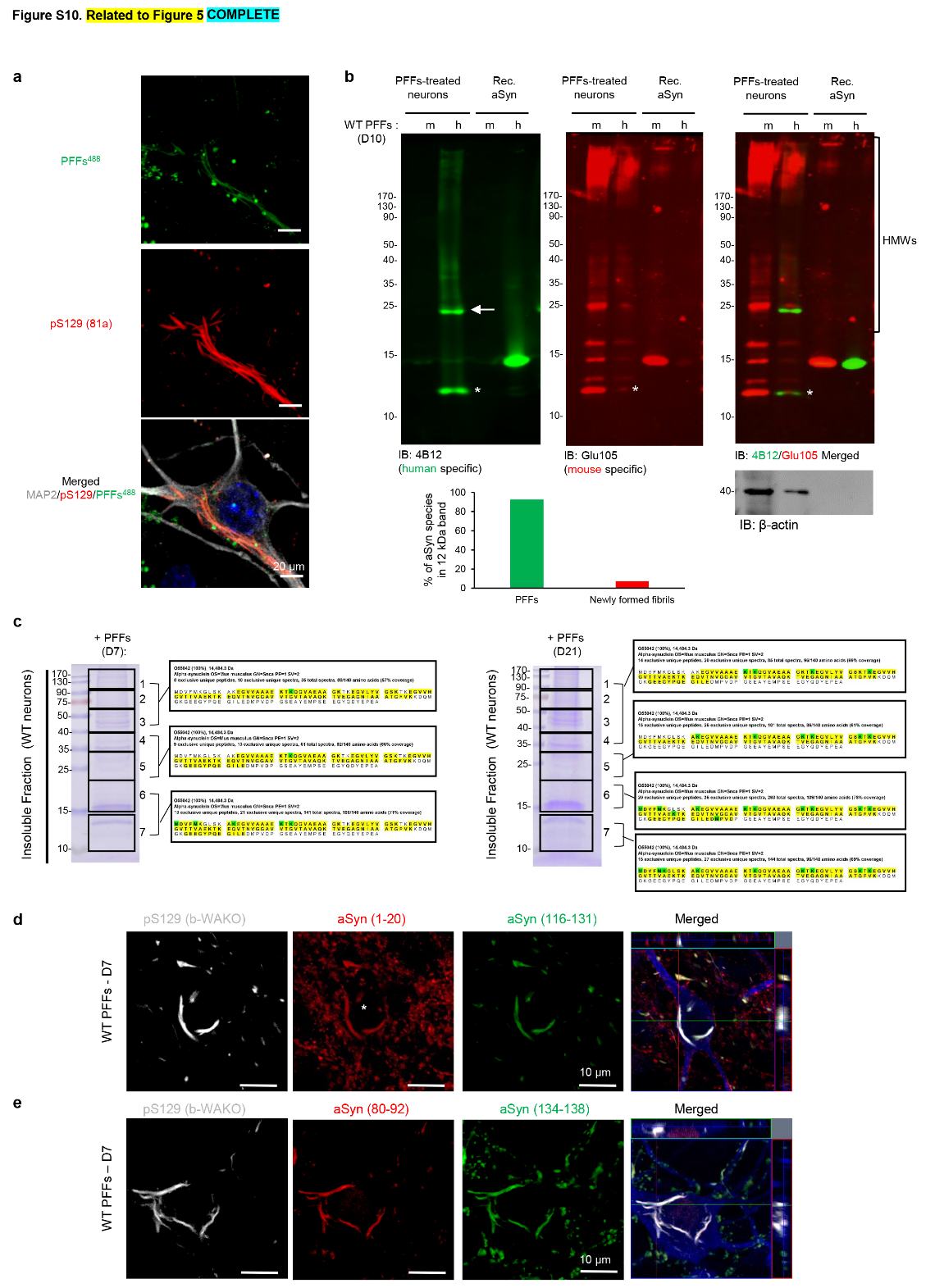

**Figure S10. aSyn fragments produced in neurons transduced with PFFs result from C-terminal truncation (related to Figure 5). a.** WT neurons were treated for 10 days with fluorescently labelled mouse WT PFFs^488^. Newly formed inclusions were detected using pS129 antibody (81a) and imaged by confocal imaging. Neurons were counterstained with MAP2 antibody, and the nucleus with DAPI staining. Scale bars = 20 μm. **b.** WB analyses of the insoluble fraction of WT neurons treated for 10 days with human or mouse WT PFFs. To discriminate the PFF seeds from the newly formed fibrils by WB analyses, mouse WT neurons were treated with human PFFs for 10 days, and the insoluble fraction was analysed by WB using a combination of a human-specific antibody (4B12) and a mouse-specific antibody (Glu105). Although the PFFs derived from human aSyn seeds were less efficient than mouse aSyn PFFs in mouse neurons^28^, we still observed seeding and the formation of inclusions. As expected, the human-specific aSyn antibody (4B12) showed that the truncated fragment at ~12 kDa detected after 10 days of treatment was composed of the initial PFF seeds added to the neurons. In contrast, the newly formed aggregates (HMW bands) were mainly detected by the mouse antibody (Glu105), reflecting the incorporation and aggregation of the endogenous mouse aSyn protein. Additionally, it was interesting to note that endogenous mouse aSyn was cleaved upon the human PFF-seeding condition. **c.** Proteomic analyses shows that the N-terminal domain of aSyn was intact in the insoluble fraction of the WT neurons treated for 7 and 21 days with WT PFFs. **d-e.** aSyn WT neurons were treated for 7 days with WT PFFs. Newly formed inclusions were detected using pS129 antibody (81a clone) in combination with N-terminal (**d**, epitope 1-20) or NAC (**e**, epitope 80-92) antibodies together with C-terminal epitopes [116-131] (**d**) or [134-138] (**e**) antibodies. The newly formed aggregates were similarly detected by the pS129, N-terminal, NAC, or C-terminal antibodies. Scale bars = 20 μm.

**Figure S11**

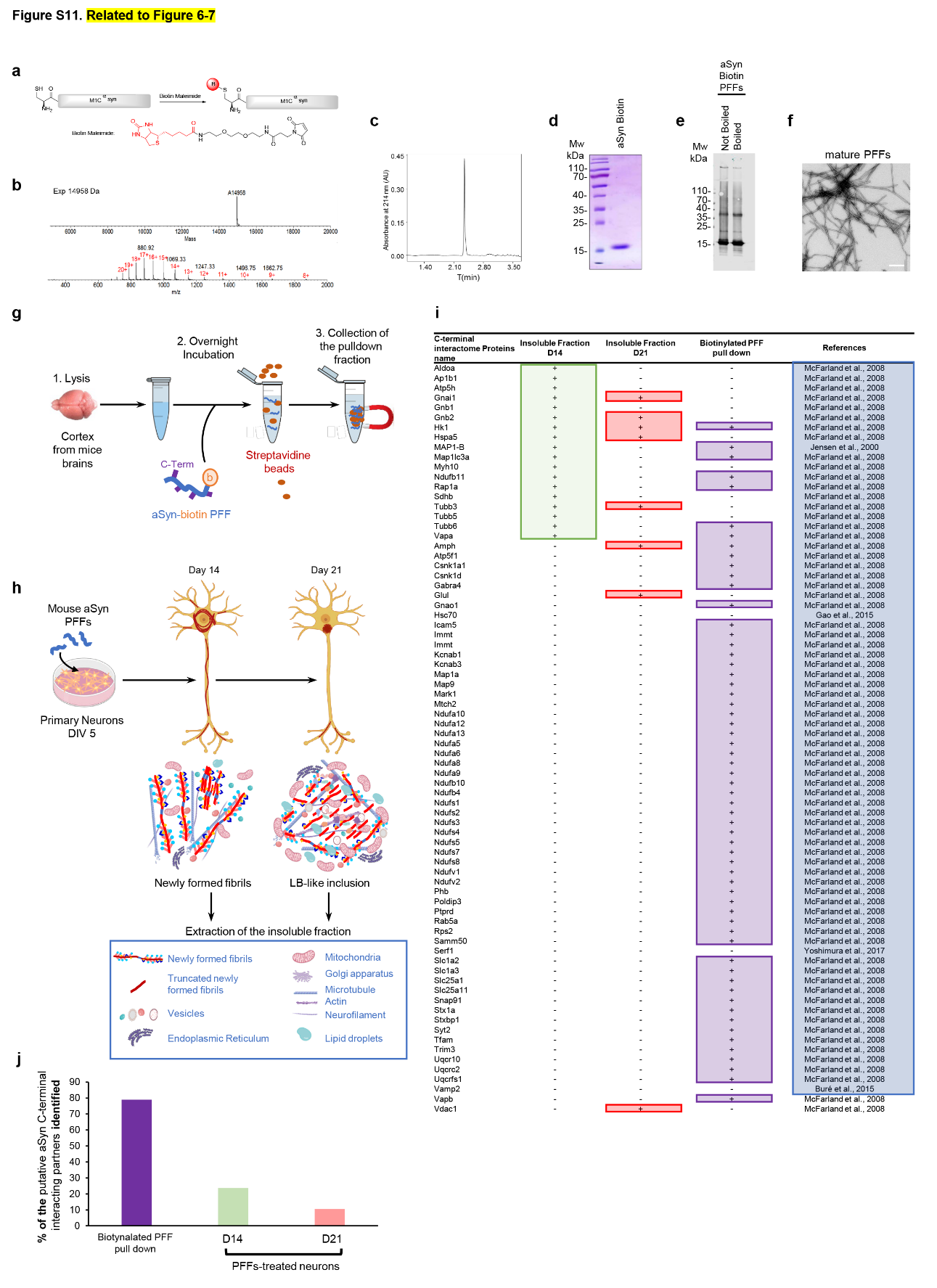

**Figure S11. C-terminal truncations alter newly formed aSyn fibrils and consequently disrupt interactions with the C-terminally interacting partners (related to Figure 5). a**. Strategy to add biotin to monomeric aSyn. **b**. ESI-LC/MS analyses of biotinylated M1C aSyn (observed mass: 14958 Da, expected mass: 14958 Da). **c**. Water acuity UPLC spectra of biotinylated M1C aSyn using a BEH 300 C4 column 2.1 mm × 100 mm, 1.7 μm with a linear gradient of 10−90% B over 4 min (solvent A: water/0.1% TFA, solvent B: acetonitrile/0.1% TFA). **d**. SDS-PAGE analyses of biotinylated M1C aSyn. **e**. WB of biotinylated M1C aSyn fibrils with and without boiling in sample buffer. **f**. TEM of biotinylated M1C aSyn fibrils. **g.** PFFs labelled with biotin were incubated with mouse cortical brain lysates, and biotinylated fibrils were specifically pulled-down using streptavidin beads. **h.** Primary neurons were treated for 14 days or 21 days with mouse WT PFFs, and the insoluble fraction was isolated. The proteome of the insoluble fractions from the pull-down fraction (**g**) and the PFF-treated neurons (**h**) was identified using LC-MS/MS analyses. Identified aSyn interactors were compared to the previously reported putative C-terminal interactors of aSyn^29^. **i-j.** The table in **i** depicts the presence of the putative C-terminal interactors of aSyn^29-34^ in the pull-down assay using full-length biotinylated WT PFFs (**g**) or in the insoluble fractions of PFF^WT^-treated neurons over time (**h**). **j.** Histogram shows the percentage of putative C-terminal interactors identified in the different pull-down assays.

**Figure S12**

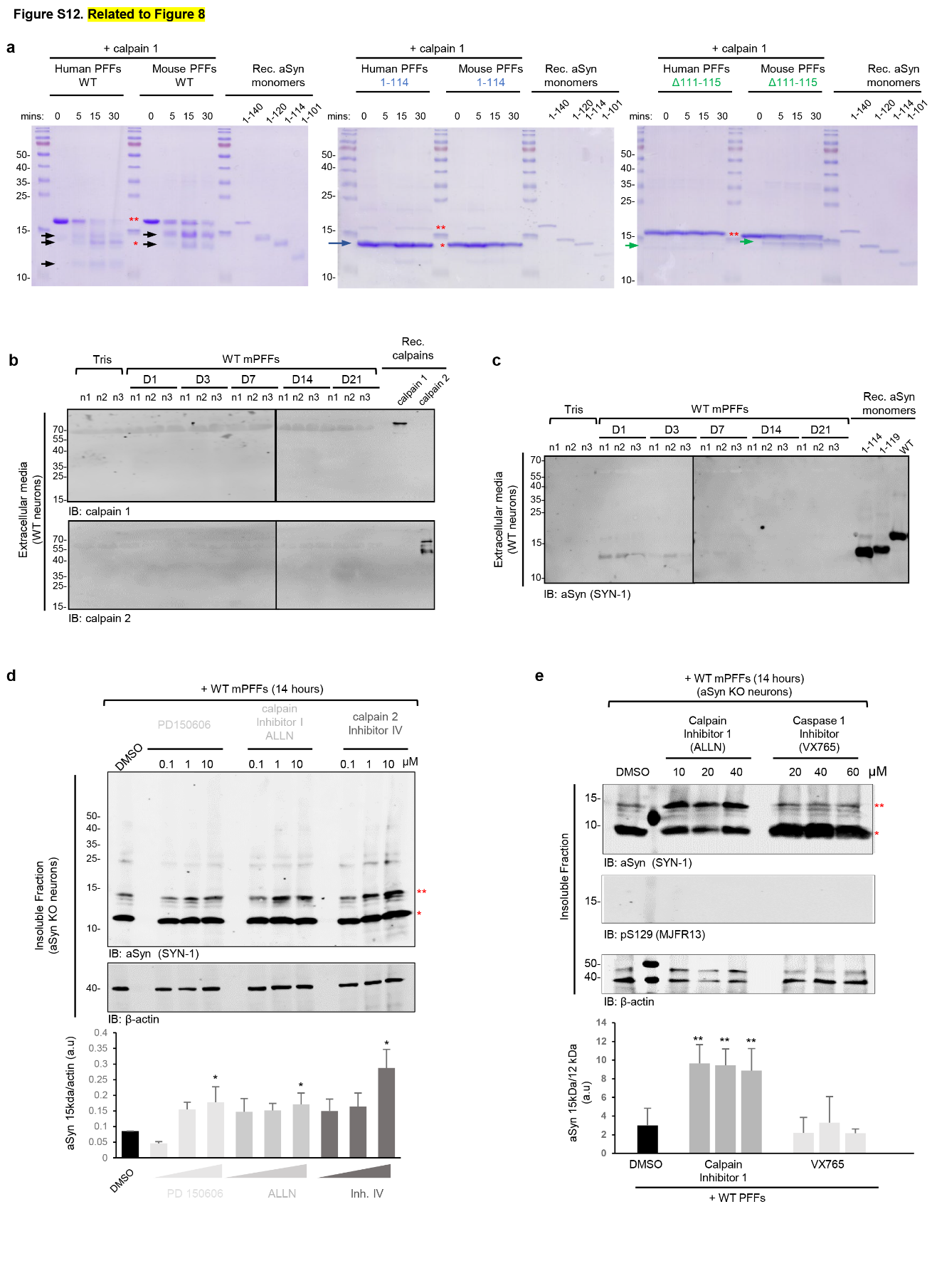

**Figure S12. Calpains 1 and 2 are involved in the truncation of aSyn *in vitro* and primary neurons (related to Figures 7-9). a.** *In vitro* calpain assays. Human and mouse aSyn PFFs^WT^, PFFs^1-114^ and PFFs^Δ111-115^ were incubated with active recombinant calpain 1 for the indicated times. Residues 111-115 are required for calpain cleavage. **b-c.** Immunoblotting analyses show the protein levels over time of calpain 1 (a top panel), calpain 2 (a bottom panel), and aSyn (**b**) in the extracellular media of PFF-treated WT neurons or in control neurons (Tris). **d-e.** aSyn KO neurons were pre-treated for 6 hours with increasing concentrations of calpain 2 inhibitor (10, 20, or 40 µM), calpain 1 inhibitor PD105606 (10, 20, or 40 µM), or DMSO as a control (**d**) or with the caspase 1 inhibitor (VX765) (20, 40, or 60 µM) (**e**). 70 nM of PFFs were then added for 14 hours. Cell lysates were analysed by immunoblotting after sequential extractions of the soluble and insoluble fractions. Total aSyn was detected by SYN-1 antibody. The histogram shows the densitometry analyses from 3 independent experiments. The graphs represent the mean +/- SD of 3 independent experiments. p<0.001=*, p<0.001=** (ANOVA followed by Tukey HSD *post-hoc* test, DMSO vs. enzymatic inhibitors in PFF-treated KO neurons).

**Figure S13**

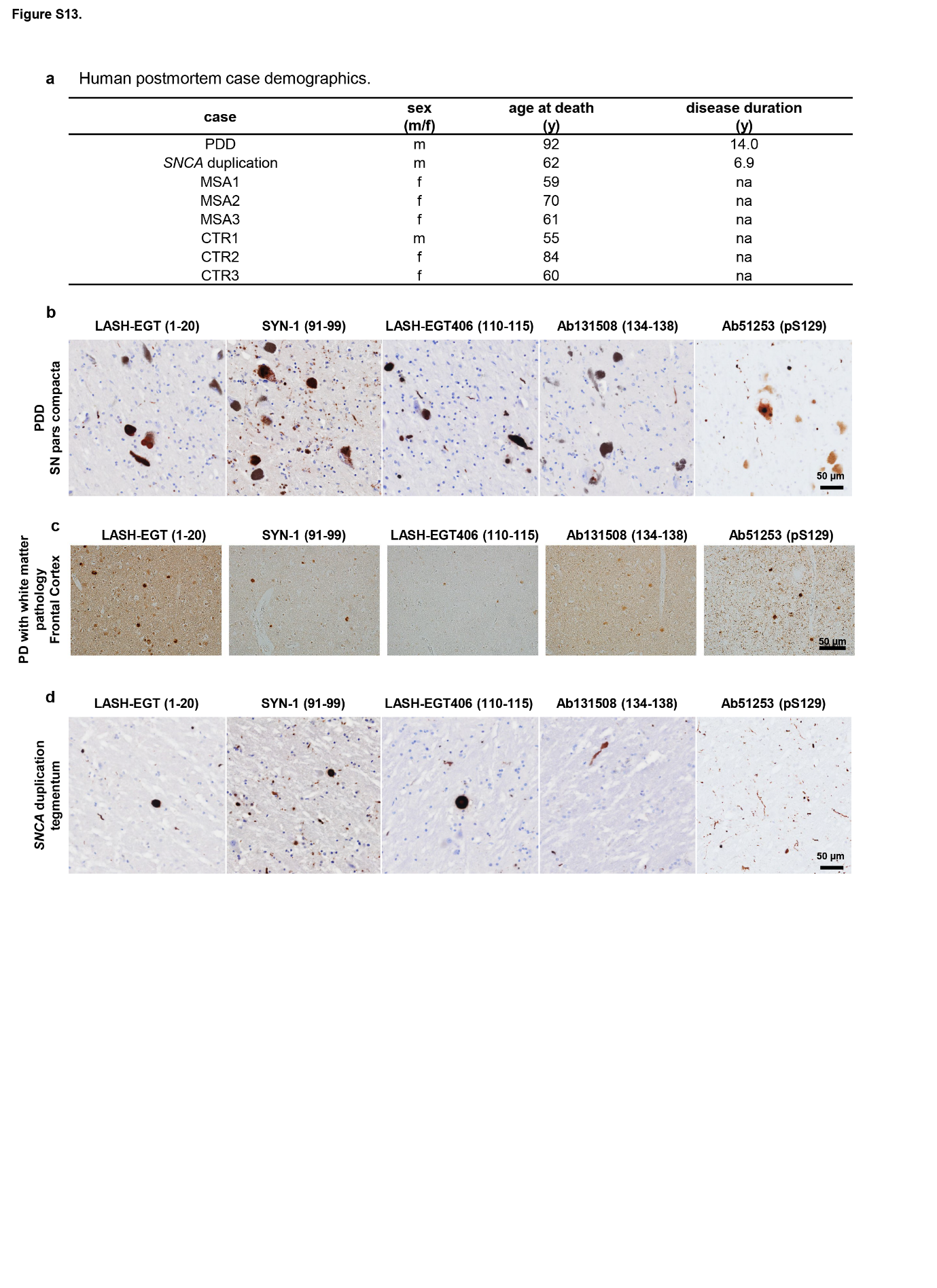

**Figure S13. The use of a set of aSyn** **antibodies to capture the morphological spectrum of aSyn pathology in MSA and sporadic PD brain tissues (related to Figure 10)**

**a.** Human postmortem case demographics for **Figure 10.** **b-d.** Serial sections from the midbrains of PDD (**b**, pars compacta), frontal cortex from PD with white matter pathology (**c**) and *SNCA* duplication (**d**, tegmentum) cases were stained with aSyn antibodies raised specifically against the N-terminal (epitope: 1-20), the NAC (91-99), the C-terminal (epitopes: 110-115 and 134-138) or pS129 (EP1536Y) regions. Lower magnification. Scale bars = 50 μm.

**Figure S14**

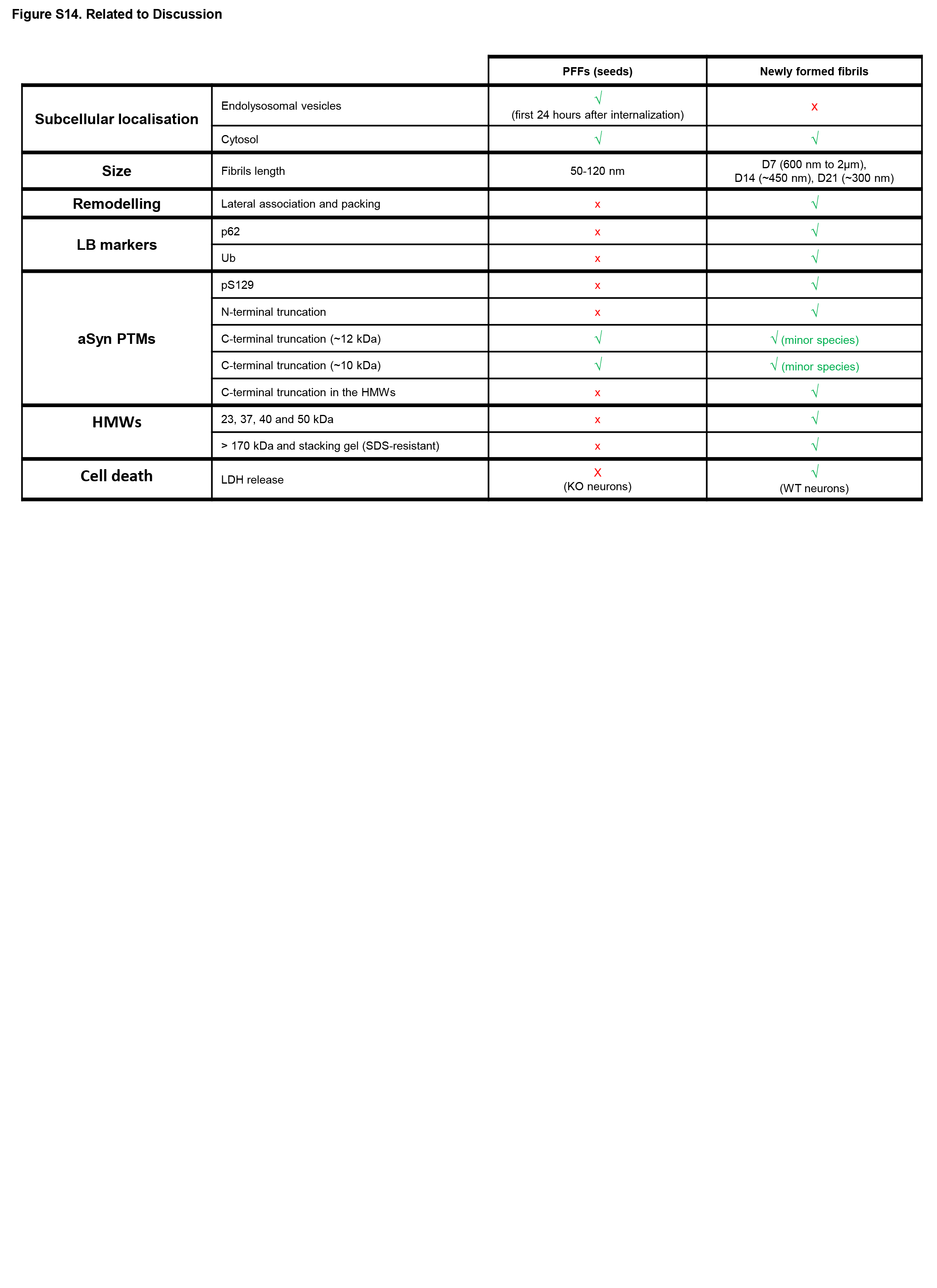

**Figure S14. Specific features of the PFF seeds vs the newly formed fibrils in the neuronal seeding model** **(related to discussion).**

**Figure S15**

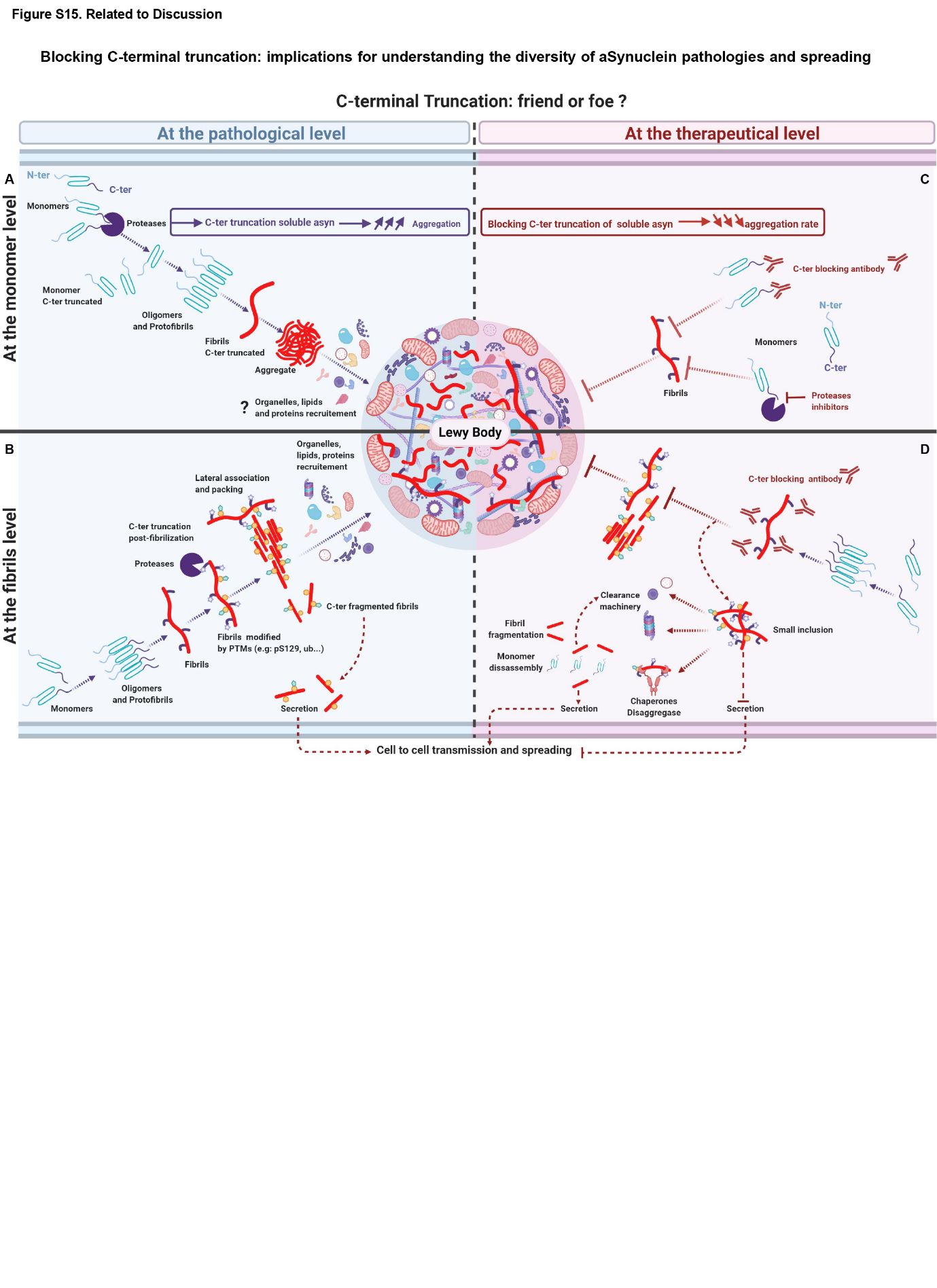

**Figure S15. C-terminal truncation: Friends or foe?** **(related to discussion)**

Our study demonstrates that the C-terminal cleavage of aSyn is one of the primary triggers for initiating aSyn aggregation *in vivo* and packing the fibrils into higher-order inclusions (**a-b**). Consistent with our working hypothesis, blocking or reducing aSyn C-terminal cleavage at the monomeric level (**c**) or at the fibrillar level (**d**) using monoclonal antibodies directed against the C-terminal part of aSyn or using inhibitors of proteases that cleave within the C-terminus of aSyn could protect against aSyn-induced neurodegeneration and could attenuate aSyn pathology spreading in human and animal models of PD and related synucleinopathies. Together, these findings suggest that inhibiting the C-terminal truncation represents a viable therapeutic strategy for the treatment of PD and synucleinopathies.
